# Epigenetic plasticity cooperates with emergent cell-cell interactions to drive neoplastic tissue remodeling in the pancreas

**DOI:** 10.1101/2022.07.26.501417

**Authors:** Cassandra Burdziak, Direna Alonso-Curbelo, Thomas Walle, Francisco M. Barriga, José Reyes, Yubin Xie, Zhen Zhao, Chujun Julia Zhao, Hsuan-An Chen, Ojasvi Chaudhary, Ignas Masilionis, Zi-Ning Choo, Vianne Gao, Wei Luan, Alexandra Wuest, Yu-Jui Ho, Yuhong Wei, Daniela Quail, Richard Koche, Linas Mazutis, Tal Nawy, Ronan Chaligné, Scott W. Lowe, Dana Pe’er

## Abstract

The response to tumor-initiating inflammatory and genetic insults can vary amongst morphologically indistinguishable cells, suggesting yet uncharacterized roles for epigenetic plasticity during early neoplasia. To investigate the origins and impact of such plasticity, we perform single-cell analyses on normal, inflamed, pre-malignant and malignant tissues in autochthonous models of pancreatic cancer. We reproducibly identify heterogeneous cell-states that are primed for diverse late-emerging neoplastic fates and link these to chromatin remodeling at cell-cell communication loci. Using a new inference approach, we reveal signaling gene modules and tissue-level crosstalk, including a neoplasia-driving feedback loop between discrete epithelial and immune cell populations that we validate by genetic perturbation in mice. Our results uncover a neoplasia-specific tissue remodeling program that may be exploited for pancreas cancer interception.

**One-Sentence Summary:** Single-cell analysis reveals that enhanced epigenetic plasticity drives pro-neoplastic crosstalk in early pancreatic cancer.

## Main Text

The initial events by which tissues diverge from normalcy to form benign neoplasms and malignant tumors remain poorly understood. Genetic changes have long been known to drive this process (*1*); however, the discovery of prevalent cancer driver mutations in phenotypically normal epithelia (*2*) challenges the classic notion of cancer pathogenesis and underscores the essential role of cellular and environmental context (*3–5*). Indeed, non-mutagenic environmental insults promote tumor initiation in mice (*6, 7*) and chronic inflammatory conditions substantially increase cancer risk in humans (*8–10*). Interestingly, these events can have heterogeneous effects even amongst morphologically indistinguishable and genetically identical cells from the same tissue (*11*). Genetic tracing studies similarly reveal that all such cells are not equally prone to undergo neoplastic and malignant reprogramming (*12*). This heterogeneity suggests that for tumorigenesis to proceed, select mutant cells either possess or gain an enhanced ability to acquire novel cell states, a phenomenon known as cellular plasticity (*13, 14*).

Developmental, regenerative, and pathologic plasticity is largely determined at the chromatin level as increases or decreases in the repertoire of transcriptional programs that can be accessed by a given cell (*14, 15*). Cells showing a high degree of plasticity, such as stem cells, often have a more ‘open’ or accessible chromatin landscape that becomes restricted during differentiation (*16, 17*). Previous work has used de-differentiation with respect to normal cell-states to characterize cancer cell plasticity with single cell genomics from lung cancer models (*18, 19*). However, we still do not know how plasticity emerges in the earliest stages of tumorigenesis, particularly in concert with the environmental insults that accelerate these initiating events. Learning how plasticity is triggered to arise in pre-malignant tissues and how it contributes to early tumor evolution is paramount to understanding and intercepting cancer at its earliest stages.

Pancreatic ductal adenocarcinoma (PDAC) is a lethal cancer that is typically diagnosed too late for intervention, and is a prime example of a disease that arises from cooperativity between genetic and epigenetic reprogramming events (*20–22*). Unlike more genetically heterogeneous cancers, PDAC is almost universally initiated by an activating mutation in the proto-oncogene *KRAS.* However, *KRAS*-mutant epithelia can remain phenotypically normal and depend on inflammatory stimuli (pancreatitis) to transform into pre-neoplastic and neoplastic lesions (*7, 23, 24*). We (*25*) and others (*26*) have reported that oncogenic KRAS, in the absence of further mutation, cooperates with inflammation to trigger large-scale chromatin remodeling events that promote tumor initiation. However, important questions remain: How does *KRAS*-mediated epigenetic dysregulation amplify plasticity in select epithelial cells that give rise to neoplastic lineages, and enable their subsequent evolution to invasive disease? What are cell-intrinsic and cell-extrinsic determinants of the propensity to acquire a plastic and ultimately a tumorigenic cell-state?

To shed light on early cancer-driving events encompassing genetic, environmental, and epigenetic factors, we compared physiological, pre-malignant and malignant epithelial heterogeneity using single-cell genomics, applying novel computational methods and functional perturbation in autochthonous genetically engineered mouse models (GEMMs) to identify the principles underlying neoplastic plasticity and tissue remodeling. Beyond providing a comprehensive charting of epithelial dynamics from normal metaplasia through malignant tissue states, our approach allowed us to expose, quantify and perturb early plasticity traits uniquely endowed by oncogene-environment interaction, and define molecular, cellular, and tissue-level principles of pre-malignant tumor evolution.

## Results

### Targeted high-resolution profiling of epithelial dynamics during damage-induced neoplasia

The study of epithelial dynamics in pancreatic cancer has been limited by the inability to capture early and transitional cell-states, which tend to be rare, short-lived and difficult to identify. To characterize the full spectrum of epithelial cell-states in both normal and pathological tissue remodeling, we generated a single-cell transcriptomic (scRNA-seq) atlas of healthy, regenerating, benign neoplastic, and malignant epithelia using GEMMs that faithfully model cancer from initiation to metastasis. Our GEMMs incorporate a *Ptf1a*-Cre-dependent mKate2 fluorescent reporter to enrich pancreatic epithelial cells (**Methods;** (*25, 27, 28*)), allowing us to comprehensively profile pancreatic epithelial dynamics in well-defined tissue states.

Specifically, we profiled pancreatic epithelial cells from (i) healthy pancreas (N1) undergoing reversible metaplasia associated with normal regeneration after injury (N1→N2), or (ii) the metaplasia-neoplasia-adenocarcinoma sequence that initiates PDAC in the presence of mutant *Kras* (K1→K6) (**Figs. 1A, S1A** and **Table S1**). In this setting, as in human cancer (*11*), *Kras*-mutant metaplasia is accelerated by an inflammatory insult (pancreatitis) (pre-neoplasia; K1→K2), proceeds to benign pancreatic intraepithelial neoplasia, (PanIN; K3, K4), and ultimately, malignant PDAC (K5) and distal metastases (K6; **Figs. 1A** and **S1A**).

**Figure 1.**
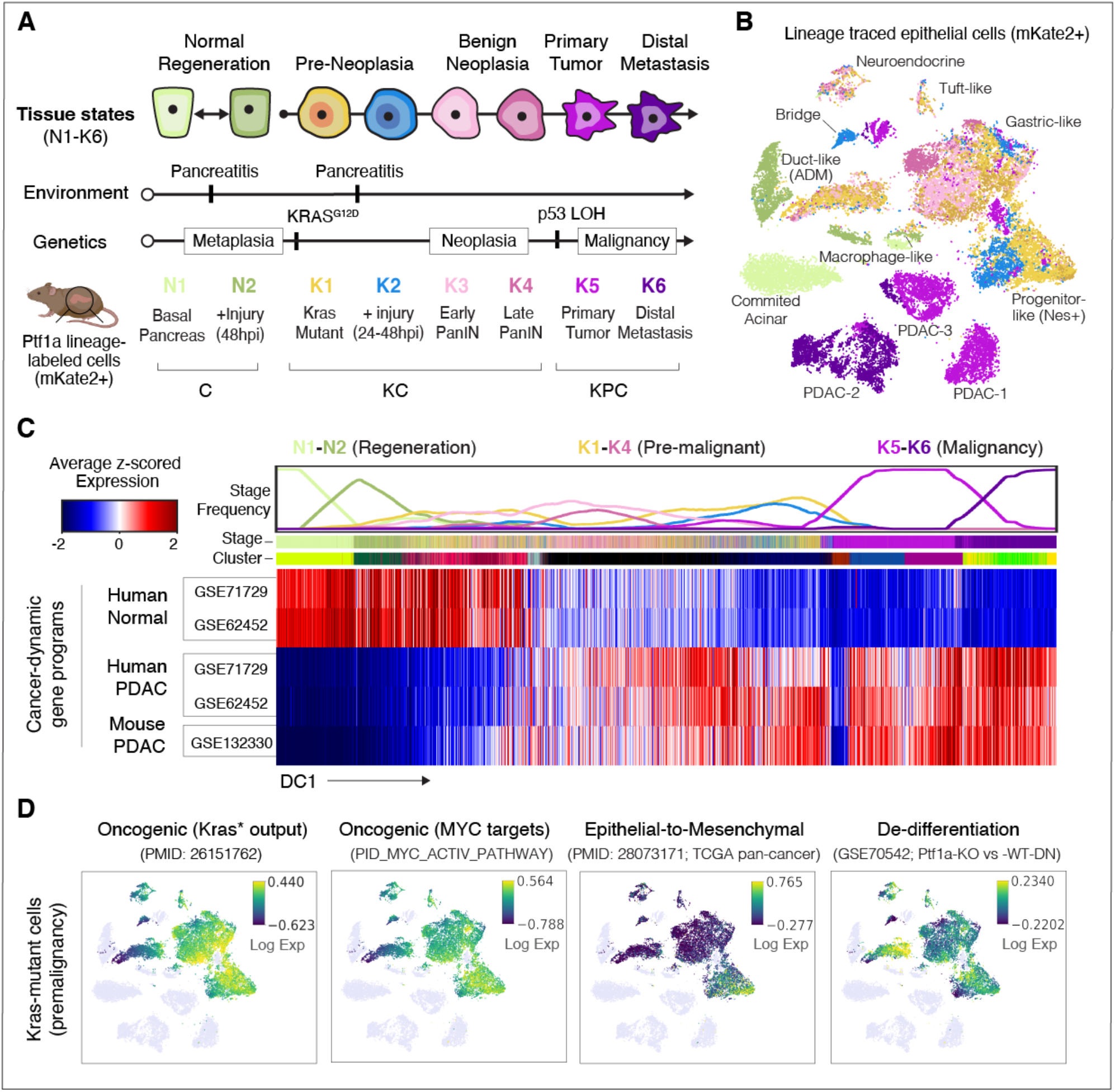
A single-cell transcriptomic atlas of pancreatic regeneration, tumorigenesis and cancer. (**A**) Experimental design for tissue collection from normal (N1) and injured (N2) pancreas, and successive stages of pancreatic ductal adenocarcinoma (PDAC) initiation and progression (K1-K6). Engineered mice expressing Cre in the pancreatic epithelium (*Ptf1a*-Cre) enable FACS-based cell enrichment of mKate2-labeled exocrine pancreas epithelial cells (genotype C) (*25*). To study physiological states, wild-type *Kras* (*Kras*^WT^) tissue was collected from this genotype in unperturbed pancreas (N1) and 48 hours post-injury (hpi) with caerulein to induce pancreatitis (N2), which causes normal metaplasia and subsequent regeneration. To study neoplastic states, tissue from aged- and sex-matched *Ptf1a*-Cre mice bearing an additional mutant *Kras^G12D^* allele (genotype KC) was collected before (K1) or 24-48 hpi with caerulein (K2), which dramatically accelerates tumorigenesis; resulting early benign PanIN lesions were collected 3 weeks after injury (K3) or from KC pancreata without injury after 27 weeks (K4). KC mice harboring a *p53* floxed (*p53^fl/+^*) or mutant (*p53^R172H^*) allele (genotype KPC), which accelerates the transition to malignancy (*31, 85*), were collected from primary PDAC tumors (K5) or metastases in liver or lung (K6). Collection thus spans a progression from pre-neoplasia (K1, K2) to benign neoplasia (K3, K4), to full-blown PDAC (K5, K6), with normal and regenerating pancreas (N1, N2) for comparison (*7, 20, 21*). We refer to stages K1–K4 as pre-malignant, and K5 and K6 as malignant. Genotype C, *Ptf1a*-Cre / RIK; genotype KC, *Ptf1a*-Cre / RIK / LSL-*Kras^G12D^*; genotype KPC, *Ptf1a*-Cre / RIK / LSL-*Kras^G12D^* / *p53^fl/+^*or *p53^R172H^*. See **Fig. S1** for additional details. (**B**) tSNE visualization of epithelial (mKate2^+^) scRNA-seq profiles from all collected stages, colored as in (A), and labeled by cell-state annotation (see **Methods**). ADM denotes an intermediate state of cells actively undergoing acinar-to-ductal metaplasia (*20*), and ‘Bridge’ denotes cells between acinar-like and malignant-like programs. (**C**) Expression of PDAC-associated gene sets across all pancreatic epithelial (mKate2+) cells (*36, 37*). Rows correspond to gene sets; columns correspond to cells, ordered by the first diffusion component (DC1) representing the major axis of progression from normal (N1) to metastatic (K6) states. Plot at top displays frequency (ranging from 0 to 1) of cells per stage in bins of n = 2000 cells. Gene set score for each cell is computed as the average of log-normalized expression, z-scored across each gene to obtain a comparable scale; heatmap is then standardized across rows to compare each gene set across cells. (**D**) tSNE plot as in (B), with pre-malignant (K1–K4) *Kras*-mutant cells colored by the expression of (from left to right) genes upregulated in bulk RNA-seq of *Kras-*mutant pancreas tissues relative to normal (*86*); genes associated with Myc activity in the PID_MYC_ACTIV_PATHWAY signature (*87*); genes associated with EMT (*38*); and genes down-regulated upon *Ptf1a* knockout, maintaining acinar differentiation (*86*). Gene set score for each cell is computed as the average of log-normalized expression, z-scored across each gene to obtain a comparable scale. Colors are scaled from 5th to 95th percentile of expression. *Kras\** denotes mutant *Kras*.

Using our lineage tagging reporter to enrich for epithelial (mKate2+) cells, we successfully captured both abundant and rare constituents of normal, regenerating and *Kras*-mutant epithelia, such as progenitor-like tuft (*Pou2f3*^+^, *Dclk1*^+^), EMT-like (*Zeb1*^+^), neuroendocrine (*Syp*^+^ *Chga*^+^) and other previously reported subpopulations (*29–31*) (**Figs. 1B**, **S1B-E,** and **Table S10)**. We also characterized highly granular routes of acinar-to-ductal metaplasia (ADM) associated with regeneration and tumor initiation (*20, 25*) (**Fig. S2A,B**). Compared to healthy and regenerating pancreata, we uncovered a staggering expansion in phenotypic diversity at the earliest stages of tumor development, including states that only emerge during *Kras*-driven tumorigenesis (**Figs. 1B, S1B–E** and **S2A-C**). Interestingly, despite such heterogeneity, the distinct cell-states captured within pre-malignant tissues were remarkably reproducible across biological replicates (individual mice) (**Fig. S2E**). In stark contrast, and consistent with reports in patients (*32, 33*), malignant tumors isolated from different mice showed extensive inter-tumor variability, only sharing one small cell cluster (**Fig. S2D,E**). Thus, highly stereotypical changes in epithelial transcriptional diversity define pre-malignant cell dynamics, suggesting that hard-wired early plasticity mechanisms underlie tumor initiation.

We next used diffusion maps (*34*) to characterize the major axes of transcriptional variation in our data, ordering cells along components associated with coherent gene expression patterns (*35*). The top component of variation closely matches progression from normal to regenerating, early tumorigenic, and finally late-stage disease, and is consistent with gene signature changes that distinguish advanced human PDAC from normal pancreas (*36, 37*). Specifically, genes upregulated in human and mouse PDAC rise along the first diffusion component, while normal pancreas programs are downregulated (**Fig. 1C**). Consistent with prior reports that use bulk data (*25*), the combined effects of *Kras* mutation and injury-driven inflammation are sufficient to induce signatures of human PDAC in pre-malignancy, as early as 24 to 48 hours post-injury (hpi) (**Fig. 1C,D**). However, these cancer-specific signatures are not induced uniformly across pre-malignant epithelial cell-states; for example, some rare early *Kras*-mutant cells express uniquely high levels of EMT gene programs (e.g. *Zeb1*, *Vim;* (*31, 38*)), previously implicated in PDAC metastasis (*39, 40*) (**Figs. 1D** and **S1D-E**). *Kras*-mutant cell-states are also observed with varying degrees of de-differentiation (e.g. downregulation of acinar genes) and reactivation of developmental (e.g. *Clu*) or oncogenic (e.g. *Kras, Myc*) programs, among others (**Fig. 1D**). Thus, pancreatic epithelial cells undergo specific and highly reproducible changes that emerge in early tumorigenesis in conjunction with inflammation and endow select epithelial subpopulations with the capacity to activate disease-relevant programs long before malignant progression.

### Aberrant, highly plastic cell-states emerge early in PDAC progression

Although histological and transcriptomic analyses have revealed rapid phenotypic changes upon *Kras* mutation and inflammation (*6, 25, 41*), we discovered substantial variation in cells’ capacities to activate cancer-relevant programs at these early pre-malignant stages. To map the cellular origins and processes underlying this diversity we first visualized heterogeneity in all *Kras*-mutant epithelia using a force-directed layout (FDL), which emphasizes cell-state transitions along axes toward malignancy. As expected, we found that *Kras*-mutant cells undergo progressive changes in gene expression to activate metaplastic (e.g. *Clu*^+^*, Krt19*^+^; (*21, 42*)), neoplastic (e.g. *Agr2*^+^, *Muc5ac*^+^*, Tff1*^+^; (*43–45*)) and ultimately, invasive cancer (e.g. *Foxa1*^+^; (*46*)) programs (**Fig. 2A**).

**Figure 2.**
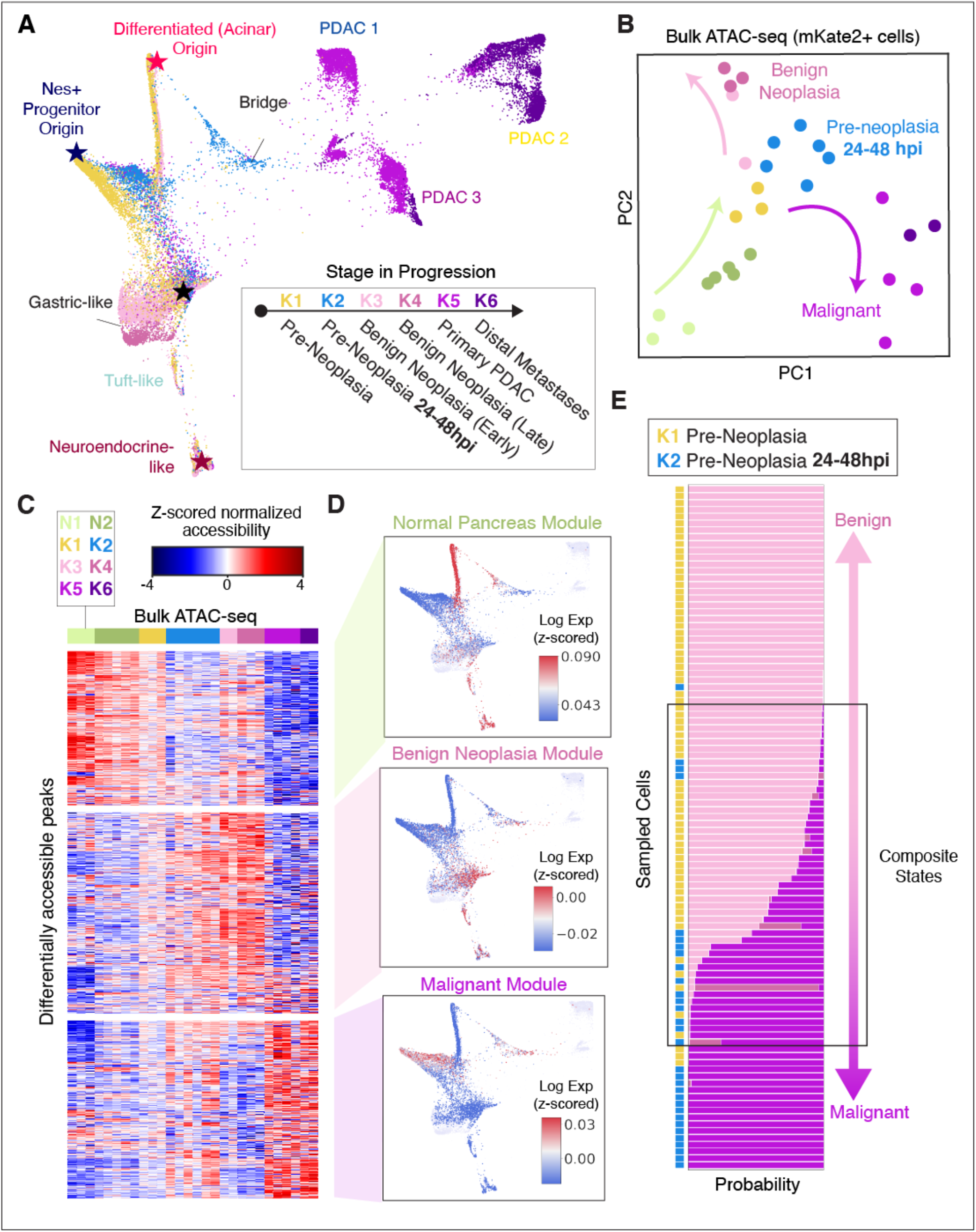
Differential epigenetic priming of *Kras*-mutant cells for fates associated with cancer-progression. (**A**) Force-directed layout (FDL) of all *Kras*-mutant scRNA-seq profiles (K1–K6). Cells colored by stage as in Fig. 1A. Stars highlight differentiated and progenitor-like ‘apex’ states inferred by CellRank (*47*) (see **Fig. S3A** for details). (**B**) Principal component analysis (PCA) of bulk ATAC-seq profiles from mKate2+ pancreatic epithelial cells computed on differentially accessible chromatin peaks. Each point shows the position of a single biological replicate (individual mice) on the first two PCs and is colored by stage as in (A). Arrows indicate longitudinal progression through PC space, including a transition upon injury and *Kras* mutation (N1-N2, K1-K2; green arrow) and a divergence between samples derived from benign neoplasia (K3-K4; pink arrow) and malignant (K5-K6; purple arrow) stages. (**C**) Chromatin accessibility along progression. Rows correspond to subsets of differentially accessible ATAC-seq peaks organized into three modules by clustering (**Methods**); columns correspond to bulk ATAC-seq replicates ordered and colored by stage as in (A). Values are normalized with DEseq2 and z-scored across rows to highlight changes in accessibility along progression. Modules exhibit distinct accessibility patterns, including peaks preferentially retained in *Kras*-wild type cells during normal regeneration (Normal Module, top), peaks appearing from injury in *Kras*-mutant cells and retained in PanIN stages (Benign Neoplasia Module, middle), and peaks appearing from injury in *Kras*-mutant cells and retained in adenocarcinoma (Malignant Module, bottom). Genes corresponding to chromatin accessibility modules are provided in **Table S2**. (**D**) Expression of genes corresponding to chromatin accessibility modules in pre-neoplastic cells (K1, K2). FDL map as in (A), colored by module expression score, displaying the heterogeneity of chromatin-associated programs in these early stages. Module expression scores are computed by z-scoring each cell to emphasize the most dominant gene programs per cell, and averaging genes mapping nearest to each peak by genomic distance within a given module. Colors are scaled between the 40th and 90th percentile of expression scores. (**E**) Classification of pre-neoplastic states toward disparate fates. Probability of classifying pre-neoplastic cells (K1, K2, labeled at left) as more similar to benign neoplasia (K3-K4) or malignant (K5-K6) scRNA-seq profiles using a logistic regression model based on expression similarity. Rows correspond to individual sampled cells, ordered from highest Benign probability (top) to highest Malignant probability (bottom); bars represent probability of classification from 0 to 1 to K3, K4, K5, or K6 labels, colored as in (A). A fraction of cells exhibit composite states with probability for both fates.

To better characterize sources of cell-state variation, we next applied CellRank (*47*), a data-driven approach that infers transcriptional dynamics from RNA velocity information ((*48, 49*); **Methods**). RNA velocities derived from the proportion of spliced to unspliced transcripts in a given cell can distinguish nascent from established transcriptional states, thus indicating likely future states in neighboring phenotypic space. CellRank integrates directional information from per-cell velocity estimates to infer transcriptional dynamics that can robustly pinpoint the origins of cell-state trajectories. Applying CellRank to early *Kras*-mutant cells acutely responding to an inflammatory insult (K1→K2) identified multiple states that potentially act as distinct origins for the observed heterogeneity (**Fig. S3A**).

These inferred origin or ‘apex’ states include both well-differentiated acinar (*Ptf1a*^+^) and de-differentiated progenitor-like (e.g. *Nes*^+^, *Aldh1b1*^+^) populations and, importantly, align with independent genetic lineage tracing experiments that have demonstrated their ‘cell-of-origin’ potential (*27, 50–53*) (**Fig. S3A,B**). Moreover, several of the inferred apex states are highly responsive to inflammation, with apparent cell-state shifts emerging in the context of tissue injury (K2). For instance, during pancreatitis, well-differentiated acinar cells generate a population intermediate for acinar (*Zg16, Cpa1*) and tumorigenesis-associated (*S100a6*) programs within 24 hpi (ADM-PDAC “Bridge”), and *Nes*^+^ progenitor-like cells shift into a state with reduced activation of tumor suppressive programs (*Cdkn2a*). Our findings thus suggest that oncogenic *Kras* enables the emergence of multiple high-potential states (not observed in healthy nor regenerating pancreata); each exhibiting unique responses to inflammatory triggers, but all upregulating cancer-associated programs.

### An epigenetic basis for high plasticity states

Given the important role for chromatin dynamics in driving neoplasia (*25*), we hypothesized that the expansive phenotypic diversity in *Kras*-mutant apex states and their injury-driven progeny arises through a diversification of permissive chromatin states. To determine how chromatin dynamics correspond to changes observed in our longitudinal scRNA-seq atlas, we first analyzed bulk ATAC (assay for transposase-accessible chromatin) sequencing data matching the above tissue stages (**Methods,** (*25*)). As illustrated in **Fig. 2B**, the dominant principal components of variation revealed global accessibility patterns characteristic to each stage of progression, including a striking, previously unappreciated, divergence between benign neoplasia precursor (PanIN, K3-K4) and full-blown adenocarcinoma (K5-K6) landscapes (**Fig. 2B**). This divergence suggests that patterns of chromatin accessibility at distinct sets of regulatory elements underlie the emergence of benign or malignant phenotypes. In agreement, a large set of regulatory elements exhibit mutually exclusive accessibility patterns consistent with this; for example, PanINs display increased chromatin accessibility at a number of loci that remain inaccessible in PDAC tumors (Benign Neoplasia chromatin module) and vice versa (Malignant chromatin module) (**Fig. 2C** and **Table S2**). These data are inconsistent with a predominantly linear phenotypic transition from ADM to PanINs and PDAC, and as such we hereafter refer to these as distinct ‘cell-fates’.

These diverging chromatin accessibility patterns of *Kras*-mutant states are induced remarkably early during tumor development, in pre-neoplastic cells. Bulk ATAC-seq data shows an initial increase in accessible chromatin in both fate-associated modules from samples collected 24-48 hpi, or without injury—well before PanIN or PDAC fates emerge (**Fig. 2C**). Moreover, mapping chromatin states to single-cell gene expression in these pre-neoplastic (K1-K2) cells (**Methods**) reveals that late-stage module-associated genes are expressed and restricted to transcriptionally distinct populations within 24-48 hours post-injury (hpi) (**Fig. 2D** and **Table S2**). These observations imply that the transcriptional diversity of pre-neoplastic *Kras-*mutant cells is established at the chromatin level and involves activation of fate-associated genes prior to the emergence of PanIN or PDAC lesions.

We postulated that pre-neoplastic *Kras*-mutant cells (K1, K2) that express gene programs associated with the chromatin landscape of a single distinct fate (Benign vs. Malignant) may be transcriptionally primed toward one or the other fate, such that they will have greater propensity to acquire that phenotype over time or in response to certain exogenous triggers. To assess this globally across the transcriptome, we reasoned that similarities in gene expression of pre-neoplastic *Kras*-mutant cells to later neoplastic stages (K3–K6) would indicate such fate potential. We therefore developed a classification-based approach that first identifies gene expression patterns that accurately discriminate between cell populations in advanced neoplasia, and then uses these programs to assign cell fate probabilities to pre-neoplastic cells whose fates are unknown but are foretold by the activation of fate-associated genes. Specifically, we trained a logistic-regression classifier to distinguish between benign neoplasia (K3, K4) and malignancy (K5, K6), and used it to classify pre-neoplastic (K1, K2) cells (**Methods**). This classifier is highly accurate (99%) in assigning fate to PanIN and PDAC cells of known fate and identified a set of discriminative genes which have been linked to either PDAC or PanINs (**Fig. S3C)**. Applied to pre-neoplastic cells, this approach indeed pinpoints *Kras*-mutant cells that are strongly skewed toward one or the other fate (**Figs. 2E** and **S3D**).

As expected, most cells are classified exclusively toward a single fate, with cells responding to inflammation (K2) attaining higher probability toward a malignant fate. We also identified an intriguing set of *Kras*-mutant cells that are not well classified (**Figs. 2E** and **S3E,F**), the majority of which express a composite program of otherwise divergent fate-associated genes (**Fig. S3G**). Consistent with their potential to produce diverse neoplastic phenotypes, these dual-primed subpopulations exist largely in the absence of tissue damage and overlap with initiating apex states (*Ptf1a^+^*acinar and *Nes^+^* progenitor) captured independently by CellRank (**Fig. S3A,B**). Collectively, our results imply that: (i) tumorigenesis can proceed from multiple well-differentiated or progenitor-like *Kras*-mutant cell populations in the pancreatic epithelium; (ii) neoplastic progression is not dictated solely by cell intrinsic determinants but is impacted by additional signals, here arising from inflammation, that shape the fates of epigenetically primed subpopulations; and (iii) this priming can be predicted in pre-neoplastic *Kras-*mutant cells prior to neoplastic development.

### Epigenetic plasticity is enhanced by inflammation

To map the epigenomic landscape at higher resolution, we generated single-cell chromatin accessibility (scATAC-seq) profiles of pre-neoplastic (K1), pre-neoplastic inflamed (K2), benign neoplastic (K3) and adenocarcinoma (K5) epithelia. Consistent with an epigenetic basis for the observed pre-malignant diversity (see **Fig. 2**), we found considerable heterogeneity in the chromatin accessibility within *Kras*-mutant epithelial cells at each stage (**Figs. 3A** and **S4A**). Moreover, a major axis of variation in accessibility reproduces the divergence between benign and malignant fates seen in bulk analyses (**Figs. 3B** and **S4B**). The scATAC-seq data supports *bona fide* epigenetic priming of divergent fate-associated programs in early, pre-neoplastic *Kras*-mutant cells. Specifically, we found substantial variation in accessibility near fate-associated genes across both stages and clusters (e.g. benign-associated *Tff2* and malignant-associated *Vim,* **Fig. S4C**), with *Kras*-mutant apex cells exhibiting a composite state defined by open chromatin at benign-associated and malignant-associated loci. This pattern extends to variation in open chromatin near other genes that define the benign and malignant chromatin modules (**Fig. S4D**).

**Figure 3.**
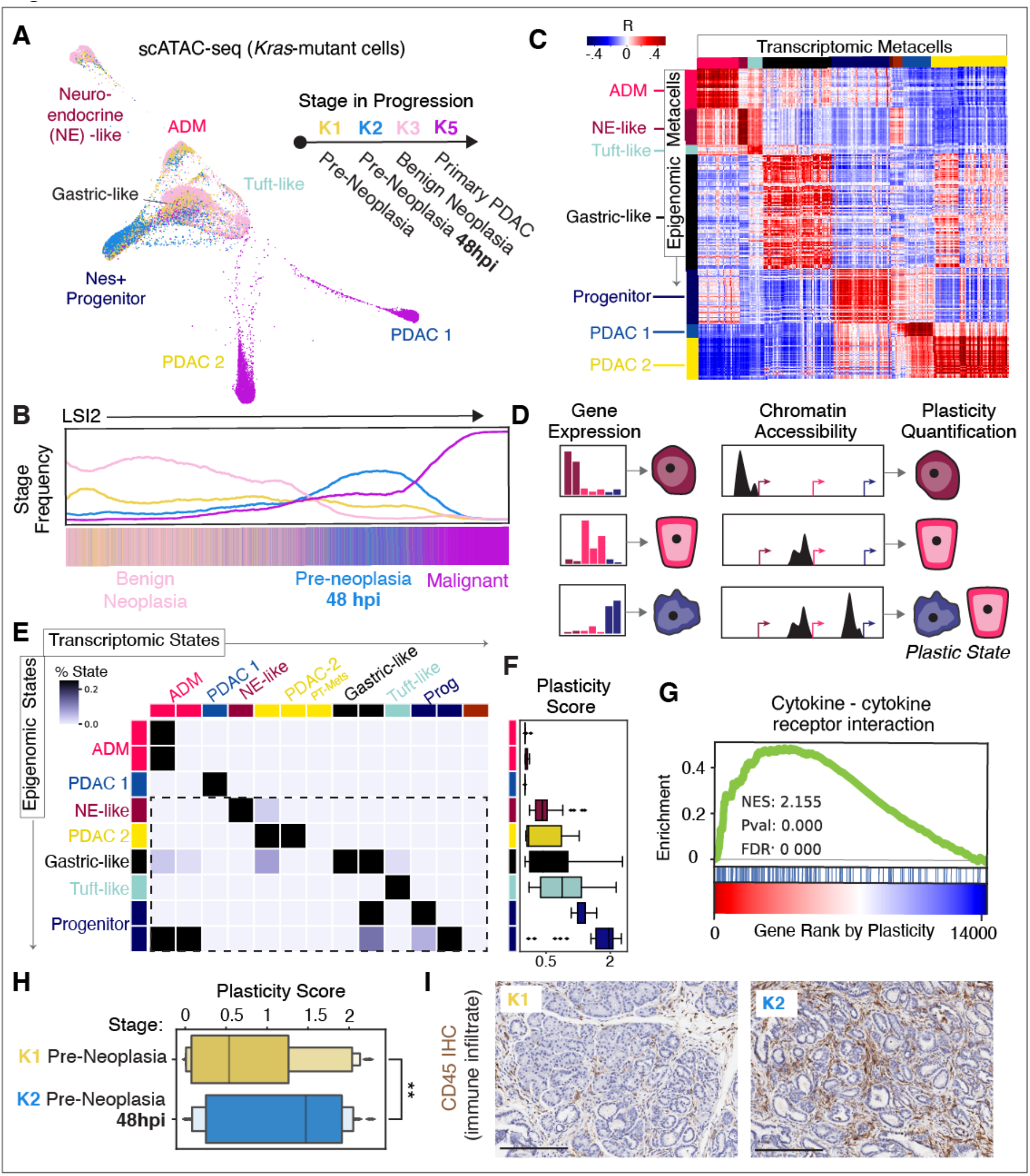
*Kras*-mutant cells display elevated epigenetic plasticity, which is associated with cell-cell communication propensity. (**A**) FDL of single cell chromatin accessibility (scATAC-seq) profiles from *Kras-*mutant epithelial cells (mKate2+) from pre-malignant (K1–K3) and malignant (K5) stages. Cells are colored by stage. (**B**) Frequency of cells of a given stage, as a function of cell order along the second high-variance component from latent semantic indexing (LSI) of scATAC-seq profiles. Benign (K3) and Malignant (K5) cells diverge at the chromatin level, with inflammation (K2) shifting pre-neoplastic *Kras*-mutant cells toward a malignant-like state along this axis. (**C**) Pairwise Pearson correlation coefficients of metacells derived from scATAC-seq (rows) and scRNA-seq (columns) data. Metacells are ordered by manually annotated Phenograph clusters (see **Figs. S1B** and **S4A**). For each vector of ArchR gene accessibility scores in one epigenomic metacell, correlation is computed to the vector of gene expression values in one transcriptomic metacell across all genes captured in both modalities. Major blocks of high positive correlation along the diagonal represent accessibility- and expression-derived cell-states that are highly similar across the two modalities, whereas high off-diagonal correlations indicate relationships across distinct cell-states at the chromatin and/or expression level. (**D**) Logic for quantifying epigenetic plasticity. A classifier is trained to learn gene features which best discriminate cell-state in transcriptomic metacells. This trained classifier is then applied to features derived from chromatin accessibility measured in the vicinity of classifier genes. The classifier will accurately assign transcriptomic cell-states to those epigenomic states which display open chromatin primarily near highly predictive marker genes (top two examples), whereas those states with additional primed chromatin near un-expressed genes will have fuzzy classification to multiple cell-states (bottom example). These latter cases are indicative of plastic cell-states. (**E**) Classifier confusion matrix mapping from epigenetic to transcriptomic cell-states based on the procedure in (D). Rows correspond to scATAC-seq Phenograph clusters (see **Fig. S4A**) and columns correspond to scRNA-seq Phenograph clusters (see **Fig. S1B,C**), colored by annotated cell state. Matrix values represent the number of cells from a given epigenomic cluster that map to a transcriptomic cluster in the classification model, normalized across rows. Epigenomic clusters are ordered from top to bottom by increasing plasticity (Fig. 3F), revealing their tendency to map to multiple transcriptomic cell-states. Dashed line highlights high plasticity epigenomic states. (**F**) Chromatin plasticity score boxplots for each epigenomic cluster in (E). Boxes represent interquartile range of plasticity scores for all epigenomic metacells assigned to that cluster, computed as the per-cell Shannon entropy in the classifier’s predicted probability distribution across all transcriptomic states. Lines represent medians and whiskers represent 1.5 times the interquartile range. (**G**) GSEA enrichment plot for the KEGG “Cytokine-Cytokine Receptor Interaction” gene set (*88, 89*). Genes are ranked along the x-axis by their Spearman rank correlation with chromatin plasticity score in epigenomic metacells. (**H**) Chromatin plasticity score boxplots for epigenomic metacells associated with K1 and K2 conditions. A significant increase in plasticity is observed in the context of inflammation (one-tailed t-test; t=2.5511, *P*=.006). (**I**) Immunohistochemistry of CD45 (brown) marking infiltrating immune cells in K1 and K2 tissue. Stronger staining for CD45 in K2 indicates the expected increase in immune infiltrate (pancreatitis) in response to injury.

To better connect primed chromatin landscapes to their transcriptional outputs, we next sought to integrate scATAC-seq and scRNA-seq profiles from comparable stages. Clustering and cell-state annotation demonstrate that cell-states derived from scRNA-seq data match those derived from scATAC-seq data at the broad cluster level, including those corresponding to *Nr5a2^+^* acinar, *Neurod1^+^* neuroendocrine, *Pou2f3^+^* tuft and *Nes^+^* progenitor cells (**Fig. S4A**). However, we also found substantial epigenomic heterogeneity within each scATAC-seq cluster. To explore this heterogeneity in more detail, we applied a recently developed algorithm that aggregates highly similar cells into granular cell states, or metacells (**Methods;** (*54, 55*)). Metacells provide much higher resolution than clusters but aggregate cells sufficiently to reduce sparsity, ensure robustness, and improve statistical power for comparison.

After separately identifying metacells for each scRNA-seq and scATAC-seq modality, we developed a framework to map between them based on similarity between a gene’s expression and its proximal chromatin accessibility (**Fig. S5A–C** and **Methods**). This integrative analysis showed the expected correspondence between the accessibility and expression programs of comparable cell-states (**Fig. 3C**). However, we also observed extensive off-diagonal correspondence, indicating that chromatin programs are shared across diverse gene expression states. Specifically, the ADM epigenomic state broadly correlates across pre-malignant transcriptional states (i.e. tuft, neuroendocrine, progenitor and gastric), reflecting the known acinar history of these *Ptf1a* lineage-sorted cells (*25, 27*). In other cases, these correspondences may indicate widespread transcriptional poising of regulatory elements near unexpressed genes. Such effects were particularly evident in apex *Nes*^+^ progenitor cells, which exist in pre-neoplastic tissues but establish chromatin landscapes at 48 hpi that are highly correlated with those of late-stage malignant populations (**Figs. 3C** and **S4B**). This further suggests that distinct subpopulations arise in pre-neoplastic *Kras*-mutant cells that are epigenetically primed to engage neoplasia programs that arise later in progression (see **Fig. 2D**).

We sought to quantify the concept of epigenetic plasticity, which we define as the diversity of transcriptional phenotypes that is enabled (or restricted) by a given chromatin accessibility landscape. To first determine these potential transcriptional phenotypes, we used a simple classifier to identify gene expression patterns that discriminate cell-states. Assuming that proximal open chromatin conveys the potential for a gene’s activation, we then applied the classifier to predict cell-states based on accessibility proximal to genes, rather than gene expression (**Fig. 3D**). We reasoned that for a given epigenomic state, uncertainty in such predictions serves as a measure of epigenetic plasticity. Following this logic, high plasticity is characterized by many accessible loci that define multiple discrete transcriptional states and thus produce high classifier prediction uncertainty, whereas low plasticity is defined by restricted potential diversity and prediction certainty. Applying this approach to epigenomic metacells identified populations of varying plasticity (**Figs. 3E,F** and **S5D**), with the most plastic states exhibiting striking overlap with the apex cells identified by CellRank (e.g. *Nes^+^* progenitors) and experimentally validated cells-of-origin from lineage tracing studies (e.g. *Pou2f3^+^* tuft cells; (*56*)) (**Fig. 3F**).

In sum, multiple orthogonal analyses consistently identify a few discrete, highly plastic progenitor states in the context of *Kras* mutation. These inflammation-sensitive populations are predicted by CellRank as apex states, they overlap remarkably with lineage tracing cell-of-origin studies, coincide with epigenetic priming towards disparate neoplastic fates, and exhibit the highest predicted epigenetic plasticity scores.

To characterize unifying hallmark features of distinct plastic cell states, we used gene set enrichment analysis (GSEA) (*57*) to identify gene signatures within populations displaying high plasticity scores. This analysis revealed robust and consistent upregulation of sets related to cell-cell communication (**Fig. S5E**), with Cytokine-Cytokine Receptor Interaction yielding the top association (normalized enrichment score = 2.155, adjusted *P* = 0.000) (**Fig. 3G**). A substantial fraction of these plasticity-associated genes encoded inflammatory mediators, receptors or ligands involved in cell-cell communication, including those previously associated with malignant progression (e.g. *Csf2*, *Cxcl1*, and *Cxcr2* (*58–60*)) (**Table S3**). Accordingly, plasticity increases significantly (one-tailed t-test; t = 2.5511, *P* = .006) upon injury in the context of *Kras* mutation (K2 vs. K1) (**Fig. 3H**), suggesting an interplay between highly plastic cells and immune infiltrates flooding the pre-neoplastic tissue environment in this context (**Fig. 3I**). Together, our results indicate that neoplastic epithelial plasticity is directly associated with an increased, epigenetically-encoded propensity for ligand-receptor mediated communication with the immune microenvironment.

### Calligraphy charts cell-state-specific communication repertoires and their interactions across cell-states

The dominance of the association between plasticity and cell-cell communication signatures drove us to investigate how heterotypic interactions may enhance plasticity in the pancreatic epithelium. We hypothesized that epigenetic remodeling of receptor and ligand gene loci (hereafter, ‘communication genes’) contributes to plasticity in pre-neoplasia by enabling cells to respond to inductive signals from the environment.

We therefore began by characterizing communication gene accessibility and expression across the pre-malignant epithelium. Each plastic cell-state reveals substantial variability in chromatin accessibility near communication genes, consistent with the maintenance of a unique molecular repertoire for potential communication (**Fig. S6A**). To identify trends suggestive of coordinately regulated gene expression, we searched for co-expression between any two communication genes (testing all combinations of two receptors, two ligands, and one of each) in individual cells across the pre-malignant epithelium and found a high degree of modularity in pairwise co-expression. This expression modularity implies that communication capabilities are driven by modules, or sets of communication genes that are mutually expressed in the same cell populations (**Fig. 4A**). We next sought to infer actual cell-cell signaling interactions that may occur between cells expressing different communication gene modules (or between a module and itself, in the case of autocrine signaling).

**Figure 4.**
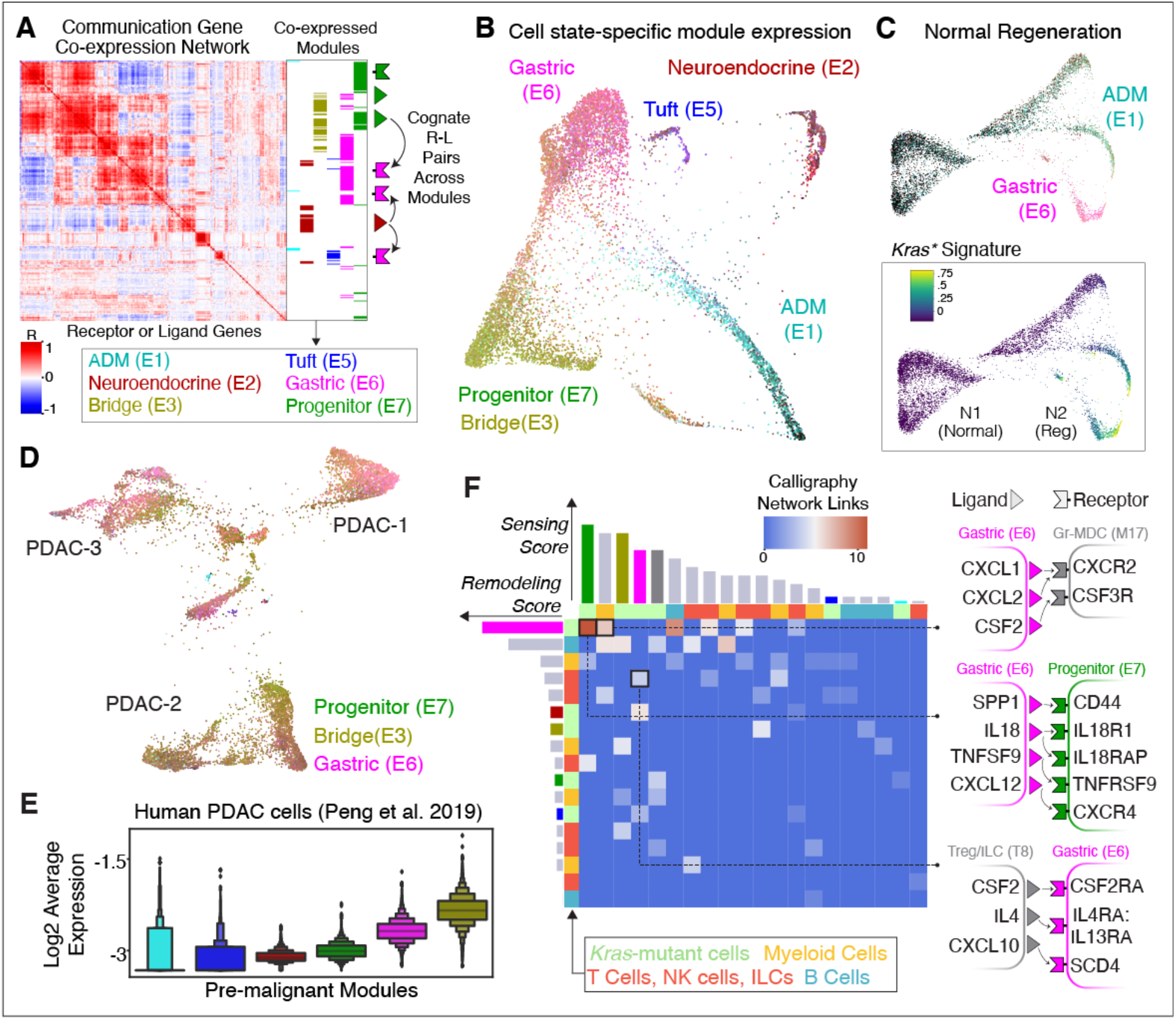
Inferred epithelial-immune crosstalk reveals chromatin-encoded cell-cell interaction promiscuity associated with plastic neoplastic states. (**A**) Receptor and ligand (‘communication’) gene co-expression modules in pre-malignant *Kras-* mutant (K1-K3) epithelial cells inferred by Calligraphy. Each row or column corresponds to one receptor or ligand. Color values represent the Pearson correlation coefficient between the expression of a pair of genes across pre-malignant cells. Rows and columns are ordered according to hierarchical clustering using centroid (UPGMC) linkage. Blocks of highly correlated communication genes along the diagonal correspond to partially overlapping modules of genes that tend to be expressed in the same cell populations. Each column of inferred communication gene co-expression modules from the OSLOM community detection algorithm (right) depicts genes belonging to a single module. Modules annotations (bottom) are based on restricted module expression in specific cell-states (see **Fig. S6B,C**). A cartoon schematic at far right indicates the second step of Calligraphy, which uses prior knowledge of cognate receptor-ligand (R-L) binding relations to infer crosstalk between modules. (**B**) FDL of pre-malignant *Kras-*mutant (K1–K3) epithelial cells, colored by communication module expression as in (A). Module expression is computed as the log of average normalized expression of each gene in the module. Cell “pseudo-color” values correspond to a mixture of module-specific colors based on the relative expression of module genes in each cell (**Methods**). The cell-state-specific nature of expression for distinct modules is revealed by the lack of blending of module colors. (**C**) FDL of normal (*Kras*^WT^) pancreas cells before and after injury (N1-N2), colored by pre-malignant communication module expression as in (A-B) (top) or average, z-scored log-normalized expression of mutant *Kras* signature genes (bottom, (*25*)). Colors in the bottom plot are scaled between the 1st and 99th percentile. Normal healthy cells (N1, Normal) lack expression of genes from any module (dark shading), whereas states induced during physiological regeneration (N2, Reg) acquire significant ADM (E1) and Gastric (E6) communication module expression coinciding with high *Kras-*mutant (*Kras*\*) signature expression. (**D**) FDL of malignant (K5-K6) cells, colored by communication module expression as in (A). Module expression is computed as the log of average normalized expression of each gene in the module as in (B). Cell color values correspond to a mixture of module-specific colors based on the relative expression of module genes in each cell (**Methods**). Progenitor (E7), Bridge (E3), and Gastric (E6) modules are highly expressed amongst all three mice. (**E**) Pre-malignant communication module expression in human pancreatic tumor scRNA-seq data (*32*). Module expression for each cell is computed as the average normalized expression of each gene in the module. Higher values of expression for Progenitor (E7), Bridge (E3), and Gastric (E6) modules indicate that genes within these modules are highly expressed across multiple advanced PDAC tumors. (**F**) Pairwise crosstalk relationships between epithelial and immune communication modules inferred by Calligraphy. Each row or column represents one module computed from either epithelial or immune scRNA-seq data; modules with no incoming or outgoing edges are excluded from columns and rows respectively. Color values represent the number of edges (cognate R-L pairs) spanning statistically significant pairs of communicating modules, indicating strength of interaction. Bars along rows and columns display sums of statistically significant edge counts, representing the total number of remodeling (transmitting) interactions (rows) or sensing (receiving) interactions (columns) for that module. Bars are colored by module as in (A) for epithelial modules or gray for immune modules. For select interactions, specific cognate R-L pairs contributing to the interaction are shown at right.

Although several methods have been developed to predict cellular interactions from single-cell data (*61, 62*), their inference relies on weak signals based on the noisy expression of a single cognate receptor-ligand (R-L) pair across fixed cell-states. We therefore developed a new approach, Calligraphy, that leverages the observed modularity in communication gene expression to infer potential cell-cell signaling events (**Methods**). Calligraphy first identifies communication modules—sets of communication genes that are co-expressed across cell-states—thereby establishing the incoming and outgoing communication each cell-state can participate in. Next, Calligraphy identifies communication events between cell-states based on prior knowledge of cognate R-L binding partners (**Fig. S9A**).

Using Calligraphy, we obtained seven communication modules of genes that are co-expressed across the pre-malignant pancreatic epithelium (**Fig. 4A**). Mapping average expression of communication genes back onto the pre-malignant epithelium, we found that most cells express a single dominant module, making it possible to annotate cells by their corresponding module (**Fig. 4B**). Strikingly, cell-states defined solely by communication gene expression coincide with those identified by clustering the entire transcriptome (**Fig. S6B,C**). We observe the same patterns in chromatin-state subpopulations, where each chromatin subpopulation can be defined through patterns of open chromatin around distinct communication modules, supporting an epigenetic basis for the emergence of these programs (**Fig. S6D**). Collectively, these observations suggest a role for cell-cell communication in defining global cell-state heterogeneity.

Consistent with the potential importance of communication modules in driving early neoplasia, normal and regenerating epithelial cells (N1, N2) showed much less diversity in communication module expression, with the majority of cells maintaining very low module expression compared to their neoplastic counterparts (**Fig. 4C**). Among cells with wild-type *Kras*, a small injury-induced population does express modules associated with ADM (E1) or Gastric (E6) cell-states, likely reflecting the expected trans/dedifferentiation of acinar cells under inflammatory conditions (*63, 64*). The same wild-type cells express high levels of a mutant *Kras* output signature (*25*), suggesting that *Kras* mutation stabilizes otherwise transient, injury-induced communication modules during tumor initiation. Consistent with this notion, a substantial fraction (20%) of pre-malignant module genes are associated with changes in gene expression that distinguish tumorigenesis from regeneration (**Table S4**; (*25*)), and with the greater cellular diversity of pre-malignant tissues. Moreover, communication modules established in the pre-malignant pancreas are maintained in advanced cancers (K5, K6), with most cells expressing at least one of the Gastric (E6), Progenitor (E7) or Bridge (E3) modules (**Fig. 4D**). These same early-established communication modules are conserved in human PDAC derived from multiple patients (**Fig. 4E**). The unique behavior of communication modules in early neoplasia and their persistence in advanced murine and human PDAC suggests a functional role in pancreatic tumorigenesis.

### Extensive epithelial-immune interactions drive oncogenic tissue remodeling

Our data raise the possibility that communication modules that arise in *Kras*-mutant epithelial cells during inflammation are important for driving cell-state plasticity during early neoplasia and, by extension, collaborate with pancreatitis to promote PDAC development (*7, 23, 24*). Tissue damage produces inflammation and changes in immune cell-state and composition that contribute to neoplasia (*65, 66*); we thus set out to investigate how mutant *Kras*-driven epithelial communication modules interact with infiltrating and tissue-resident immune cells. scRNA-seq analysis of immune cells (CD45^+^ sorted) from pre-neoplastic *Kras*-mutant tissues, before and after induction of pancreatitis (K1–K3), identified all expected immune subtypes, including both abundant (macrophage) and rare (Treg, ILC) types whose profiling was enabled by CD45^+^ enrichment (**Fig. S7A-C** and **Methods**). As expected, injury-induced inflammation causes dramatic remodeling of the immune cell landscape, including the enrichment or depletion of specific lymphoid and myeloid cell-states (**Fig. S8A**).

Applying Calligraphy to these data identified consistent and structured communication modules among distinct immune populations. To achieve even greater resolution, we ran Calligraphy separately on T cells/ILCs/NK cells, myeloid cells, and B cells, and found numerous modules containing known regulators as well as novel candidates of pancreatic tumorigenesis (**Fig. S8B,C**). For example, myeloid module 20 is mainly expressed in macrophages of injured *Kras*-mutant epithelium (absent from uninjured pancreas) and includes the type 2 IL-4 receptor (*Il4r*/*Il13ra1*) and CSF2 receptor (*Csf2ra*/*Csf2rb*) genes as well as *Mif* (*67*) and *Cxcl1* (*68*), which have known roles in advanced PDAC progression. Within the lymphoid (T/ILC/NK) compartment, module 8 is highly expressed in ILC2, ILC3/LTi and Treg cells; these cells express the receptor for IL-33 (*Il1rl1*/*Il1rap*), a ligand that accelerates the formation of mucinous PanIN lesions (*25*).

Reasoning that such rapid immune and epithelial remodeling could arise through heterotypic crosstalk in pancreata undergoing neoplastic transformation, we utilized a feature in Calligraphy that nominates potential cell-cell interactions that drive this process (**Fig. S9A** and **Methods**). We first limited our search to crosstalk events triggered in the epithelium through a combination of *Kras* mutation and inflammation, by filtering Calligraphy modules to retain those cognate R-L pairs in which at least one partner is significantly upregulated in inflamed pre-neoplastic cells (K2) relative to normal regeneration (N2). This filtering step reduced the space of possible interacting molecules from 340 total communication genes down to 55 receptors and 46 ligands potentially involved in tumorigenesis-associated communication. We then assumed that two modules potentially interact across cell subsets if they are significantly enriched in the number of shared cognate R-L pairs spanning them (**Methods**).

Calligraphy identified a network of potential neoplasia-specific interactions between the *Kras*-mutant epithelium and the immune environment, revealing the emergence of ‘master communication hubs’ that participate in numerous interactions (**Fig. S9B,C**). We calculated a receiving score (ability to sense the environment via expressed receptors) and a transmission score (ability to remodel the environment via expressed ligands) based on the number of statistically significant incoming and outgoing edges for each module (**Methods**). Interestingly, the two most prominent hubs for transmitting and receiving interactions are the epithelial Gastric (E6) and Progenitor (E7) modules, respectively (**Fig. 4F**), which notably correspond to ‘high-plasticity’ populations identified above. These same communication hubs are enriched in advanced mouse and human PDAC (**Fig. 4D,E**).

### Neoplastic tissue remodeling involves feedback communication loops between epithelial and immune compartments

One of our most striking observations is the dramatic remodeling of epithelial and immune compartments within 24 to 48 hours of injury (see **Figs. 1A,B** and **S8A**). This remodeling is expansive, involving numerous new cell states that are quickly adopted by a majority of cells, and it is highly reproducible. The dynamics of such a rapid and robust response suggest a feedback loop, by which immune cell intermediates may amplify tumor-promoting epithelial cues (*69*). We probed the Calligraphy module-module interaction network to systematically enumerate cycles involving any epithelial or immune subsets, and identified only one putative tumorigenesis-associated feedback loop in the system (**Fig. 5A** and **Methods**). A unique strength of our approach is the ability to dissect the complexity inherent to tissue crosstalk, by employing modules that encode multiple putative interactions and by linking signaling events between populations in serial fashion.

**Figure 5.**
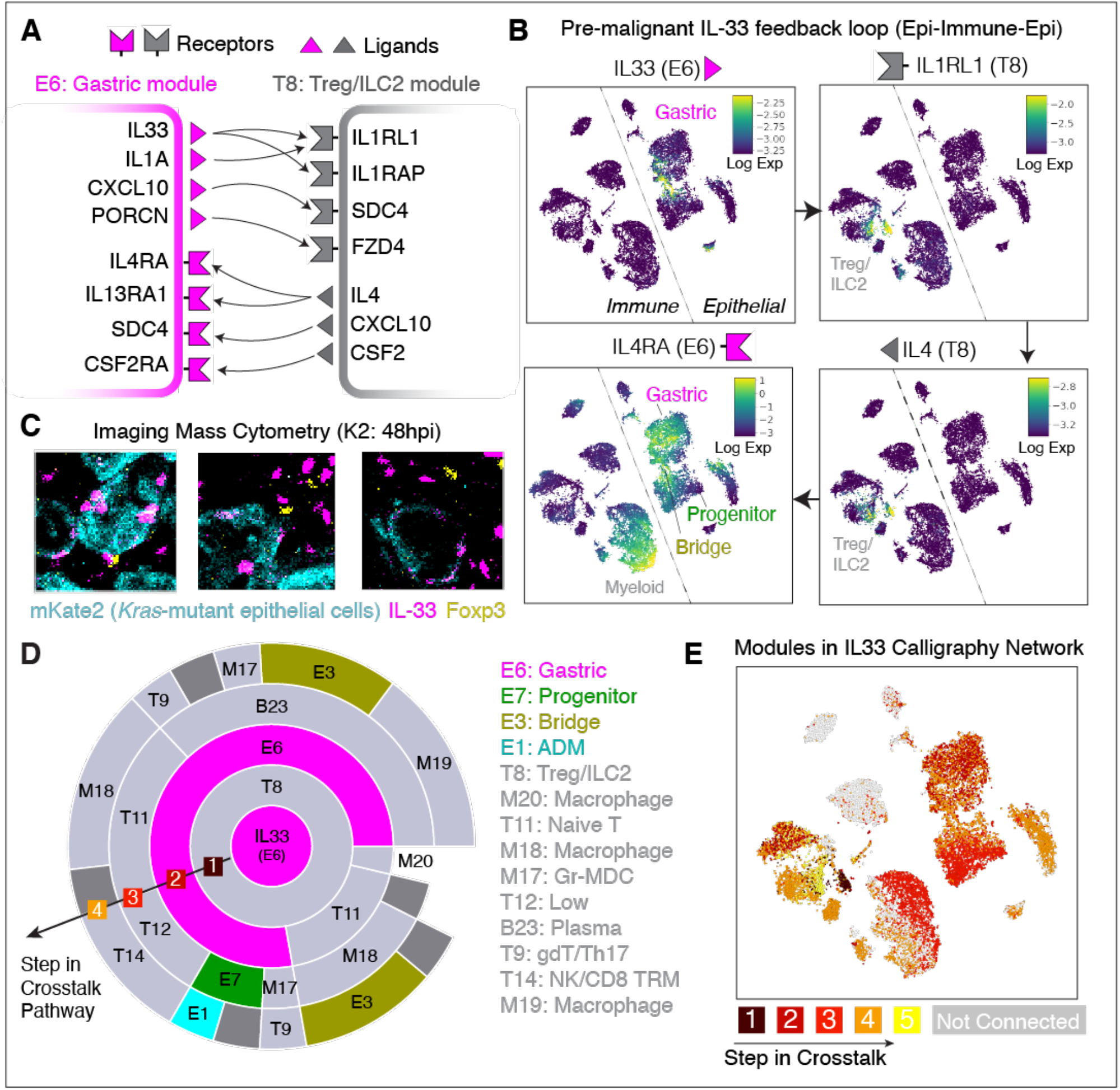
A feedback loop between *Kras*-mutant epithelial states and immune populations has widespread impact. (**A**) A feedback loop identified by Calligraphy in the pre-malignant pancreas in which receptor and ligand genes in Module E6 (Gastric, left) communicate with those in Module T8 (Treg/ILC2, right). Arrows depict cognate signaling partner interactions. (**B**) tSNE of integrated immune and epithelial scRNA-seq data from pre-malignant (K1-K3) stages displaying MAGIC (*90*) imputed expression of key genes from the feedback loop in (A). Each cell is colored by log-normalized expression of the indicated gene, and arrows between plots indicate subsequent steps of the loop. Genes within the same module (along columns) tend to be co-expressed in the same populations. *Il4ra*, the final gene in the loop, is highly expressed across many *Kras*-mutant epithelial populations. (**C**) Imaging Mass Cytometry (IMC) staining of pancreatic tissues from the K2 stage (*Kras*-mutant pre-neoplasia 48hpi) showing IL-33 (magenta), mKate2 (cyan) and Foxp3 (green). Each panel shows an example of co-expression of IL-33 and mKate2 (marking epithelial cells), and apposition of Foxp3-expressing Tregs and IL-33-expressing epithelial cells, supporting their potential for direct communication. (**D**) IL-33-centric crosstalk paths originating from epithelial cells. Central circle denotes the IL-33 source (epithelial Gastric module E6, magenta), and each outward concentric circle illustrates possible communication paths from inner to outer modules based on significant crosstalk links inferred by Calligraphy. The arc length of each neighboring module pair is proportional to the number of inner-module ligands that can bind to cognate receptors in the outer module, and hence is proportional to the strength of the inferred connection. (**E**) tSNE as in (B), colored according to the step in which communication events from the IL-33-centric path in (D) reaches the module expressed by that cell. Module expression is computed as the log of average normalized expression of each gene in that module; cells are assigned to the highest-expressed module. The module is then scored by the earliest step at which it appears along any paths through the Calligraphy network emanating from E6-derived IL-33. Cells expressing modules which are not downstream of IL-33 are colored in gray.

The loop comprises the Gastric (E6) hub module, which is maintained in late disease, and further involves cytokines with reported roles in pancreatic cancer or pancreatitis, including IL1A and IL-33 (*70–72*). Specifically, it engages Treg and ILC2 cells via IL-33 signaling before feeding back to epithelial cells (**Fig. 5A**). The importance of this IL-33-mediated loop is further supported by our previous work showing that exogenous IL-33 accelerates the *Kras*-driven transition from metaplasia to PanINs (*25*).

*Il33* is expressed during pancreatitis by a small subset (4%) of TFF1/ANXA10^+^ *Kras*-mutant epithelial cells within the Gastric module and is predicted to initiate signaling to Tregs and ILCs (Module T8) by binding with its cognate receptor *Il1rl1* and co-receptor *Il1rap* (**Figs. 5B** and **S10A-C,F**). Supporting the relevance of these interactions, imaging Mass Cytometry (IMC) data collected using a customized murine antibody panel reveal that IL-33-expressing epithelial cells (IL-33^+^ mKate2^+^) and Tregs (Foxp3^+^) are in close spatial proximity in *Kras*-mutant pancreata under injury conditions (**Fig. 5C**). Subsequently, many receiving cells that express *Il1rl1* (Module T8) also express the Th2 cytokine gene *Il4* (Fisher’s exact test; odds ratio = 21.47, *P* = 9.88 x 10^-35^), consistent with the known role of IL-33 in triggering Th2-type immune responses (*73*). Module T8 cells then apparently signal, largely through IL-4, back to the Gastric Module (E6), thereby closing the loop and potentially propagating signals to other modules in both immune and epithelial compartments (**Fig. 5B**).

The broad expression of the IL-4 receptor (*Il4ra*) across *Kras*-mutant epithelial cell-states, including gastric, *Nes*^+^ progenitor, and tuft cell populations (**Figs. 5B** and **S10A,C**), implies that this signaling loop has a system-wide impact on pre-malignant tissue (45% of pre-malignant epithelial cells appear impacted). In contrast, few wild-type normal pancreas cells express both sending (*Il33*) and receiving (*Il4ra*) factors, and do so at low levels, even during injury-induced regeneration (**Fig. S10D, E**).

Calligraphy provides the ability to map key communication circuits, including the series of populations directly and indirectly affected by epithelial-derived IL-33. IL-33-driven communication has a large predicted impact on pre-malignant tissue, and the proportion of affected tissue increases via signaling cascades between communication modules that each utilize multiple cognate R-L pairs, ultimately reaching the vast majority of the pre-malignant pancreas (72% of cells, **Fig. 5D,E**). While it is unlikely that the IL-33 loop is solely responsible for KRAS-driven tumor progression in the context of the high complexity of observed intercellular communication circuits in the pre-malignant tissue, the number of populations directly and indirectly impacted by epithelial IL-33 expression suggests that this communication circuit plays an important role in driving tumorigenesis. More broadly, our analyses establish that emergent immune-epithelial signaling networks are a prominent feature of early pancreatic neoplasia and suggest that they play important biological roles in distinguishing neoplasia from normal tissue regeneration.

### KRAS-dependent IL-33 feedback loop directs rapid tissue remodeling in early tumorigenesis

To test our prediction that epithelial-derived IL-33-driven feedback contributes to tumor initiation *in vivo*, we developed a new GEMM that enables lineage tracing and epithelial-specific *Il33* suppression in mutant *Kras* -expressing epithelium. Specifically, animals were produced from multi-allelic embryonic stem (ES) cells engineered to harbor a conditional *Kras*^G12D^ allele together with a doxycycline (dox) -inducible green fluorescent protein (GFP)-coupled short hairpin RNA (shRNA) capable of suppressing *Il33* (KC-sh*Il33*), allowing potent *Ill33* suppression following doxycycline (dox) administration (**Fig. 6A,B**). We performed both scRNA-seq and spatial (IMC) analysis on this model under *Il33* perturbation during injury-accelerated KRAS-driven neoplasia, assessing epithelial and immune compartments at an early time-point (48 hpi), when inflammation unleashes neoplastic remodeling (K2), as well as 3 weeks thereafter, when PanIN lesions normally emerge (K3). Notably, as IL-33 expression remains intact in non-epithelial pancreatic cells (*Il33^+^ Vim*^+^ or *Il33^+^ aSMA*^+^) (**Fig. 6C,D, and S11A,B**), this system allows us to uncouple the impacts of abundant stroma-derived IL-33 from those of our mutant *Kras*-associated feedback loop. Upon injury, IL-33 is expressed in a subset of *Kras*-mutant cells positive for TFF1/ANXA10+ gastric module markers (**Fig. 6B,C**), and this induction is prevented in on-dox KC-sh*Il33* mice (**Fig. 6B-D**).

**Figure 6.**
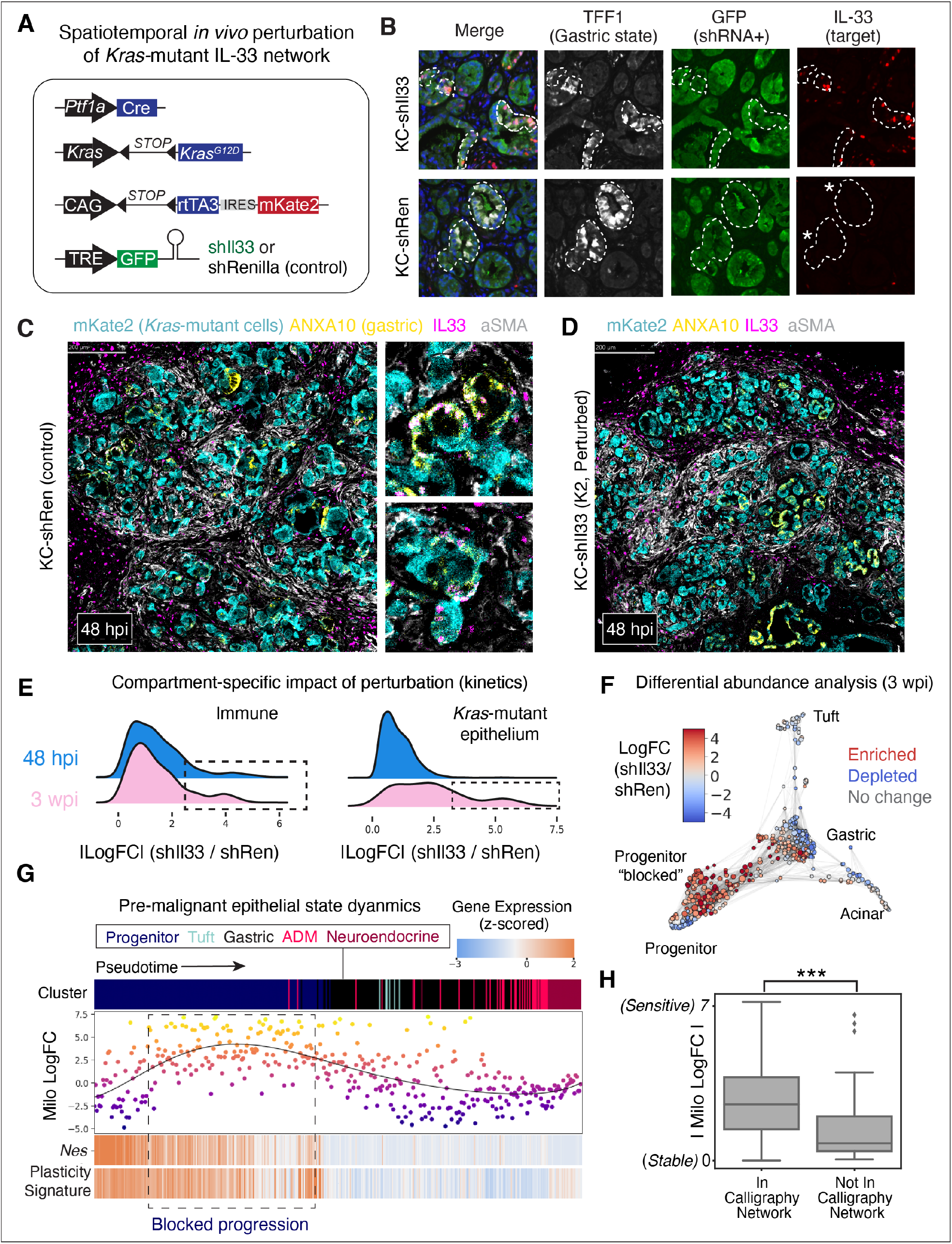
Spatiotemporal *in vivo* perturbation of *Il33* impairs neoplastic progression. (**A**) Genetic mouse modeling approach for selective and inducible targeting of *Il33* (KC-sh*Il33*, 2 independent strains) or Renilla (KC-shRen, control) in *Kras*-mutant epithelial cells, as restricted by *Ptf1a*-driven expression of Cre. (**B**) Representative immunofluorescence in pancreata from KC-shRen (top) or KC-sh*Il33* (bottom) placed on dox fed at 5 weeks of age and analyzed 9 days later, at 48 hpi timepoint (K2). GFP marks *Kras*-mutant epithelia expressing an shRNA driving potent *Il33* suppression in that same compartment (but not in surrounding stroma) in KC-sh*Il33* but not in KC-shRen control mice. TFF1 marks the *Kras*-mutant gastric epithelial cell-state in which IL-33 is activated upon injury (48 hpi) in control (top) but not sh*Il33*-perturbed (bottom) animals. Dashed lines demark boundaries between epithelium and stroma, and asterisks highlight potent suppression of *Il33* in TFF1+ metaplastic cells of KC-sh*Il33* mice. (**C**) IMC staining of pancreatic tissues from KC-shRen mouse (48 hpi, K2 stage) showing activation of IL-33 (magenta) within mKate2 (marking epithelial cells, cyan) cells positive for gastric state marker (ANXA10, yellow), as well as in surrounding stroma. (**D**) IMC staining of pancreatic tissues from KC-sh*Il33* mouse (48 hpi, K2 stage) showing depleted expression of IL-33 (magenta) in mKate2+ *Kras*-mutant epithelial cells (cyan), and intact IL-33 expression fibroblasts marked by aSMA (white). (**E**) Distribution plots of Milo (*74*) log fold change (logFC) magnitudes across cell neighborhoods. Higher |logFC| in a given neighborhood indicates a greater change in abundance comparing sh*Il33* to control conditions. Rightward shifts in the distribution thus indicate a greater degree of cell-state re-modeling upon perturbation. Top plots display distributions from K2 immune (left) and epithelial (right) cells; bottom plots display corresponding distributions for K3 cells. (**F**) Milo neighborhoods overlaid on FDL colored by Milo logFC, with increased (red) or decreased (blue) abundance in sh*Il33* samples relative to controls, at the late (3 wpi) timepoint. PanIN-associated lineages (e.g. gastric) are depleted, and with concomitant enrichment of the injury-activated *Nes*^+^ progenitor state (blocked). (**G**) Impacts of *Il33* perturbation across the *Kras*-mutant epithelial neighborhoods at K3 (3wpi) timepoint. Top indicates cell neighborhoods (columns) ordered by pseudotime and colored by cell-state annotation. Scatterplot below shows Milo logFC for each neighborhood (points also colored by logFC for emphasis). Higher logFC tracks with increased abundance in the sh*Il33* condition relative to control. A 6^th^-order polynomial regression line is fit to the trend of logFC over pseudotime. Bottom indicates z-scored, log-normalized expression of *Nestin* and average z-scored, log-normalized expression of plasticity-associated genes (Fig. 3F and **Methods**) along pseudotime. Heatmap colors are scaled to 2 s.d. above and below the mean. (**H**) Milo logFC of neighborhoods mapped to modules that are (left) or are not (right) downstream of Calligraphy’s IL-33 -centric network. Asterisks indicate statistical significance (unpaired, one-tailed t-test, *P* = 1.24 × 10^-7^).

We applied the Milo algorithm (*74*), which characterizes phenotypic shifts by grouping similar cells into local ‘neighborhoods’ and identifying those neighborhoods which are differentially abundant between perturbed and control conditions. Consistent with Calligraphy’s prediction of IL-33 signaling from epithelial to immune modules, we found that *Il33* suppression in the *Kras*-mutant epithelium results in rapid remodeling of the immune landscape; at 48 hpi (K2), multiple immune subpopulations shift significantly (**Fig. 6E**). Epithelial remodeling is delayed by comparison; a lack of significant changes at the early time point is followed by dramatic remodeling of many epithelial cell states at the 21-day PanIN stage (K3).

By 3 weeks post-injury (wpi) (K3), perturbation of IL-33-mediated crosstalk likely impacts cell-state transitions underlying neoplastic epithelial remodeling. Thus, we used Palantir (*75*) to infer trajectories between perturbed epithelial cell-states at this time point, computing a pseudotime ordering of neighborhoods beginning from the *Nes*^+^ progenitor apex state (**Fig. S11C**). Ordering Milo log-fold change values along this axis, we observed an increase in the abundance of cells from perturbation-derived samples early in pseudotime, prior to exit from the progenitor cluster (**Figs. 6F,G** and **S11C,D**). Examining gene expression programs along this axis, we confirmed that *Il33* suppression ‘blocks’ *Nes*^+^ progenitor cells in a state that is phenotypically similar to the native progenitor state (e.g. *Nes*^+^), and maintains expression of genes universally associated with epigenetic plasticity in the pancreas epithelium. Therefore, *Il33* perturbation prevents the transition of a plastic progenitor-like state into distinct populations that normally occur in PanIN, such as gastric-like cells, that are abundant in the unperturbed epithelium by 3 wpi.

Beyond delineating communication circuits, Calligraphy can also predict the specific populations impacted, as well as their relative distance along the chain of signal propagation (see **Figs. 5D,E).** To evaluate the correspondence between Calligraphy’s predictions and the observed changes under *Il33* perturbation, we mapped each Milo neighborhood to Calligraphy communication modules (see **Figs. 4B** and **S11D**) and evaluated the extent to which cell-states predicted to be downstream of IL-33-mediated crosstalk overlap with cell-states impacted by the perturbation. Qualitatively, we found the largest impact of *Il33* perturbation on Progenitor and Bridge modules, both of which are predicted to be downstream of IL-33 and express IL-4 receptor (*Il4ra)* (**see Fig. 5B,D**), whereas the only two modules not downstream of IL-33 are those with the smallest effect sizes (E2 and E5). Quantitatively, we found more robust effects of *Il33* perturbation in populations predicted to participate in crosstalk (one-sided t-test; t = -5.25, *P* = 1.24 × 10^-7^) (**Fig. 6H, F**). Thus, these results functionally validate Calligraphy as an approach to infer both communication circuits and the specific subpopulations impacted (directly and indirectly) upon perturbation of such networks.

Collectively, our findings support a role for oncogenic *Kras* in directing epigenetic changes that permit the rapid injury-triggered expression of a cytokine (IL-33) and Th2 receptors (IL-4 receptor, IL-13 receptor), which act in a cell-cell feedback loop to fundamentally remodel epithelial and immune lineages of the pre-malignant pancreas. More generally, we find that permissive chromatin states in *Kras*-mutant cells mediate diversification of communication programs in pre-neoplastic tissue, expanding channels of communication downstream of the loop and globally throughout the tissue. This work thus establishes a critical link between genetic mutation, microenvironmental insult, and epigenetic reprogramming in the highly dynamic (within 2 days) emergence and maintenance of cancer-driving populations.

## Discussion

While the research community has learned much about the genetics of PDAC development and the cellular and epigenetic processes that influence tumor progression to advanced disease, we know little about how PDAC arises from relatively homogenous epithelial cells—in human patients diagnosis occurs late, and experimental models for accessing early stages have just begun to appear. By combining deep single-cell sequencing in mouse models with powerful and novel computational approaches, we reproducibly identify select early progenitor-like cell-states primed for diverse neoplastic fates. We formalize the concept of epigenetic plasticity and reveal that these progenitor-like states exhibit enhanced plasticity, which is strongly associated with dramatic remodeling of accessible chromatin near cell-cell signaling genes. The altered communication repertoires potentiate a feedback loop between epithelial and immune cells that contributes to the neoplastic state. Our results reveal new insights into the mechanisms of pancreatic tumor initiation and have important implications for understanding the origins, output, and targeting of epigenetic plasticity during pancreatic neoplasia.

Previous studies have established that mutations in *Kras*, *p53*, or both genes drive a loss of AT1 or AT2 lineage identity in the lung within several weeks, and they define plasticity as an acquired phenotype that is intermediate between these two normal states (*18, 19, 76*). In the pancreas, loss of normal acinar identity and the presence of a similar intermediate (acinar-ductal) state have long been observed in both tumorigenesis and normal regeneration; hence, this notion of plasticity is not sufficient to fully describe PDAC disease mechanisms. Moreover, unlike previous studies (*77*), we focus on the dynamic cancer initiating events that are triggered by inflammatory processes known to initiate disease. To quantify the differences observed between injury-induced regeneration and neoplasia, we defined plasticity as the potential of a cell to manifest diverse future (neoplastic) fates, and developed a measurable plasticity score that tracks with the degree of transcriptional or epigenetic priming exhibited by a given cell-state. This score allowed us to identify highly plastic cell-states in which open chromatin unlocks access to multiple distinct programs observed in neoplasia and malignancy, and further enabled us to assess how inflammation enhances plasticity across these states. Our metric is general and can be applied to investigate plasticity and tumor progression in any cancer.

To better elucidate the emergence of these plastic states, we sought to reconcile prior work in the field nominating different cells-of-origin for neoplasia. A unique aspect of our experimental system is that all pancreatic acinar cells harbor mutant *Kras*, allowing us to comprehensively explore which cell-states can initiate tumorigenesis. Using CellRank, we were able to trace the origins of epithelial transcriptional diversity to multiple populations, which show striking correspondence with experimentally determined cells-of-origin. Indeed, our computational methods coupled to our experimental design provides a unique, high-resolution view of epithelial cell diversity that unifies the results of previous experiments by suggesting that multiple mutant *Kras* expressing cell-states can generate neoplasia.

The resolution of epithelial cells uncovered by the lineage tracing capability of our model enables this finding, but it also implies that the diverse cell-states in the *Kras*-mutant context can arise from a predominantly acinar-like state. As direct evidence of this, we observe that nearly all pre-malignant epithelial cell-states display substantial correlation with an acinar-like chromatin state (epigenetic ‘memory’). Indeed, one of CellRank’s predicted ‘apex’ states is a well-differentiated acinar-like state, which exhibits lower overall plasticity than the dedifferentiated states unique to the oncogenic context. Together, our results reflect prior knowledge that *Kras* mutation induces loss of normal lineage identity, but further reveal the emergence of *new* apex progenitor-like states that are primed for both neoplastic and malignant states. Dual-primed cells go on to generate the observed heterogeneity in benign and malignant neoplastic states.

Critically, multiple unbiased analyses converge in support of the CellRank-identified apex inflammation-sensitive *Nes*^+^ progenitor state (*52, 53*). We show that this state is the first to exhibit PDAC-associated chromatin alterations, it expresses progenitor-like gene programs, scores highest for our plasticity metric, and expresses a large number of cell-signaling receptors—all hallmarks of highly plastic cells.

The genes that correlate most strongly with our plasticity score are cytokine and receptor genes, such that plastic populations are primed to both respond to signals from the environment and to remodel their environment by sending their own. This striking enrichment of cell-signaling genes in plastic cell populations, in addition to the rapid and stereotypical remodeling of the tissue environment, prompted us to investigate how cell-cell communication may drive new cell-states. Computational approaches to study cell-cell communication typically score cognate R-L gene pairs one at a time, and they therefore failed to find robust and specific signals among the large number of cytokines and receptors expressed across cell populations. We used Calligraphy, a method that uses modularity in gene expression rather than individual R-L pairs to gain robustness, to focus on genes particularly influenced by perturbation, and to predict cell-cell communication networks unique to the neoplastic state. As desired, Calligraphy captured a clear modularity of communication genes in our data, consistently identifying modules of co-expressed communication genes that showed a one-to-one mapping to transcriptional cell-states. These data imply that communication is critically important to establishing cell-state diversity within the pre-malignant pancreas.

The communication genes involved in signaling networks identified in the pre-malignant pancreas were markedly absent from those expressed in normal regeneration. In the latter context, we found a small, transient subpopulation of cells with an active *Kras* signature that express the Gastric communication module. The Gastric module has the highest propensity for tissue remodeling, demonstrating that cancer commandeers gene programs from normal regeneration. Moreover, the Gastric, Progenitor and Bridge communication modules persist in advanced murine and human cancers, implying that these programs are important for maintaining disease. Our results go beyond producing an atlas of early or late neoplasia (*32, 78– 80*) to demonstrate an epigenetic basis for cell-state diversity, with an output of that diversity being novel cell-cell interactions that drive further diversity and distinguish normal regeneration from cancer.

We reasoned that neoplasia-specific tissue remodeling is likely amplified by feedback between epithelial and immune compartments. Indeed, our analyses revealed a feedback loop initiated by IL-33 signaling from epithelial cells expressing the Gastric module to Th2 cytokine -expressing Tregs and ILCs, which signal back to the epithelium (among multiple other routes). Furthermore, we used a new mouse model to experimentally validate the profound consequences of this signaling loop on the neoplastic remodeling of the pre-malignant epithelium. These findings link previous results on the relevance of Th2 cytokine expression in PDAC (*81*) to an IL-33-driven feedback loop. Spatial analysis revealed co-localization of these signaling populations, and *in vivo* perturbation using a newly constructed GEMM showed that knockdown of *Il33* in the epithelium blocks cells in a highly plastic progenitor state. Empirical results of *Il33* perturbation match Calligraphy’s predictions with remarkable precision in terms of which populations are perturbed, to what degree, and in what temporal sequence. We first observe remodeling of the immune compartment (2 days), which is a direct IL-33 target, followed by extensive remodeling of the epithelium (3 weeks). The highly plastic *Nes*^+^ progenitor population that uniquely emerges in the context of oncogenic *Kras* is blocked from progression; thus, the gastric populations normally abundant in PanINs do not emerge at high frequency and are depleted. While we and others previously showed that IL-33 plays an important role in early tumorigenesis (*25*) in addition to advanced PDAC (*71*), our work here allowed us to dissect its profound impact by mapping the mechanisms and the striking number of cell populations on which it acts. Interestingly, we also show that this mechanism can be uniquely driven by epithelial-derived IL-33, despite the high degree of stromal IL-33 expression and previous reports implicating fibroblast-derived IL33 in disease phenotypes (*82–84*).

Pancreas cancer is a lethal disease that is frequently detected too late for therapeutic intervention and, as such, a detailed understanding of early neoplastic events may enable the development of rational strategies to prevent, detect and intercept tumors before they evolve to an intractable stage. Nonetheless, the development of such strategies has been hampered by our incomplete understanding of the earliest stages of neoplasia, a setting that remains difficult to access and study in the human disease. Our results reveal cellular and molecular details whereby epigenetic plasticity established cell-states and signaling networks important for neoplasia that are not observed in the normal or regenerative pancreas. Efforts to further understand the mechanisms dictating neoplasia-specific cell-cell communication networks driving tumor initiation holds promise for the development of therapeutics that block early cancer progression and may also be effective against advanced disease.

## Acknowledgments

We thank J. Simon and the MSKCC animal facility for technical support with animal colonies; MSKCC Flow Cytometry facility for support with FACS sorting experiments; the Single Cell and Imaging Mass Cytometry Platform at Goodman Cancer Research Centre; the members of the Sloan Kettering Institute’s Single Cell Analytics and Innovation Lab (SAIL) computational unit, and members of the Lowe and Pe’er laboratories for advice and discussions.

## Author contributions

**Cassandra Burdziak**: Conceptualization, Methodology, Software, Formal analysis, Data curation, Investigation, Writing-original draft presentation, Visualization, Funding Acquisition; **Direna Alonso-Curbelo**: Conceptualization, Methodology, Data curation, Investigation, Writing-original draft presentation, Visualization, Funding Acquisition; **Thomas Walle**: Formal analysis, Data curation, Investigation, Writing-review and editing, Visualization; **Francisco M. Barriga**: Investigation; **José Reyes**: Data curation, Investigation, Writing-review and editing; **Yubin Xie**: Formal analysis, Data curation, Investigation; **Zhen Zhao**: Investigation; **Chujun Julia Zhao**: Formal analysis; **Hsuan-An Chen**: Investigation; **Ojasvi Chaudhary**: Investigation; **Ignas Masilionis**: Investigation; **Zi-Ning Choo**: Resources; **Vianne Gao**: Data curation; **Wei Luan**: Investigation; **Alexandra Wuest**: Investigation; **Yu-Jui Ho**: Data curation; **Yuhong Wei**: Resources; **Daniela Quail**: Resources; **Richard Koche**: Formal analysis; **Linas Mazutis**: Investigation; **Tal Nawy**: Writing-original draft presentation; **Ronan Chaligné**: Investigation; **Scott W. Lowe**: Conceptualization, Methodology, Writing-original draft presentation, Funding Acquisition, Study supervision; **Dana Pe’er**: Conceptualization, Methodology, Writing-original draft presentation, Funding Acquisition, Study supervision

## Data and materials availability

All sequencing data have been deposited at the Gene Expression Omnibus (GEO). Code for data analysis is available at https://github.com/dpeerlab/pdac-progression.

## Supplementary Materials

Materials and Methods

Figure S1 (Related to figure 1)

Figure S2 (Related to figure 1)

Figure S3 (Related to figure 2)

Figure S4 (Related to figure 3)

Figure S5 (Related to figure 3)

Figure S6 (Related to figure 4)

Figure S7 (Related to figure 4)

Figure S8 (Related to figure 4)

Figure S9 (Related to figure 4)

Figure S10 (Related to figure 5)

Figure S11 (Related to figure 6)

Table S1 Sample and GEMM Information

Table S2 Benign and Malignant Signature Genes

Table S3 Plasticity Score Gene Correlations

Table S4 Calligraphy Communication Modules

Table S5 Calligraphy Module-Module Interactions

Table S6 GEMM Alleles (Materials and methods)

Table S7 Experimental Conditions (Materials and methods)

Table S8 IMC Panel (Materials and methods)

Table S9 scRNA-seq Cohorts and QC (Materials and methods)

Table S10 scRNA-seq Epithelial Annotation Strategy (Materials and methods)

Table S11 Epithelial scRNA-seq Analysis Groups (Materials and methods)

Table S12 scRNA-seq Immune Genes (Materials and methods)

Table S13 Immune scRNA-seq Analysis Groups (Materials and methods)

Table S14 scATAC-seq Sample Pairings (Materials and methods)

Table S15 scATAC-seq Annotation Strategy (Materials and methods)

Tables S16 Cognate R-L Pairs (Materials and methods)

### Materials and Methods

#### Mouse models

All animal experiments in this study were performed in accordance with protocols approved by the Memorial Sloan Kettering Institutional Animal Care and Use Committee. Mice were maintained under specific pathogen-free conditions, and food and water were provided ad libitum. In all experiments with PDAC models, tumors did not exceed a maximum volume corresponding to 10% of the mouse’s body weight (typically 12 mm diameter). Mice were evaluated daily for signs of distress or end-point criteria. Specifically, mice were immediately euthanized if they presented signs of cachexia, weight loss of more than 20% of initial weight or breathing difficulties, or if they developed tumors of 12 mm in diameter. No tumors exceeded this limit.

##### Generation and authentication of KC-shIl33 embryonic stem cell clones

KC-sh*Il33* mouse embryonic stem (mES) cells were generated by targeting established KC embryonic stem (ES) cells (*Ptf1a*-*cre*;*LSL-Kras^G12D^;RIK;CHC* (*91*)) with two independent GFP-linked *Il33* shRNAs (shIl33.668 and shIl33.327) cloned into miRE-based targeting constructs (*92*), as previously described (*91, 93*). Targeted ES cells were selected and functionally tested for single integration of the GFP-linked shRNA element into the *CHC* locus as previously described (*93*). Before injection, ES cells were cultured briefly for expansion in KOSR+2i medium (*94*). Two clones (KC-shIl33.668 clone 3 and KC-shIl33.327 clone 2) were used for cohort generation, single-cell omics and phenotypic analyses. The identity and genotype of the ES cells, resulting chimaeric mice and their progeny were authenticated by genomic PCR using a common Col1a1 primer (5′-CACCCTGAAAACTTTGCCCC-3′) paired with a transgene specific primer: shRen.713: 5′-GTATAGATAAGCATTATAATTCCTA-3′; shIl33.668: 5′-TTCAAATGAAACAAAGTCC-3′; shIl33.327: 5′-TTAAAAGTGAAGTTCCTTGGA-3′, yielding a product of around ∼250-bp. ES cells were confirmed to be negative for mycoplasma and other microorganisms before injection.

##### Mouse alleles

All alleles have been previously described. *Ptf1a-cre* (*27*), *LSL-Kras^G12D^* (*95*), *p53^fl^* (*31, 85*)*, CHC* (*96*), and *LSL-rtTA3-IRES-mKate2 (RIK)* (*97*) and *TRE-GFP-shRen* (*98*) strains were interbred and maintained on mixed Bl6/129J backgrounds. Combinations of these alleles enable selective isolation of epithelial cells from pancreatic tissues. Specifically, we used the pancreas-specific Cre driver *Ptf1a*-*cre* and the lineage-tracing allele *RIK* that, by themselves or in combination with a Cre-activatable *Kras^G12D^* allele, enable tagging of pancreatic epithelial cells that contain wild-type or mutant *Kras* by the fluorescent reporter mKate2. The following table summarizes the nomenclature used for multiallelic strains used in this work.

**Table S6:** GEMM Alleles.

| Short name | Alleles |
| --- | --- |
| C | *Ptfla-cre;RIK* (C;RIK) |
| KC | *Ptfla-cre;RIK;LSL-Kras<sup>G12D</sup>* (KC;RIK) |
| KPC | *Ptfla-cre;RIK;LSL-Kras<sup>G12D</sup>;p53<sup>fl/+</sup> Ptfla-cre;RIK;LSL-Kras<sup>G12D</sup>;p53<sup>R172H/+</sup>* (KPC;RIK) |
| KC-shRen | *Ptfla-cre;RIK;LSL-Kras<sup>G12D</sup>;TRE-GFP-shRen.713* (KC;RIK-shRen.713) |
| KC-shI133 | *Ptfla-cre;RIK;LSL-Kras<sup>G12D</sup>;TRE-GFP-shI133.668*<br>*Ptfla-cre;RIK;LSL-Kras<sup>G12D</sup>;TRE-GFP-shI133.327*<br>(KC;RIK-shI133.668 or KC;RIK-shI133.327) |

##### Cohort generation

C, KC and KPC mice were generated by strain intercrossing. To generate KC-shRen and KC-shIl33 mice, chimaeric cohorts of male mice derived from the ES cells described above were generated by the Center for Pancreatic Cancer Research (CPCR) at Memorial Sloan Kettering Cancer Center (MSKCC) or the Rodent Genetic Engineering Core at New York University as previously described (*91*). Only mice with a coat color chimaerism of over 95% were included for experiments.

##### Acute pancreatitis

To compare the effects of tissue injury in the transcriptional and chromatin accessibility landscapes of mutant-*Kras*-and wild-type-*Kras*-expressing pancreatic epithelial cells, C, KC, KC-sh*Ren* or KC-sh*Il33* 5-week-old male mice were treated with eight-hourly intraperitoneal injections of 80 μg per kg of caerulein (Bachem) or PBS for two consecutive days.

##### Il33 perturbation

For induction of shRNA expression, KC-sh*Ren* or KC-sh*Il33* mice were switched to a doxycycline diet (625 mg per kg, Harlan Teklad) that was changed twice weekly at 4 weeks of age to induce shRNA expression, and were subsequently treated with caerulein (acute pancreatitis protocol) 6 days thereafter to study contribution to cell-cell networks and tissue phenotype during injury (pancreatitis)-accelerated neoplasia.

##### Sample collection timing

Samples were collected to span the entire range of PDAC progression, from initiation to metastasis. In malignant tissue states (K5, K6), PDAC cells were isolated from primary or metastatic cancer lesions arising in autochthonous transgenic models (KPC) that were macro-dissected away from premalignant or normal tissue. The following table summarizes the conditions and timepoints of sample collection.

**Table S7:**
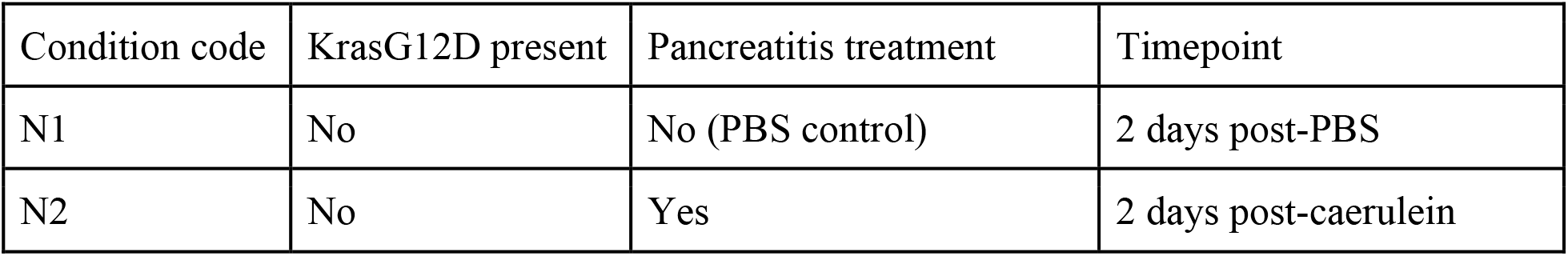
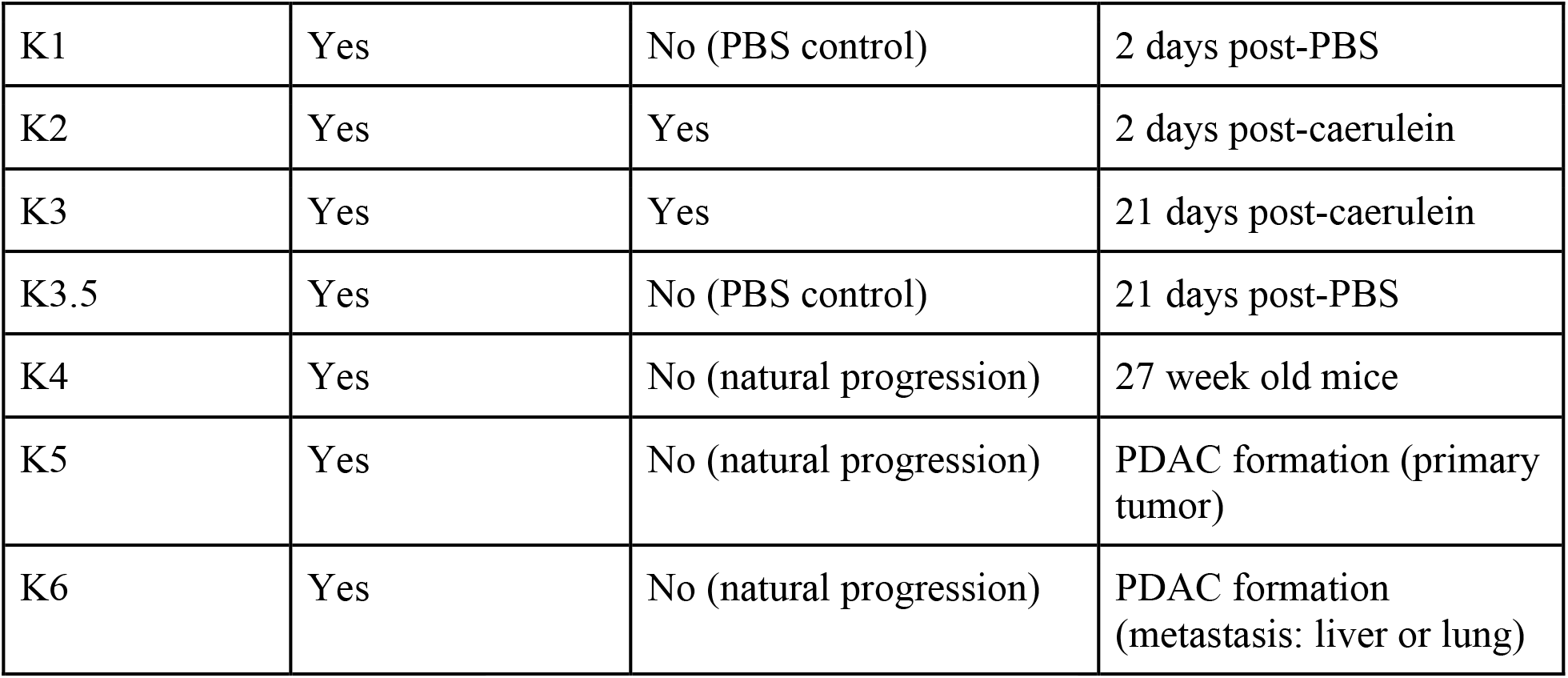
Experimental conditions.

| Condition code | KrasG12D present | Pancreatitis treatment | Timepoint |
| --- | --- | --- | --- |
| N1 | No | No (PBS control) | 2 days post-PBS |
| N2 | No | Yes | 2 days post-caerulein |
| K1 | Yes | No (PBS control) | 2 days post-PBS |
| K2 | Yes | Yes | 2 days post-caerulein |
| K3 | Yes | Yes | 21 days post-caerulein |
| K3.5 | Yes | No (PBS control) | 21 days post-PBS |
| K4 | Yes | No (natural progression) | 27 week old mice |
| K5 | Yes | No (natural progression) | PDAC formation (primary tumor) |
| K6 | Yes | No (natural progression) | PDAC formation (metastasis: liver or lung) |

#### Immunofluorescence, IHC and histological analyses

Tissues were fixed overnight in 10% neutral buffered formalin (Richard-Allan Scientific), embedded in paraffin, cut into 5-μm sections and, immunofluorescence, immunohistochemical or H&E stainings were performed following standard protocols, as previously described (*25*). The following antibodies were used: mKate2 (AB233, Evrogen), CD45 (ab25386, Abcam), TFF1 (TA322883, Cedarlane/OriGene Technologies), GFP (ab13970, Abcam), IL-33 (AF3626, R&D). CD45 Immunohistochemistry was performed on a Bond Rx autostainer (Leica Biosystems) with Histowiz. H&E and IHC slide scanning was performed with Histowiz. IF images were acquired on a Zeiss AxioImager microscope using using a 10× (Zeiss NA 0.3) or 20× (Zeiss NA 0.17) objective, an ORCA/ER CCD camera (Hamamatsu Photonics), and Axiovision or Zeiss (ZEN 2.3) software.

#### IMC collection and analysis

Antibodies were optimized via immunofluorescence and conjugations were carried out in house and by the Single Cell and Imaging Mass Cytometry Platform at the Goodman Cancer Research Centre (McGill University), using Maxpar Conjugation Kits (Fluidigm), according to manufacturer’s instructions. Deparaffinization and heat-induced epitope retrieval were performed using the Ventana Discovery Ultra auto-stainer platform (Roche Diagnostics). FFPE slides were incubated in EZ Prep solution (preformulated, Roche Diagnostics) at 70 °C to deparaffinize, followed by antigen-retrieval in standard Cell Conditioning 1 solution (CC1, preformulated; Roche Diagnostics) at 95 °C. Slides were then washed in 1× PBS, blocked in Dako Serum-free Protein Block solution (Agilent), followed by antibody staining overnight at 4 °C as described by Fluidigm for FFPE tissues. Tissues were stained with a panel of multiplexed metal-conjugated antibodies (**Table S8** specifies the antibodies and metals shown in **Fig. 6 and S11**). IMC images were acquired at a resolution of roughly 1 μm, frequency of 200 Hz and area of 1 mm2, Hyperion Imaging System and CyTOF Software v7.0.8493.0. (Fluidigm).

**Table S8:** IMC Metal-conjugated primary antibodies.

| Antibody | Company | Catalog# | Metal |
| --- | --- | --- | --- |
| mKate2 | Evrogen | AB233 | 142Nd |
| Vimentin | Fluidigm | 3143027D | 143Nd |
| IL-33 | R&D systems | AF3626 | 152Sm |
| ANXA10 | Abcam | ab223131 | 154Sm |
| FoxP3 | Cell Signaling Technology | 12653 | 158Gd |
| aSMA | Abcam | ab5694 | 175Lu |
| GFP | Abcam | ab220802 | 176Yb |
| DNA1 | Fluidigm | 201192B | 191Ir |
| DNA2 | Fluidigm | 201192B | 193Ir |

Sliding windows were used to extract regional information from the Imaging Mass Cytometry image data. The size of the window was 30×30 pixels. The images were padded with 0 values to be exact division of the window size. The average value of a maker is used to represent the marker expression in the window (**Fig. S11A,B**).

#### Tissue dissociation and single cell analyses (scRNA-seq, scATAC-seq, bulk ATAC-seq)

For scRNA-seq and scATAC-seq collection, lineage-traced (mKate2+) epithelial cells or CD45+ immune cells were freshly isolated from pancreatic tissues from C, KC, KPC, or KC-shRNA mice by FACS sorting. Specifically, pancreases were finely chopped with scissors and incubated with digestion buffer containing 1 mg ml^−1^ collagenase V (C9263, Sigma-Aldrich), 2 U ml^−1^ dispase (17105041, Life Technologies) dissolved in HBSS with Mg^2+^ and Ca^2+^ (14025076, Thermo Fisher Scientific) supplemented with 0.1 mg ml^−1^ DNase I (Sigma, DN25-100MG) and 0.1 mg ml^−1^ soybean trypsin inhibitor (STI) (T9003, Sigma), in gentleMACS C Tubes (Miltenyi Biotec) for 42 min at 37 °C using the gentleMACS Octo Dissociator. Normal (non-fibrotic) pancreas samples were dissociated as above, except that the digestion buffer contained 1 mg ml^−1^ collagenase D (11088858001, Sigma-Aldrich) instead of collagenase V. After enzymatic dissociation, samples were washed with PBS and further digested with a 0.05% solution of Trypsin-EDTA (15400054, Thermo Fisher Scientific) diluted in PBS for 5 min at 37 °C. Trypsin digestion was neutralized with FACS buffer (10 mM EGTA and 2% FBS in PBS) containing DNase I and STI. Samples were then washed in FACS buffer containing DNase I and STI, and filtered through a 100-μm strainer. Cell suspensions were blocked for 5 min at room temperature with rat anti-mouse CD16/CD32 with Fcblock (Clone 2.4G2, BD Biosciences) in FACS buffer containing DNase I and STI, and an APC-conjugated CD45 antibody (Clone 30-F11, Biolegend, 1:200) or APC-Cy7 CD45 antibody (Clone 30-F11, Biolegend, 1:200) was then added and incubated for 10 min at 4 °C. Cells were then washed once in FACS buffer containing DNase I and STI, filtered through a 40-μm strainer and resuspended in FACS buffer containing DNase I and STI and 300 nM DAPI as a live-cell marker. Sorts were performed on a BD FACSAria I or BD FACSAria III cell sorters (Becton Dickinson) for mKate2 (co-expressing GFP for on-doxycycline shRNA mice), excluding DAPI^+^ and CD45^+^ cells. Cells collected in 2% FBS in PBS. For single-cell analyses, single-cell suspensions were spin down and resuspended in 0.05% BSA containing 1 U/µL Invitrogen Ambion RNase Inhibitor (AM2682, Thermo Fisher Scientific).

#### Immunophenotyping

Collection of immune cells for immunophenotyping followed the dissociation protocol used to isolate epithelial cells, with the following differences: (i) digestion buffer did not include dispase, and (ii) trypsin digestion step was not performed, to optimally preserve surface epitopes. For multi-parametric flow cytometry analysis, cell suspensions were stained with LIVE/DEAD fixable viability dye (Invitrogen) for 30 min at 4 °C. After this, cells were washed, incubated with Fc block (BD, 1:200) for 15 min at 4 °C, and then stained with a cocktail of conjugated antibodies for 30 min at 4°C. After staining cells were washed and fixed (BD Cytofix) for 20 min at 4C, washed again, and stored for analysis. Samples were analyzed in a BD LSRFortessa with 5 lasers, where gates were set by use of fluorescence-minus-one (FMO) controls. The following antibodies were used to quantify the fraction and identity of IL1RL1 ST2 receptor expressing cells: AF700 CD45 (clone 30-F11 BioLegend #103128), BUV395 CD11b (clone M1/70 BD Biosciences #563553), PerCP Cy5.5 Nkp46 (clone 29A1.4 BioLegend #137609), PE eFluor 610 CD3e (clone 145-2C11 eBioscience #61-0031-82), PE Cy7 CD8 (clone 53-6.7 BioLegend #100722), Strep BUV661 (BD Biosciences #612979), biotin-ST2 (clone RMST2-33 eBioscience #13-9333-82), BV650 CD19 (clone 1D3 BD Biosciences #563235), APC-Cy7 Gr1 (clone RB6-8C5 BioLegend #108424), BV605 CD4 (clone RM4.5 BD Biosciences #563151), APC F4/80 (clone BM8 BioLegend #123116), BV785 CD44 (IM7 Biolegend #103041), BV711 EpCAM (clone G8.8 BioLegend #118233), PE CD11c (clone N418 BioLegend #117307), FITC MHCII (clone M5/114.15.2 Thermo Scientific #11-5321-82).

#### Encapsulation and sequencing of scRNA-seq samples

Cells were resuspended in 1X PBS and BSA (0.04%) and checked for viability using 0.2% (w/v) Trypan Blue staining (Countess II). All sequencing experiments were performed on samples with a minimum of 80% viable cells. Single-cell encapsulation and scRNA-seq library prep of FACS-sorted cell suspensions was performed on the Chromium instrument (10x Genomics) following the user manual (Reagent Kit 3’ v2). Each sample loaded onto the cartridge contained approximately 5,000 cells at a final dilution of ∼500 cells/μl. Transcriptomes of encapsulated cells were barcoded during reverse transcription and the resulting cDNA was purified with DynaBeads, followed by amplification per the user manual. Next, the PCR-amplified product was fragmented, A-tailed, purified with 1.2X SPRI beads, ligated to sequencing adapters and indexed by PCR. Indexed DNA libraries were double-size purified (0.6–0.8X) with SPRI beads and sequenced on an Illumina sequencer (R1 – 26 cycles, i7 – 8 cycles, R2 – 70 cycles or higher) to a depth of >50 million reads per sample (>13,000 reads/cell) at MSKCC’s Integrated Genomics Operation Core Facility.

#### Encapsulation and sequencing of scATAC-seq samples

Approximately 50,000 mKate2^+^ epithelial (mKate2^+^;CD45^−^;DAPI^−^) or CD45+ cells were isolated from premalignant pancreata by FACS and subjected to scATAC-seq protocol (10X Genomics, CG000168 RevA) (*99*). In brief, FACS-sorted cells were lysed in cold lysis buffer (0.1% NP-40, 0.1% Tween 20, 0.01% digitonin, 10 mM NaCl, 3 mM MgCl_2_ and 10 mM Tris-HCl (pH 7.4)), washed and processed according to ‘Nuclei Isolation for Single-Cell ATAC Sequencing’ protocol (CG000169 RevD), according to the manufacturer’s instructions. The resulting nuclei suspension was subjected to transposition reaction for 60 min at 37 °C and then encapsulated in microfluidic droplets using a 10X Chromium instrument following the manufacturer’s instructions with a targeted nuclei recovery of approximately 5,000. Barcoded DNA material was cleaned and prepared for sequencing according to the Chromium Single Cell ATAC Reagent Kits User Guide (10X Genomics; CG000168 RevA). Purified libraries were assessed using a Bioanalyzer High-Sensitivity DNA Analysis kit (Agilent) and sequenced on an Illumina platform at approximately 150 million reads (R1 50 bp, R2 50 bp, i7 8 bp, i5 16 bp) per 1 sample (around 5,000 nuclei) at MSKCC’s Integrated Genomics Operation Core.

#### Single-cell RNA-seq data processing and basic analysis

*Custom cluster-based filtering pipeline:* All scRNA-seq datasets were initially processed (de-multiplexed, barcode-corrected, aligned, UMI-corrected) with SEQC (*100*) using mouse genome mm10 and default parameters for the v2 3’ scRNA-seq kit. After constructing a preliminary molecule count matrix from all barcodes, SEQC filters empty drops and poor quality cells using four main criteria: cell library sizes (total transcript counts), cell coverage (average reads per molecule), mitochondrial (MT) RNA content reflecting dead or dying cells, and library complexity (number of unique genes captured by library size). However, we find that scRNA-seq data from pancreas are prone to increased technical noise, including higher MT RNA expression; thus, we opted to bypass SEQC’s automated filtering to ensure our analysis did not inadvertently discard relevant cells. Instead, we developed a manual, cluster-based filtering pipeline that relies on the metrics listed above plus additional inclusion criteria for calling real, high-quality cell populations. The guiding principle of this pipeline is that low quality or dying cells tend to form distinct phenotypic clusters with poor technical characteristics, which can be removed by iterative rounds of clustering and filtering. Manual filtering also has the advantage of allowing system-specific biological knowledge to guide filtering decisions, which is especially important when rare populations may be confounded with technical features of the data (e.g. smaller cells correlate with low library sizes, or specialized cell types express a restricted set of genes, resulting in low library complexity).

We first performed a liberal per-sample filtering step to remove obvious empty drops. Specifically, we retained the 15,000 barcodes with highest total transcript counts (∼5,000 cells were loaded into the lane), a permissive threshold that avoids excluding real cells with lower library size. We then clustered our data with Phenograph (*101*) and nominated clusters potentially comprising empty drops or dying cells based on one or more SEQC metrics of coverage, MT content and library complexity. To further assess nominated low-quality clusters, we constructed per-cluster gene-gene correlation matrices and inspected the matrix block structure, which represents co-regulated gene modules. We reason that the tight regulatory programs driving biological functions are loosened in dying cells, which should be reflected in a breakdown of modularity in gene-gene co-expression matrices. To suggest potential real populations, we also looked for coherent expression of pancreas or immune marker gene programs in each population. Finally, we removed clusters that 1) had substantially poorer-than-average SEQC metrics, and/or 2) lacked co-regulated gene modules and evidence of coherently expressed gene programs. We retained any clusters for which there was uncertainty.

After filtering each sample individually, we pooled sets of samples to increase our power to detect rare populations, and repeated clustering and filtering iteratively until only high-quality cells remained. Following the iterative filtering of pooled sets, filtered count matrices were generated independently for each sample. To guide pooling and downstream analyses, samples were grouped into three pre-defined cohorts which address questions associated with (1) full tumor progression (“Progression Cohort”), (2) the impacts of *Il33* perturbation on particular stages of progression (“Perturbation Cohort”), and (3) involvement of the immune microenvironment in both of these cases (“Immune Cohort”) (see **Table S9** for assigned cohorts, filtering groups and filtered count matrix statistics).

**Table S9.** scRNA-seq Cohorts and QC. Samples were assigned to one of three cohorts addressing broad questions, as well as groups for pooling prior to cluster-based filtering. Cell number and mean library size reflect the final filtered count matrix for each sample.

| Cohort | Sample ID | Stage | Genotype | Filtering Group | No. of Cells | Median Library Size |
| --- | --- | --- | --- | --- | --- | --- |
| Progression | DACD511_Kate_plus | N1 | C;RIK | 1 | 2491 | 4534 |
| Progression | DACD550_kate_plus | N1 | C;RIK | 1 | 2890 | 5855 |
| Progression | DAC_B530-Kate+ | N2 | C;RIK | 2 | 2804 | 5610.5 |
| Progression | DACD403_Kate_plus | N2 | C;RIK | 2 | 1424 | 5309.5 |
| Progression | DACD394_Kate_plus | K1 | KC;RIK-shRen.713 (Off Dox) | 2 | 2834 | 7562.5 |
| Progression | DACD406_Kate_plus | K1 | KC;RIK | 2 | 2189 | 10276 |
| Progression | DACD351-Kate+ | K1 | KC;RIK-shRNA (Off Dox) | 2 | 2781 | 17184 |
| Progression | DAC_C263_EPI | K1.5 | KC;RIK-shRNA (Off Dox) | 2 | 1851 | 14922 |
| Progression | D396_EPI | K1.5 | KC;RIK-shRNA (Off Dox) | 2 | 2776 | 1732.5 |
| Progression | DACD404_Kate_plus | K2 | KC;RIK | 2 | 1595 | 10470 |
| Progression | DACD407_Kate_plus | K2-Day 1 | KC;RIK | 2 | 2075 | 12076 |
| Progression | DAC_DI143_Epi | K3 | KC;RIK | 2 | 2423 | 15665 |
| Progression | DACD482_Kate_plus | K3 | KC;RIK | 2 | 2668 | 10252.5 |
| Progression | DAC_C301-EPI_1 | K4 | KC;RIK | 2 | 1812 | 2433.5 |
| Progression | DAC_C301-EPI_2 | K4 | KC;RIK | 2 | 1142 | 615.5 |
| Progression | Ag-PDAC-PT-Kate | K5 | KPC;RIK (KPR172HC;RIK) | 2 | 3387 | 15828 |
| Progression | Ag-Lung-Mets-Kate | K6 | KPC;RK (KPR172HC;RIK) | 3 | 389 | 40691 |
| Progression | DAC_D020_p5_Epi | K5 | KPflC;RIK | 2 | 3646 | 22107 |
| Progression | DACC963PT_Kate_plus | K5 | KPflC;RIK | 2 | 679 | 5941 |
| Progression | DACC963LIVERmet | K6 | KPflC;RIK | 2 | 2244 | 30530 |
| Progression | DACC963_mKate_plus | K6 | KPflC;RIK | 2 | 1863 | 30134 |
| Perturbation | DACD350-Kate+ | K2 Control | KC;RIK-shII33.668 (Off Dox) | 2 | 3088 | 15149.5 |
| Perturbation | DACE621-mKate2 | K2 Control | KC;RIK-shRen.713 (On Dox) | 4 | 2670 | 18234 |
| Perturbation | DACD346-Kate+ | K2 + IL33 KD | KC;RIK-shII33.668 (On Dox) | 2 | 3067 | 16170 |
| Perturbation | DACE604-mKate2 | K2 + IL33 KD | KC;RIK-shII33.668 (On Dox) | 4 | 3178 | 16897.5 |
| Perturbation | DACE605-mKate2 | K2 + IL33 KD | KC;RIK-shII33.668 (On Dox) | 4 | 2792 | 18296 |
| Perturbation | DACD349_mKate2+;GFP+ | K3 Control | KC;RIK-shII33.668 (Off Dox) | 2 | 903 | 18997 |
| Perturbation | DACE610-EPI | K3 Control | KC;RIK-shII33.327 (Off Do) | 4 | 1053 | 20917 |
| Perturbation | DACD347_mKate2+;GFP+ | K3 + IL33 KD | KC;RIK-shII33.668 (On Dox) | 2 | 1049 | 18285 |
| Perturbation | DACE607-EPI | K3 + IL33 KD | KC;RIK-shII33.668 (On Dox) | 4 | 2082 | 15419.5 |
| Perturbation | DACE614-EPI | K3 + IL33 KD | KC;RIK-shII33.327-(OnDox) | 4 | 1255 | 16008 |
| Immune | DACD407_CD45+_Day1 | K2-Day 1 | KC;RIK | 5 | 1937 | 2007 |
| Immune | DACD404_CD45 | K2 | KC;RIK | 5 | 918 | 4243.5 |
| Immune | DACD143_CD45+ | K3 | KC;RIK | 5 | 1825 | 1752 |
| Immune | DACD482_CD45+ | K3 | KC;RIK | 5 | 2186 | 2779.5 |
| Immune | DACD406-CD45+ | K1 | KC;RIK | 5 | 2250 | 5860 |
| Immune | DACD408-CD45+ | K1 | KC;RIK | 5 | 2167 | 2465 |
| Immune | DACD351-CD45+ | K1 | KC;RIK-shII33.668 (Off Dox) | 5 | 2033 | 8389 |
| Immune | DACD350-CD45+ | K2 Control | KC;RIK-shII33.668 (Off Dox) | 5 | 2414 | 1752 |
| Immune | DACE621-CD45 | K2 Control | KC;RIK-shRen.713 (On Dox) | 6 | 3272 | 1724 |
| Immune | DACD346-CD45+ | K2 + IL33 KD | KC;RIK-shIl33.668 (On Dox) | 5 | 2047 | 8017 |
| Immune | DACE604-CD45 | K2 + IL33 KD | KC;RIK-shIl33.668 (On Dox) | 6 | 2068 | 1802.5 |
| Immune | DACE605-CD45 | K2 + IL33 KD | KC;RIK-shIl33.668 (On Dox) | 6 | 2469 | 7197 |
| Immune | DACD349_CD45+ | K3 Control | KC;RIK-shIl33.668 (Off Dox) | 5 | 705 | 3876 |
| Immune | DACE610-CD45 | K3 Control | KC;RIK-shIl33.668 (Off Dox) | 6 | 1555 | 3222 |
| Immune | DACD345_CD45+ | K3 + IL33 KD | KC;RIK-shIl33.668 (On Dox) | 5 | 228 | 3085 |
| Immune | DACD347_CD45+ | K3 + IL33 KD | KC;RIK-shIl33.668 (On Dox) | 5 | 1110 | 3320 |
| Immune | DACE607-CD45 | K3 + IL33 KD | KC;RIK-shIl33.668 (On Dox) | 6 | 1496 | 5020 |
| Immune | DACE614-CD45 | K3 + IL33 KD | KC;RIK-shIl33.327 (On Dox) | 6 | 1908 | 3807 |

The following sections describe the cohort-specific analysis of filtered count matrices, including data integration, normalization, dimensionality reduction, and visualization:

##### Progression Cohort (N1–K6 epithelial cells, sorted based on mKate+)

The Progression Cohort (**Table S9**) contains all epithelial datasets from normal and regenerating pancreata (N1, N2), *Kras*-mutant pre-malignant pancreata with and without injury (K1–K4), and *Kras/p53*-altered malignant tumors (K5, K6), forming the basis for our investigation of tumor progression at various scales. All filtered datasets from this cohort were combined into a single data matrix without additional data harmonization, given the high degree of similarity and overlap between our biological replicates (**Fig. S2E**), thus allowing us to preserve the original counts. Standard log library size normalization was applied to the combined data using a scaling factor of 10,000 in lieu of median library size. We removed ribosomal genes, MT genes, *Malat1*, and genes expressed in fewer than 20 cells, resulting in a count matrix containing 16,828 genes.

An additional round of filtering was performed to remove low-quality cells which do not form separate clusters and hence passed the custom filtering described above. We inspected the log-library-size distribution of each sample separately and filtered cells below a manually determined threshold for the lower mode. By performing cluster-level analysis first (allowing us to detect populations that can only be discovered upon sample aggregation), we could ensure that entire cell types were not lost in subsequent cell-level filtering of each sample by library size. Finally, we clustered cells with Phenograph and removed 3 remaining outlier clusters with very low library sizes, mainly consisting of acinar cells from the basal pancreas (N1) condition. The resulting dataset comprised 30,661 high-quality cells.

After removing putative empty drops and low-quality cells, we ran DoubletDetection (*102*) for each sample individually with default parameters. Given that we expected this data to contain cells residing along continuous trajectories, we were conservative in doublet filtering so as not to remove false positives arising from true intermediate cell states. Hence, we removed only discrete Phenograph clusters (k = 40, computed on all samples in the combined count matrix) which contained a high fraction (>15%) of doublets, for a total of 214 cells across samples.

Finally, we removed clusters that express mesenchymal markers typically associated with fibroblasts (*Acta2, Col1a1, Lum*), representing potential stromal contaminants. Four clearly stroma-like clusters were filtered, resulting in a final dataset of 28,131 high-confidence mKate2+ epithelial cells.

Next, highly variable gene (HVG) selection was performed in a similar fashion to (*103*). Briefly, for each gene, we computed the mean and standard deviation of normalized (unlogged) expression across all cells, then fit a Lowess regression model to the trend of log(coefficients of variation, pseudocount = 0.1) vs. log(mean expression). HVGs were identified by coefficients of variation that substantially exceed mean expression according to this trend, generating high residuals in the regression. To avoid biasing toward highly expressed genes, we binned genes by mean expression (40 total bins of expressed genes) and selected the top 200 with high residuals in each bin, resulting in a total of 8,000 genes for downstream analysis. We included a large number of HVGs for Progression Cohort analysis to ensure that we retained important pancreas and cancer-associated genes and captured the spectrum of normal and disease states. To analyze subsets of this cohort, including specific stages or clusters, we chose fewer HVGs to focus on variability specific to that set (see **Table S11** for description of each analysis and parameters).

Next, we performed principal component analysis (PCA) on the filtered, log-normalized Progression Cohort matrix restricted to 8,000 HVGs. We used automatic knee point detection based on the cumulative variance explained per component to determine a cutoff for the number of PCs to carry forward, selecting 49 top PCs which explained ∼41% of the variance. We further verified that adding subsequent PCs has trivial impact on the overall variance explained (i.e. inclusion of 50 additional PCs explains <3% further variance). To visualize the full map in 2 dimensions, we ran tSNE using the bhtSNE package (*104*) on selected PCs with perplexity = 150 and theta = 0.5. We then ran diffusion maps and selected 13 significant diffusion components (DCs) via Eigen gap to obtain a non-linear embedding of cells for downstream analysis.

To assess whether the DCs represent known PDAC biology, we examined how published signatures of normal regeneration and disease states were distributed across DCs. We obtained published gene sets derived from bulk RNA-seq of human tumors and matched normal samples (Differentially expressed genes (DEGs) between Human PDAC vs Human Normal samples determined by > 2-fold change in gene expression with adjusted P-value < 0.05; (*36, 37*)), as well as complementary mouse models (DEGs between murine PDAC vs Normal, determined by > 2-fold change in gene expression with adjusted P-value < 0.05; (*25*)). For human gene sets, we mapped orthologous gene names to obtain equivalent mouse gene sets with Ensembl annotations. We then obtained z-scores for each gene to scale log-normalized features for comparison, and computed the average z-score across all captured genes per set. We visualized these scores for each gene set and sorted cells by DC 1, which roughly captures the order of progression determined from our scRNA-seq data (**Fig. 1C**). This visualization supports that diffusion maps derived from our unbiased Progression Cohort analysis effectively capture biologically meaningful axes of disease state variation.

Our next step was to define discrete cell types within the phenotypic map by clustering with Phenograph on selected PCs using the Leiden algorithm for community detection. First, we performed a grid search over values of *k* and clustering resolution to ensure robustness to these parameters. Specifically, we computed the pairwise Rand index between each pair of clusterings and found that the clusters are similar within small windows of parameters (i.e. small perturbations in *k* had little impact on the clustering within reason; Rand index > 0.9 for nearly all combinations of *k =* 35 … 50 for fixed resolution = 0.5). We chose one coarse clustering (k = 35, resolution = 0.5), which captures major populations (e.g., grouping all neuroendocrine populations) (**Fig. S1B**) and one more refined clustering (k = 50, resolution = 1.0) which captures highly resolved structure (e.g., separating clusters along resting-to-injured axes) (**Fig. S1C**). We then applied MAST (*105*) for one-versus-all differential expression analysis at both levels of resolution to determine a set of marker genes for each population. We annotated clusters (**Figs. 1B** and **S1B,C**) by inspecting differentially expressed genes (DEGs) and boxplots of log-normalized gene expression across clusters to identify cell types based on the marker lists below. For known intermediate cell states (e.g. Acinar-to-Ductal metaplasia, or ADM), a combination of these markers may be expressed. Other markers are expressed by a more specific subset of the cells assigned to each coarse state (e.g. *Ins* expressed in rare pancreatic cells amongst Neuroendocrine-like cells). Drop-out is observed for lowly expressed genes (such as transcription factors *Ptf1a* and *Bhlha15* in acinar cells).

**Table S10.** scRNA-seq Annotation Strategy.

| Cell Type | Markers |
| --- | --- |
| Acinar (differentiated) | *Zg16, Cpa1* |
| Ductal | *Krt19, Sox9, Clu* |
| Nes+ Progenitor | *Nes, Cd44, Vim, Cdkn2a* |
| Tuft | *Pou2f3, Dclk1, Trpm5* |
| Neuroendocrine (NE) | *Chga, Chgb, Neurod1, Syp, Ppy, Gcg, Ins, Sst* |
| Gastric | *Muc1, Muc6, Gkn3, Muc5ac, Tff2, Tff1, Agr2* |

Finally, we also computed separate embeddings and analyses on subsets of the Progression Cohort, including individual stages or clusters (**Table S11**). In all cases, we performed the same steps above for gene selection, dimensionality reduction and visualization on the log-normalized count matrix of that subset, so as to capture the HVGs and major axes of variation within each group. To maintain consistent cell type annotations, however, each subset analysis retained the original Phenograph clusters computed on the entire Progression Cohort (**Fig. S1B**). In some cases, we chose to visualize a subset with a force-directed layout (FDL) built on a cell-cell affinity graph which accounts for uneven densities along the cellular manifold as in MAGIC (*90*). In brief, we first computed a k-nearest neighbor (kNN) graph on cells based on Euclidean distance between points in PC space using k = 30 by default. This kNN graph was then converted to a cell-cell affinity matrix by applying an adaptive Gaussian kernel (width = 10 for k=30) to edges on the graph and symmetrizing the graph, from which an FDL was computed using Harmony’s plot.force_directed_layout function (*106*). This visualization emphasizes the non-linear continuous structure of the phenotypic manifold measured by a graph, as opposed to tSNE built on PCs, which emphasizes separation of cell states across linear dimensions. The choice between FDL and tSNE for each visualization was based on the expected continuity of discrete cell-states across the progression. For instance, acinar cells of the basal pancreas are expected to be extremely distinct from cell-states of the malignant pancreas. In this case, any spurious edges in the cell-cell affinity graph connecting basal acinar cells to pre-malignant or malignant (K1– K6) cells may inappropriately display a continuity between these populations.

Parameters used for each visualization and analysis metrics for the Progression Cohort subsets are noted in the following table:

**Table S11.** Epithelial scRNA-seq Analysis Groups. *All genes were included for these samples so that heterogeneity driven by lowly expressed receptors or ligands was preserved. **50 PCs (occurring before the knee point) were selected to improve separation of rare cell types in visualization.

| Figure | Analysis | # HVGs | # PCs | Variance Explained | Samples Included |
| --- | --- | --- | --- | --- | --- |
| 2,S3 | Pre-malignant to malignant (K1–K6) | 4000 | 51 | 41.6% | DACD351_Kate_plus;<br>DACD394_Kate_plus;<br>DACD406_Kate_plus;<br>DACD404_Kate_plus;<br>DACD407_Kate_plus;<br>DAC_DI143_Epi;<br>DACD482_Kate_plus;<br>DAC_C301-EPI_1;<br>DAC_C301-EPI_2;<br>Ag-PDAC-PT-Kate;<br>Ag-Lung-Mets-Kate;<br>DAC_D020_p5_Epi;<br>DACC963PT_Kate_plus;<br>DACC963LIVERmet;<br>DACC963_mKate_plus |
| 3, S5 | Pre-malignant scATAC integration (K1–K3, K5) | 4000 | 49 | 41.7% | DACD351_Kate_plus;<br>DACD394_Kate_plus;<br>DACD406_Kate_plus;<br>DACD404_Kate_plus;<br>DACD407_Kate_plus;<br>DAC_DI143_Epi;<br>DACD482_Kate_plus; |
|  |  |  |  |  | DAC_D020_p5_Epi;<br>DACC963PT_Kate_plus;<br>DACC963LIVERmet;<br>DACC963_mKate_plus |
| 4B | Pre-malignant immune integration (K1–K3) | * | 42 | 23.0% | DACD406_Kate_plus;<br>DACD351_Kate_plus;<br>DACD404_Kate_plus;<br>DACD407_Kate_plus;<br>DAC_DI143_Epi;<br>DACD482_Kate_plus |
| 4C,<br>S10E | Regeneration Immune Integration (N1, N2) | * | 63 | 35.9% | DACD511_Kate_plus;<br>DACD550_kate_plus;<br>DAC_B530-Kate+;<br>DACD403_Kate_plus |
| S2C | Neuroendocrine and Tuft (Clusters 9 and 13) | 4000 | 50** | 35.7% | All Progression Cohort |
| S2A,B | Acinar-to-Ductal Metaplasia (Clusters 2, 7, 3 and 12) | 4000 | 88 | 50.4% | All Progression Cohort |
| S1D,<br>4D | Malignant (K5, K6) | 4000 | 57 | 42.8% | Ag-PDAC-PT-Kate;<br>Ag-Lung-Mets-Kate;<br>DAC_D020_p5_Epi;<br>DACC963PT_Kate_plus;<br>DACC963LIVERmet;<br>DACC963_mKate_plus |

##### Perturbation Cohort, shIL33 epithelial cells sorted on mKate^+^GFP^+^CD45^−^

The Perturbation Cohort consists of epithelial *Il33* depleted and control samples to address the effects of cytokine signaling on progression. Individual datasets from this cohort were combined according to stage (K2/K2-sh*Il33* and K3/K3-sh*Il33*) to compare across perturbation and control conditions at each time point, resulting in separate K2 and K3 matrices. We filtered data from each stage for low library size and contaminating mesenchymal cells, as well as for lowly expressed genes (captured in <20 cells) and for ribosomal and MT genes, and log-normalized as described for the Progression Cohort. PCA was applied to 5,000 HVGs in each stage determined using the approach from (*103*) as above. Using automatic knee point detection, we retained 53 PCs describing 28% of the variance for stage K2 and 51 PCs describing 33% of the variance for stage K3.

##### Immune Cohort (K1–K5 & shIL33 immune cells, sorted based on CD45^+^mKate^−^)

The immune data was processed with slightly different steps compared to the epithelial data, due to its unique structural features and the extensive knowledge available on immune subsets. Briefly, we performed the initial preprocessing in two different batches hereafter referred to as Immune Cohorts 1 and 2 (corresponding to Filtering Groups 5 and 6, respectively, **Table S9**) because theses samples were acquired at different timepoints. After removing low quality cells and doublets in each Immune Cohort, we merged Immune Cohorts 1 and 2 to jointly annotate cell types thereby increasing power in detecting rare cell types and ensuring consistent annotations across samples. Cell type annotation was performed in a hierarchical manner first annotating major immune subtypes (T cells, B/plasma cells, myeloid cells) by clustering on the dominant PCs only. After partitioning the data into major immune subtypes, re-normalizing and re-clustering the data on a higher number of PCs, we annotated granular cell types within each major immune subtype. The annotated data was then combined with the epithelial data for the cellular crosstalk analysis. The detailed parameters of this analysis are outlined below.

Immune cohorts 1 and 2 were processed separately as outlined for the Progression Cohort above. After this initial processing, we detected heterotypic doublets (e.g. cells expressing both *Cd3e/Cd3f/Cd3g* and *Cd19/Ms4a1/Ighd*) and therefore introduced a doublet filtering step. Our strategy was based on the idea that doublets will form distinct clusters in phenotypic space, and we chose to cluster cells from all samples within each Group (rather than individually) to ensure that smaller clusters would not be lost.

To prepare for clustering, we filtered genes detected in fewer than 20 cells, as well as MT and ribosomal genes, and cells expressing fewer than 200 genes. We normalized gene expression to median library size and log1p-transformed the data instead of employing scran normalization (used for cell type annotation, below) to avoid nonsensical scran size factors (size factor <0) which may be assigned to a minority of low-quality cells. We then computed HVGs using scanpy’s scanpy.pp.highly_variable_genes function with the seurat_v3 method on raw gene expression counts. However, this method tends to discard genes relevant to cell type annotation, which results in worse cell type separation among the resulting clusters. We therefore added a manually curated list of marker genes to the computed list of HVGs for downstream applications including PC calculation and tSNE embedding (**Table S12**). This list includes 425 key genes for discriminating cell identities or functions of T cells, innate lymphoid cells including NK cells, B cells including plasma cells, and myeloid cells including dendritic cells. After testing a range of 5,000–15,000 HVGs and all genes, we selected 10,000 HVGs and added the 425 marker genes to calculate PCs for both Immune Cohorts, as this led to the best separation of easily confounded cell types in clustering (particularly dendritic cells, mast cells, innate lymphoid cells and plasma cells).

We then clustered the data on PCs using Phenograph, selecting the number of PCs by the knee point of the PC vs. explained variance curve (calculated using the kneed package v0.7.0), or the fewest PCs that explain >20% of total variance, whichever is higher. Based on this procedure, Phenograph was calculated on the first 9 PCs explaining 23.2% or 6 PCs explaining 22.8% of total variance for Immune Cohorts 1 and 2 respectively. We selected *k* = 10 for kNN graph construction because this most clearly delineated clusters containing 20% or more doublets from those with lower doublet content.

To call doublets, we applied DoubletDetection to each immune cohort, and in addition, re-ran DoubletDetection for each sample in Immune Cohort 2 because this detected additional doublets. To filter called doublets, we first eliminated entire clusters comprising >20% doublets, then removed all remaining annotated doublets. We made sure not to remove entire cell subpopulations with biologically sensical phenotypes. Moreover, we removed remaining clusters of obvious heterotypic doublets manually (e.g. cells expressing both *Cd3e/Cd3f/Cd3g* and *Cd19/Ms4a1/Ighd*); these generally overlapped with DoubletDetection annotations but did not reach the 20% threshold. Using this procedure, we removed 3,062 and 349 cells, yielding a final dataset of 22,658 and 12,768 cells for Immune Cohorts 1 and 2, respectively.

To prepare for cell type annotation (**Fig. S7A)**, we combined Immune Cohorts 1 and 2 to increase the power to detect rare immune subsets, and to maintain consistent annotation across analyses. We first partitioned the immune data into major subsets (T/ILC/NK cell, B cell, myeloid cells), focusing on the first few dominant PCs which separated these subsets, and performed finer clustering and annotation on each subset individually as outlined below.

We used scran to normalize the count matrix of the combined Immune Cohort, because median library size normalization can artificially generate differential expression between cells with highly differing library sizes, such as leukocytes (*107, 108*). Scran circumvents this problem by normalizing cells using cluster-specific size factors (*107*). After normalizing raw counts by scran and log1p-transforming the data, we re-calculated HVGs and added the 425 marker genes to the top 7,500 highly-variable genes, which we used to calculate the top 6 PCs explaining 23.6% of total variance. This small number of dominant PCs was used to define the major immune subtypes, whereas normalization and clustering of refined subsets involved many more PCs (see below). We clustered the data in PC space using the immune cell workflow described above and Phenograph k = 50, and ensured that the clustering was stable in a window of adjacent parameters as explained above with a pairwise Rand index of > 0.7.

To assign cell type labels, we manually assessed patterns of mean z-scored marker gene expression in clusters using our custom marker genes (**Table S12**). We also calculated DEGs for each cluster versus all other clusters with scanpy’s scanpy.tl.rank_gene_groups function using the Wilcoxon rank sum test with Benjamini-Hochberg correction. This simple differential expression method was chosen for scalability, given the number of comparisons required for cell type annotation. We then calculated a Szymkiewicz-Simpson overlap coefficient (size of the intersection divided by size of the smaller set) between the top 1,000 DEGs and each marker gene set to assign cell type labels to clusters. This procedure separated leukocytes into three major populations: T/ILC/NK, B/plasma and myeloid cells.

To prepare for more granular cell annotations, we partitioned immune subpopulations and repeated normalization and clustering for each subset separately. We normalized raw counts using scran, log1p-transformed the data, then repeated the calculation of HVGs and added cell-type-specific marker genes (T/ILC/NK, B or myeloid markers) to the top 7,500 HVGs. PCA yielded 48 PCs explaining 20.09% variance for T/ILC/NK cells, 69 PCs explaining 20.05% variance for B cells and 50 PCs explaining 28.9% variance for myeloid cells. We re-clustered the data with Phenograph with k = 40 for T/ILC/NK cells, k = 30 for B cells and k = 50 for myeloid cells, choosing a k value that gives stable clustering in a window of adjacent k parameters as explained above (pairwise Rand indices > 0.7).

We next iterated our annotation procedure using marker genes specific to subtypes of T/ILC/NK, B or myeloid cells (**Table S13**). The rationale for our iterative partitioning approach is based on (1) computational considerations, as it removes major cell-type-specific signals which may confound more granular subtyping, and (2) our biological knowledge of granular cell type markers, which is often limited to comparisons within major subpopulations; e.g. differentially expressed markers between tissue-resident memory (TRM) and naive T cells are well defined, but we know less about how naive T cell markers and TRM markers differ from plasmacytoid dendritic cell markers. A notable drawback of this approach is that during re-normalization with scran, some cells with lower library sizes (e.g. granulocytes and naive T cells) can obtain nonsensical negative size factors, leading to automatic removal. These cells could not be annotated on a more granular level and were removed from further analyses (1.04% of all cells, mostly lower-quality T cells). We also made sure that no known immune cell subpopulations were removed in their entirety by this caveat of scran normalization. Final immune cell annotations are presented in **Fig. S7A**, along with their marker expression patterns in **Fig. S7C**.

Finally, downstream crosstalk analyses were based on these annotated immune cell subsets derived from samples selected for comparison with pre-malignant Progression Cohort (**Table S11** “pre-malignant immune integration” set) with unique processing parameters described below:

**Table S13.**
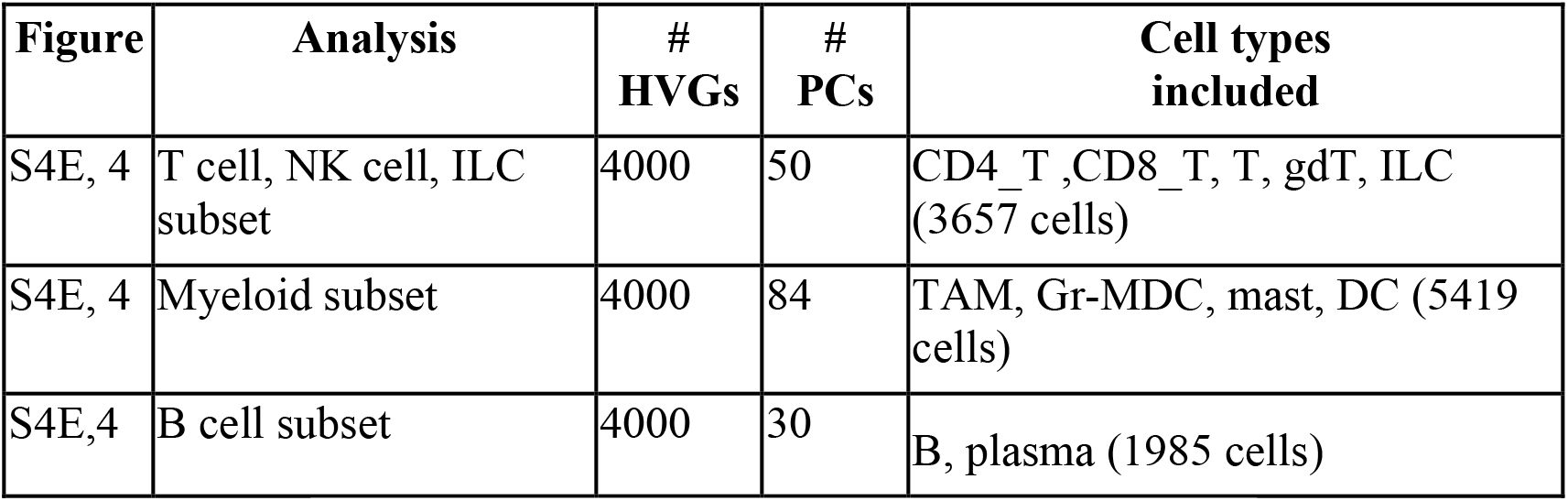
Immune scRNA-seq Analysis Groups.

##### Combined embedding of immune and epithelial data

In addition to the above analyses, we generated one combined embedding including subsets of both Immune and Progression Cohorts to visualize patterns of crosstalk between epithelial cells and their microenvironment (see <u>Heterotypic cell-cell communication analysis</u> below). For this analysis, we combined cells of the Progression Cohort pre-malignant Immune Integration (K1–K3) set (**Table S11**) and the three immune subsets above (**Table S13**). We log-normalized the combined data with standard library size normalization, computed 8,000 HVGs as above, and performed PCA selecting 40 PCs explaining ∼36% of the overall variance. tSNE was applied to these PCs with perplexity = 150.

##### Human scRNA-seq data analysis

In addition to the above GEMM-derived data sets collected in this study, we analyzed previously published data from primary tumors of patients and normal pancreas. We acquired a processed scRNA-seq count matrix combining across samples from (*32*). Median library size normalization was performed. MT and ribosomal genes were removed, as well as genes expressed in fewer than 20 cells and *MALAT1*.

The dataset contained both abundant stromal cells including fibroblasts and endothelial cells as well as immune cells expressing CD45. To exclude these from the analysis for comparison with our epithelial (mKate+) murine datasets, we performed Phenograph clustering with k=100 to capture major subsets and removed stromal clusters (expressing *LUM*, *PLVAP*, and low levels of epithelial markers *CDH1* and *EPCAM*) and immune clusters (expressing CD45 plus subset specific markers such as *CD79A*, *CD3E*, *MS4A1*). The resulting dataset consisted of 20,386 epithelial cells and 20,830 genes for downstream analysis.

#### Prediction of initiating pre-malignant states with scVelo and CellRank

Unbiased analysis of the Progression Cohort with diffusion components reproduced the general chronology from normal (N1) to malignant (K5, K6) states (**Fig. 1C**). However, substantial heterogeneity within each pre-malignant condition, as well as overlapping cell-states across distant time points (e.g. mixing of neuroendocrine-like cells across K1–K3 conditions), suggests that cells are undergoing state transitions that cannot be defined by time series information alone. Most pseudo-time approaches assume cell-states transition from less to more differentiated states from a known origin (*75*). However, regeneration and tumorigenesis involve both differentiation and de-differentiation and origins are not well characterized. Thus we chose to use a RNA velocity based approach to infer directionally in an unbiased manner (i.e. without strong assumptions about origin populations or directionality typically required for trajectory inference (*75*)).

RNA velocity (*49*) has been shown to successfully infer directionality in systems of differentiating cells using ratios of spliced and unspliced transcripts. The RNA velocity vector of a given cell predicts which genes are currently being up- or downregulated and points towards the likely future state of that cell (visualized as arrows projected on FDL coordinates, e.g. **Fig. S3A**). However, RNA velocity is only able to capture short term transcriptional dynamics (∼1 day) and we therefore limited our analysis to the earliest time points (K1 and K2). We used the pre-processing functions of Velocyto (*49*) to efficiently separate reads from K1 and K2 BAM files into spliced and unspliced count matrices, and to generate loom files for downstream velocity algorithms. However, Velocyto assumes that a subset of cells are in steady state, which is unlikely to be true of cells responding to oncogene activation and/or injury as in our case; we thus opted to use the scVelo algorithm (*48*), which can model expression dynamics which are not in steady state. We first applied log-normalization and filtering of lowly expressed genes (<20 spliced and/or <20 unspliced transcripts) in scVelo to retain 9,294 genes. We then applied the scVelo velocity estimation in “dynamic” mode with default parameters on this subset of genes (**Fig. S3A)**.

However, individual velocity estimates are known to be very noisy and these individual vectors do not reveal origin states. Moreover, projecting the high-dimensional velocity vectors to 2D often fails to faithfully capture the global transcriptional dynamics. To infer global dynamics of the system, CellRank (*47*) combines gene expression similarity with RNA velocity to robustly estimate directed trajectories of cells. The robustness is gained through the use of the similarity-based neighbor graph and cell-state transitions are modeled as directed random walks along this graph. The more a neighboring cell lies in the direction of the velocity vector, the higher its transition probability. However, unlike RNA velocity, all transitions are enforced to remain within the phenotypic manifold. By globally modeling the cell-state transitions as a Markov chain, CellRank is able to successfully identify initial (‘apex’) states, inferred to be the sources of the cell-states observed in the data.

We ran CellRank with default parameters, applying the initial_states function to identify 4 high-confidence originating apex populations with an initial state probability cut-off automatically determined by the algorithm’s eigengap-based threshold (**Figs. 2A** and **S3A**). A set of marker genes, derived from populations whose initiating potential was discerned through Cre-mediated lineage tracing of neoplastic pancreas in previous works (*27, 50–53*), exhibited strong concordance with CellRank-predicted initial states (**Fig. S3B**).

#### Bulk ATAC-seq data generation, processing and integration

The substantial heterogeneity generated from high-potential oncogenic states identified by CellRank occurs rapidly (24–48 hpi), a time scale that is more consistent with changes in chromatin accessibility than genetic mutation. To test whether epigenetic features can explain observed scRNA-seq heterogeneity, we integrated bulk ATAC-seq data from published studies for stages N1-N2, K1-K2, and K5 (*25*). We also collected additional samples from intermediate pre-malignant conditions K3-K4 to complete the Progression series. To do so, we generated new epigenomic data from mKate2+ cells FACS-isolated from benign neoplasia (PanIN) tissue states (K3, K4), as previously described (*25*), and analyzed it with previously generated bulk ATAC-seq profiles from mKate2+ cells isolated from pancreata representing normal (N1), regenerating (N2), pre-neoplastic (K1, K2) and malignant (K5) states (extracted from GSE132440).

These data sets were processed as described in (*25*). Briefly, trimmed paired-end reads were aligned to mm9 using Bowtie2 (*109*), peaks were called for each sample individually with MACS2 (*110*), and a global atlas was derived by merging peaks from each sample within 500 bp of one another. Read counts within each peak were normalized with DEseq2 (*111*) to account for differences in per-sample sequencing depth.

##### Identification of chromatin modules describing progression

We sought to learn broad accessibility patterns over stages, and to map these to single cells. We first called differentially accessible peaks across early stages (N1-N2, K1-K4) in our bulk ATAC-seq data using DEseq2 (log_2_-transformed fold change ≥ 0.58 and FDR ≤ 0.1). To identify global progression trends relevant to early progression, we performed PCA on normalized differential peaks contrasting all pre-malignant conditions pairwise, then visualized the first two PCs explaining ∼54% of total variance (**Fig. 2B**). The distribution of samples in this embedding suggests consistent, distinct genome-wide accessibility patterns defining replicates (individual mice) of the same stage.

Next, to study the specific accessibility patterns underlying these trends, we applied kmeans with n=15 clusters to peaks based on the z-scored accessibility across samples. This clustering organized peaks into several dominant patterns of regulatory dynamics (henceforth “chromatin modules”), comprising peaks only accessible in normal pancreas (Normal chromatin module), peaks which become most accessible in early PanINs (Benign Neoplasia chromatin module), and peaks which become most accessible in PDAC (Malignant chromatin module) (**Fig. 2C**). Notably, we observed that peaks of the Benign Neoplasia module are closed in malignancy, despite the chronological relationship between pre-malignant PanIN lesions (K3, K4) and their eventual malignant PDAC states (K5, K6). However, some trends exist across modules: both Benign Neoplasia and Malignancy-associated peaks are accessible to some extent in K1 and/or K2 before becoming predominantly accessible in K3-K4 and K5-K6, respectively.

To group peak clusters according to these patterns, we removed clusters from downstream analysis if they did not clearly fit in one of the 3 modules of interest, and visualized the resulting peak groups by heatmap (**Fig. 2C**). Manual curation was preferred over automated cluster assignment to ensure that clusters lacking coherent patterns over progression were not included in each module.

##### Integration of bulk chromatin with single-cell RNA-seq

We investigated chromatin-directed gene expression at the single-cell level by integrating bulk-derived chromatin modules with K1– K6 scRNA-seq data from the Progression Cohort (**Table S11**). First, we mapped peaks to associated genes using ChIPseeker (*112*) as in (*25*). We then aggregated all genes mapping to peaks associated with each module, yielding three module-associated gene signatures. We identified unique genes in each signature to define mutually exclusive patterns associated with each chromatin module (**Table S2**). Finally, we z-scored the expression of each individual gene across each cell to emphasize those that are relatively overexpressed in each cell, and visualized each module’s relative activity as the average z-score of the genes in that module (**Fig. 2D**).

#### Predictive model of late-stage fates

Our analyses of chromatin dynamics generated three complementary observations: (1) a divergence in PC space between Benign Neoplasia (K3, K4) and Malignant (K5, K6) stages (**Fig. 2B**), (2) an early increase in accessibility (K1, K2) of regulatory elements that become highly accessible in Benign Neoplasia and Malignant modules (**Fig. 2C**), and (3) an association of early (K1, K2) cell state expression with one or more late-stage modules (**Fig. 2D**). Based on this, we hypothesized that *Kras* mutant cells are epigenetically primed, such that their early chromatin accessibility patterns establish propensities for Benign and/or Malignant fates. To define epigenetic priming in concrete terms, we assert that cells primed for different fates can be discerned by the presence of open chromatin proximal to fate-specific genes, prior to the establishment of fate-associated states.

To quantitatively evaluate this hypothesis, we first assumed that gene expression is a reasonable surrogate for accessibility of that gene’s regulatory elements for at least a subset of genes, an assumption which derives from the observed correspondence between expression state and chromatin modules derived from bulk ATAC-seq (**Fig. 2D**). We thus aimed to devise a computational framework which can identify transcriptional signatures that discriminate between Benign and Malignant phenotypes, then use it to probe for these programs in early tumorigenesis. To this end, we trained a classifier to distinguish Benign and Malignant fate labels using single-cell samples from late timepoints (PanIN K3, K4 and PDAC K5, K6). The resulting expression-based predictive model of cellular fate was used to interrogate cells from early pre-neoplastic time points (K1, K2) for these fate-associated programs. We assume the inferred probability distribution of distinct fates for each early cell reveals its tendency to skew toward particular expression endpoints.

Specifically, we used a logistic regression multi-class classifier on log-normalized expression features to predict stage labels. We randomly split late-stage cells (K3–K6) into a training set (10,758 cells, or 80% of late-stage data) and a held-out validation set (2,689 cells/20%). We trained the model with sklearn (*113*), which provides a maximum likelihood estimate of logistic regression parameters to predict stage labels (K3, K4, K5, or K6) in the training data. Our model performed well on held-out data (accuracy = 0.9966, precision = 0.9973), suggesting that gene expression features are informative of late-stage cell fates and that our classification strategy captures fate-relevant patterns. We further inspected coefficients of our fitted models to identify strongly predictive genes for each late-stage phenotype, finding expected programs such as EMT (*Epcam and Vim*), MYC activity, and tumor suppressor activity distinguishing Benign from Malignant phenotypes (**Fig. S3C**).

Next, we applied our trained classifier to pre-neoplastic (K1, K2) cells, and visualized the per-cell probabilities of each fate category. We collapsed the classifier into Benign and Malignant probabilities by summing probabilities for K3-K4 and K5-K6 cells, respectively (**Figs. 2E and S3D,F**). We then binned early cells into categories: highly associated with a Benign state (Malignant probability <0.4); highly associated with a Malignant state (Malignant probability >0.6); and mixed state (0.4 <= Malignant probability <= 0.6), representing rare cells (<5% of K1-K2 cells) that express a composite Benign-Malignant program (**Fig 2E**). We find uninjured (K1) cells are enriched in the Benign class, while injured cells are enriched in the Malignant class (Fisher’s exact test, odds ratio = 17.3671, p-value = ∼0.0), recapitulating previous reports that injury shifts the oncogenic pancreas epithelia toward a cancer-like state (*25*). To visualize intermediate states, we computed the density of these cells in PC space (using significant PCs, described above in <u>Single-cell RNA-seq data processing</u>) with a Gaussian kernel density estimate via scipy.stats.gaussian_kde with default bandwidth parameters, which we visualized in two dimensions on a FDL (**Fig. S3E**).

Intermediate or fuzzy classification to one or more states can occur for one of two reasons: cells either simultaneously express programs of more than one class, or cells can express neither class-associated program. As such, it was important to ensure that both Benign and Malignant programs were detectable in a majority of our mixed probability cells (0.4 <= Malignant probability <= 0.6) to designate them as composite states. We developed a gene signature for each program by extracting the genes with the top 200 coefficients (by coefficient value) amongst Benign (K3) and Malignant (K5) logistic regression models. We then intersected these gene lists with Benign chromatin module and Malignant chromatin module genes respectively to ensure that these transcriptional programs were primarily associated with underlying epigenetic differences. We then computed a separate Benign and Malignant signature score per cell by taking the average z-scored expression of each gene in the signature. **Fig. S3G** confirms that true Benign (K3, K4; pink contour) and Malignant (K5, K6; purple contour) cells are clearly distinguishable based on these values, and further shows that the majority of mixed probability cells (orange) assume an intermediate value between these modes. Importantly, many of these mixed cells highly express both programs, suggesting that many assume a truly composite phenotype with respect to late-stage fates.

#### Single-cell ATAC-seq processing and analysis

*Initial processing*: To extend and support our analysis of chromatin dynamics, we analyzed scATAC-seq data from comparable or paired (i.e. from the same mouse, **Table S14**) pre-malignant and malignant conditions (K1–K3, K5). Two samples were derived from a previously published study (*25*) and the remaining were collected for this work as described above.

**Table S14.** scATAC-seq Sample Pairings.

| Sample ID | Stage | Genotype | Paired scRNA-seq Sample ID |
| --- | --- | --- | --- |
| DACD408_b_mKATE2_ATAC | K1 | KC;RIK | N/A |
| DACD404_b_mKATE2_ATAC | K2 | KC;RIK | DACD404_Kate_plus |
| DACE270_Epi | K3 | KC;RIK | N/A |
| DACE271_Epi | K3 | KC;RIK | N/A |
| DACC963_PT_B_mKate | K5 | KPflC;RIK | DACC963PT_Kate_plus |
| DAC_D020_p5_Epi_ATAC | K5 | KPflC;RIK | DAC_D020_p5_Epi |

Samples were processed individually using a modified CellRanger ATAC pipeline as described in (*25*) to derive alignments to mm10, then analyzed with ArchR (*114*), a tunable pipeline for filtering, normalization, dimensionality reduction and visualization of aligned scATAC-seq fragments. We chose ArchR for its flexibility in representing accessibility features in defined regulatory elements, proximal to genes of interest, or globally in bins along the genome. This presents an advantage over approaches focused on accessibility solely in inferred regulatory elements (i.e. called peaks), which may disregard dynamics driving rare cell populations (from false negative peak calls) or erratic dysregulation that may occur in cancer (resulting in many noisy peak calls). We applied a custom modified version of ArchR (*55*) which is better suited to capture chromatin features of rare populations, as described below.

To begin, individual CellRanger ATAC fragment files from all stages (K1–K3 and K5) were combined into a single dataset in ArchR, and basic filtering was performed with parameters filterFrags = 8000 (determined by manual inspection of the fragment count distribution) and maxFrags = 1e+7. Doublet score computation and doublet filtering were performed with default parameters, resulting in 19,666 cells. For consistency with scRNA-seq analysis, we removed mesenchymal contaminants, yielding a final count of 18,211 cells. We then derived the two standard ArchR feature representations for our scATAC-seq data:

*Tile Features* bin counts in 500-bp windows across the accessible genome, which allows flexible representation of the data without peak calling error or bias toward coding regions. Tile features tend to produce embeddings with the most faithful phenotypic structure, hence we used them for major downstream analyses such as dimensionality reduction, visualization and clustering.

*Gene Score Features* are distance-weighted aggregations of accessibility proximal to each gene that allow functional interpretation of the data. Using our modified pipeline, we only computed gene scores on regions identified as highly variable across clusters to improve the extent of heterogeneity captured by these values. We used the resulting scores for cell type annotation and integration with scRNA-seq features (See <u>Identifying and integrating pancreas metacells</u> below).

##### Dimensionality reduction and visualization

To compute a low-dimensional embedding of single-cell epigenomes, we applied ArchR’s iterative Latent Semantic Indexing (LSI) function to the tile matrix on 150,000 highly variable features. Each round of LSI generates a preliminary low-dimensional embedding from which a clustering is estimated. Variable tile features are then computed across clusters to select the most informative tiles with respect to the major phenotypes. In our case, we modified parameters in each LSI iteration to capture more refined cell types; a clustering resolution parameter of 0.1 distinguished small cell populations (e.g. tuft cells) better than default parameters. LSI components from ArchR were then used as input to compute visualizations, and for downstream analysis in python. As described above (see <u>Single cell RNA-seq data processing</u>), we computed an FDL on the cell-cell affinity matrix constructed from Euclidean distances in LSI embedding coordinates (as opposed to PCs used for scRNA-seq) (**Figs. 3A** and **S4A,B,D**). LSI components are directly visualized in **Figs. 3B** and **S4B**.

##### Clustering and annotation

To then identify discrete cell types, we clustered scATAC-seq data with Phenograph (k=30) using the Leiden algorithm on LSI component features. At this level of resolution, we achieved desirable granularity with respect to rare cell types (i.e. tuft cells), but also observed several groups of highly-related clusters with shared accessibility at key markers (**Table S15**) which may potentially originate from a single cell type. To quantitatively identify these phenotypically-similar clusters, we examined each initial Phenograph cluster for overlap in accessibility patterns proximal to genes. We first performed a Wilcoxon rank test (the default approach in ArchR for differential analysis) for each gene’s accessibility in one cluster versus all others based on values in the gene score matrix (described above) using the getMarkerFeatures function. For each cluster, we filtered the resulting significant genes (FDR<=0.01) to retain those with a log2 fold change of at least 1.25 as marker genes. We then computed the percentage of shared marker genes between each pair of clusters, indicating extent of phenotypic similarity between clusters at the chromatin level. We observed substantial sharing of marker genes between several pairs of clusters (minimum 15% shared genes), which we reasoned could be merged for annotation and downstream analysis. Resulting merged clusters are displayed in **Fig. S4A**.

We then identified marker genes for the merged clusters once again following the Wilcoxon rank method on gene scores in one merged cluster versus all others. We visualized shared marker genes between pairs of merged clusters as before and observed elimination of high inter-cluster sharing (<15% shared), thus confirming the merged clusters as highly phenotypically distinct. Lastly, to identify the cell type of each cluster and further validate the merging, we computed the average gene accessibility scores of every marker gene and compared the results against known biological signatures for various pancreas populations. We identified individual merged clusters associating with gastric-like, *Nes*^+^ progenitor-like, PDAC, neuroendocrine-like, ADM, and tuft cell populations, in concordance with our scRNA-seq analysis, according to the following annotation criteria:

**Table S15.** scATAC-seq Annotation Strategy. *Due to low accessibility captured near cell-state markers (e.g. *Dclk1*), tuft cells in scATAC-seq data are mainly identified by genome-wide similarity to tuft cells identified in scRNA-seq (see <u>Identifying and integrating pancreas metacells</u>). **Broadly accessible in metaplastic cells.

| Cell Type | Markers |
| --- | --- |
| Acinar (differentiated) | *Ptf1a, Bhlha15, Cpa1, Nr5a2* |
| Ductal | *Krt19\*\**, *Sox9\*\** |
| *Nes*<sup>+</sup> Progenitor | *Nes, Cd44, Vim* |
| Tuft* | *Vill* |
| Neuroendocrine (NE) | *Chga, Chgb, Neurod1, Ppy, Sst* |
| Gastric | *Gkn3, Tff2, Tff1, Agr2* |

##### ATAC-seq signal tracks

We visualized signal tracks of scATAC-seq profiles aggregated over clusters or samples using custom python code utilizing the tabix package (*115*) (**Figs. S4C** and **S6A**). For each track, we aggregated reads from fragment files in a 5-kb to 50-kb window around a gene, and normalized coverage to the total depth of cells included in the cluster or sample set of interest, to provide comparable scale across tracks. Coverage was smoothed using a window size of 100 bp to aid visualization.

#### Metacells algorithm

Clustering and cell-type annotation suggest that cell-states derived from scRNA-seq and scATAC-seq data match at the broad cell-type level, but we also find substantial heterogeneity within each cluster. For example, the *Nes*^+^ Progenitor population (coarse Phenograph Cluster 1) includes a spectrum of phenotypes primed for distinct fates (PDAC probabilities spanning from nearly 0 to 0.9999). We therefore needed a higher-resolution mapping between epigenetic and transcriptional states, yet the extreme sparsity of scATAC-seq hinders our ability to characterize chromatin states of individual cells. Our first goal for data integration is thus to overcome the sparsity of single-cell profiles while maintaining a high-resolution of diversity in cell-states.

We addressed this challenge by developing an algorithm that aggregates cells which share the same state and an intermediate resolution between single cells and typical cell-state clusters, inspired by the metacell approach (*54, 55*). Each metacell represents a distinct, highly granular and homogenous cell state, such that differences between cells within a given metacell are due to technical noise rather than biological disparities. As such, aggregating counts across cells in each metacell overcomes noise and dropout, providing a robust characterization of that distinct cell state. The concept of metacells was introduced by Baran and colleagues (*54*), but the accompanying method is unsuitable for scATAC-seq, and it removes outliers aggressively. Our algorithm is non-parametric (requiring little parameter tuning by the user) and flexible to diverse modalities, including both scRNAseq and scATAC-seq. In this work, we aimed to derive metacells from epigenomic and transcriptomic data separately, so that they could then be reliably integrated across the span of progression.

Our algorithm first constructs a kNN graph over cells, determining neighbors based on Euclidean distances in PC space for scRNA-seq or iterative LSI space for scATAC-seq (computed as in <u>Single-cell RNA-seq data processing</u> and <u>Single-cell ATAC-seq processing</u> above). The algorithm then constructs a shared nearest neighbor (SNN) graph by pruning edges in the kNN graph that are not bidirectional. The remaining edges are mutually highly similar to one another and better reflect the phenotypic manifold; they are weighted using an adaptive bandwidth Gaussian kernel as previously described (*90*). Grouping highly similar cells within small neighborhoods on the SNN graph should then produce a representative sampling of cell states, provided that the resulting metacells (1) are reasonably distinct from one another and (2) sample all regions of the graph to capture the full extent of heterogeneity. To encode this intuition, metacells are initialized with cells that occupy an “extreme” position on the manifold according to their *leverage*, representing geodesic distance to all other cells in the graph (*116*).

We first compute leverage for each cell *i*:

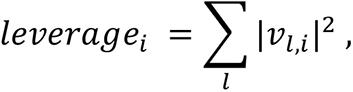

where *v_l,i_* is the i^th^ element of the l^th^ eigenvector of the normalized SNN graph. Representative cells (metacell “centroids”) are then initialized by randomly sampling cells in SNN graph space in proportion to their leverage using the following sampling procedure for each cell *i*:

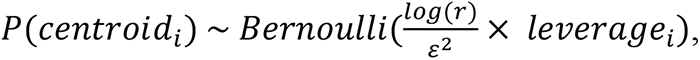

where *r* is a user-defined rank parameter scaling the Bernoulli probability parameter (and therefore controlling the number of centroids selected) and *ε* is an error tolerance parameter. Unselected (non-centroid) cells are then assigned to the nearest centroid to form groupings of highly similar cells, after which the centroid is updated iteratively by picking a new cell that minimizes the distance to cells of that group. On each iteration, metacells consisting of fewer cells than a user-defined threshold are pruned to limit sparsity of the resulting profiles. This procedure is continued until no centroids are updated between iterations or a maximum number of iterations is reached; the result is that every cell is assigned to a distinct metacell. Expression or accessibility features are then computed for each metacell by averaging across constituent cells.

#### Identifying and integrating pancreas metacells

To identify metacells in scATAC-seq data, we applied the above algorithm using the 30 LSI components from ArchR as features for SNN graph construction. For metacell computation, we used k = 30 nearest neighbors, a rank of 200, and minimum metacell size of 10 cells to capture heterogeneity within rare cell populations. This resulted in 264 epigenomic metacells spanning conditions and clusters. For scRNA-seq metacells, we pooled a subset of data from comparable conditions (K1–K3, K5, K6; **Table S11**), excluding one tumor for which we did not collect scATAC-seq to avoid confounding intra- and inter-tumor heterogeneity. For one case in which the primary tumor sampled few transcriptomes (n = 261), we included both the primary tumor and metastases—despite lack of scATAC-seq data for metastases—in order to capture enough malignant cells for robust data integration. We applied our metacell algorithm on 49 selected PCs (chosen using the knee point method; **Table S11**) from the log-normalized scRNA-seq data subset (processed as in <u>Single-cell RNA-seq data processing</u> for Progression Cohort). Parameters consistent with epigenomic metacell computation (k = 30, rank = 200, 10 minimum cells per metacell) were used to derive a comparable set of 230 transcriptomic metacells.

Next, we constructed a common feature space to match epigenomic to transcriptomic metacells. We treated ArchR gene scores as a proxy for expression, assuming that, on average, cells with high expression for a given gene will also have relatively high accessibility in the vicinity of that gene (and vice versa for low expression). We first averaged gene expression over cells in each transcriptomic metacell, and averaged ArchR gene scores over cells in each epigenomic metacell, to obtain complementary matrices. Next, we z-scored every gene separately within each data modality to derive a standardized metacell matrix for each that was of comparable scale. Finally, we computed a PCA embedding on the standardized, concatenated matrix containing both epigenomic and transcriptomic metacells, to produce a common reduced-dimension space for downstream computation.

To pair RNA and ATAC metacells, we computed a Mutually Nearest Neighbor (MNN) graph across data modalities with k = 30. Magnitude differences between modalities may impact comparison, even with z-scored data; we thus used cosine similarity as a distance metric for nearest neighbor computation to de-emphasize such differences. Moreover, cosine similarity captures an element of gene-gene relationships which is better conserved across data modalities (*100*). Given these mutually nearest pairings, we sought a common visualization across modalities that preserves the general structure of the phenotypic space and reflects both cross-modality and intra-modality cell-state similarities. To achieve this, we first created a metacell-by-metacell combined graph with dimension *N* = *N_t_* + *N_e_* where *N_t_* and *N_e_* represent the number of transcriptomic and epigenomic metacells, respectively. We defined edges between metacell nodes as a weighted sum of cross-modality MNN edge weights and intra-modality nearest neighbor (NN) graph (also computed on cosine distance) weights:

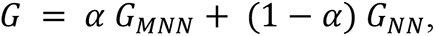

where *α* is a parameter that defines the trade-off between cross-modality and intra-modality cell-state similarity. We set *α* = 0.4 to slightly favor the phenotypic landscape and projected the data as an FDL on *G* (**Fig. S5A**). In this layout, we found that metacells group by their predefined clusters, and transcriptomic and epigenomic metacells belonging to the same cell state tend to co-cluster. Indeed, standardized accessibility and expression show strong concordance for key marker genes (*Ptf1a, Tff2, Nes*, and *Neurod1*) in this visualization (**Fig. S5B**).

We proceeded to further validate the agreement between accessibility and expression for our MNN pairs. For each transcriptomic metacell, we computed a corresponding average accessibility profile across its epigenomic MNN pairs by first averaging the non-standardized ATAC signal across neighbors of each RNA metacell, then z-scoring the averaged value. We then inspected the per-gene Pearson correlation of expression and aggregated accessibility, finding that the majority of genes have strong positive correlations, particularly cell-type markers such as *Nr5a2, Krt19, Tff2,* and *Nes* (**Fig. S5C**). This demonstrates the consistency in our computationally derived pairings between transcriptomic and epigenomic datasets.

#### Measuring epigenetic plasticity

With MNN analysis, we matched transcriptomic and epigenomic metacells based on the *most similar* states. However, various degrees of gene priming—open chromatin in unexpressed regions—should lead to mismatched chromatin and expression features for a subset of genes. We therefore asked whether epigenomic metacells might be similar to additional transcriptional states beyond their MNN pairs. We computed pairwise correlations on expression HVGs between each transcriptomic and epigenomic metacell, and observed many cases in which metacells from the two modalities were highly correlated despite representing disparate cell-states (**Fig. 3C**). Epigenomic metacells with similarity to multiple transcriptional states may encode multipotential states, which we define to be a cell’s epigenetic plasticity.

##### Quantifying epigenetic plasticity

We assume that (1) plastic cell states have *access to* diverse transcriptional programs that drive distinct cell phenotypes, and that (2) a common mechanism for providing access is epigenetic priming, i.e., opening the proximal chromatin of fate-associated genes prior to receiving fate-specifying signals. To quantify this plasticity, we use a classification-based approach to detect cell-type-specific gene expression encoded at the chromatin level (see similar approach in <u>Predictive model of late-stage fates</u>). Classification methods are adept at learning relevant features (e.g. gene expression programs) to discriminate classes (e.g. cell-states). Such approaches can extrapolate knowledge learned from training data to held-out datasets, or in our case, from one data modality (gene expression) to a second (chromatin accessibility). We reasoned that a classifier trained to learn cell-state-specific features in transcriptomic data could be used to predict expression phenotypes from chromatin-derived features. Furthermore, we posit that uncertainty in such predictions is a measure of plasticity; epigenomic states that map to a wide range of transcriptional cell types have the greatest potential to express diverse phenotypic programs. This concept is summarized in **Fig. 3D**.

We first defined discrete expression phenotypes using our refined Phenograph scRNA-seq clusters to capture a finer granularity of transcriptional states, particularly those separating injured and uninjured states (**Fig. S1B**, see <u>Single-cell RNA-seq data processing</u> for details). To ensure robust classification, we discarded rare clusters (<200 cells) corresponding to 6 metacells; all remaining clusters had reasonable amounts of data for training. Using this filtered dataset, we trained a multiclass logistic regression classifier with sklearn on transcriptomic metacells to predict cell-state label (Phenograph cluster of the metacell center) from aggregated, standardized gene expression features for that metacell. We trained the model on a subset of 60% of transcriptomic metacells (N = 138) and achieved strong predictive performance on a held-out validation set (accuracy = 0.9457, averaged precision = 0.9489, averaged recall = 0.9444), indicating that the model can successfully identify phenotype-specific features in gene expression.

Our classifier provides a function *f* which takes as input an expression program of metacell *i* (with dimension *D* = number of genes) and outputs a probability distribution over cellular state (with dimension = number of transcriptomic states), where a class label is given as:

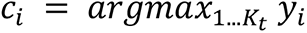

We next asked, for each cell-state, whether using accessibility features as a proxy for expression results in clear mapping to the expected cell-state, or an uncertain mapping to more than one cell-state, suggesting chromatin primed for diverse gene programs. Specifically, we applied our logistic regression model (trained on transcriptomic metacells) to classify epigenomic metacells by using accessibility features for *x*, thereby assigning a most-probable Phenograph cluster class (initially identified from expression data) to each epigenomic metacell.

To summarize the predictions, we counted the number of epigenomic metacells from each epigenomic cluster that classified to each transcriptomic cluster. For the set of epigenomic metacells belonging to epigenomic cluster *k_e_*, we computed as the number of metacells in which classify to transcriptomic cluster *k_t_* as follows:

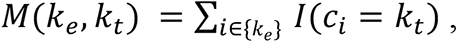

where *I* is the indicator function, and *c_i_* is the class label, as described above. These values define a “confusion matrix” with dimension *K_e_* (number of epigenomic cell states) by *K_t_* (number of transcriptomic cell states), where the element *M*(*k_e_*, *k_t_*) quantifies the extent to which each epigenomic state (rows) classify to each transcriptome state (columns). We then examined this confusion matrix, standardizing across rows to emphasize the tendency of each distinct chromatin profile toward various expression states (**Fig. 3E**). Highly plastic states can thus be identified as epigenomic clusters (rows) containing metacells that map to multiple transcriptomic cell states (columns), particularly from disparate cell types. Besides this approach, which captures only the most likely transcriptomic class per epigenomic metacell, we further summarize these cross-modality mappings as the average log probability per epigenomic state (row) toward each transcriptomic state (column) in **Fig. S5D**. This emphasizes plasticity in certain states whose probability distribution is peaked near their correct class (e.g. tuft cells), but have slight probabilities toward other classes. This is in contrast to acinar-like ADM cells, which consistently classify to ADM states and rarely have any substantial probability of classifying to another state.

##### Calculation of plasticity score

To enable analysis beyond coarse cluster-level metrics, we quantified this observed plasticity on a *per-cell basis* by computing Shannon entropy of the probability distribution *y* across classes from the classifier predictions:

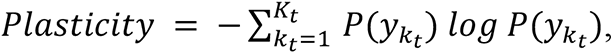

for each transcriptomic state *k* = 1…K. This score will be high when the probability is widely distributed across many transcriptomic classes, hence a metacell’s epigenome displays accessibility for diverse gene expression programs. Conversely, it will be low when the distribution peaks at a single cell-state, indicating that a metacell’s epigenome is strongly linked with its own expression readout.

Measurement of plasticity on a numerical scale enables us to address questions about which factors (e.g. gene programs, stage in progression) influence plasticity. In **Fig. 3F**, we visualize its distribution across cell-states to identify the states which display high epigenetic plasticity.

Furthermore, we find that plasticity is enhanced upon inflammation by comparing distributions of plasticity scores per metacell in *Kras* mutant cells (K1) versus those responding to inflammation (K2) (**Fig. 3H**).

##### Robustness of plasticity approach

To test whether our plasticity metric is robust to cell-states captured (i.e. rare populations), we randomly subsampled transcriptomic metacells 100 times, selecting 75% of profiles with replacement for individual trials. Each trial measures whether our discriminative model of cell-state is consistent with respect to observed transcriptomic profiles. Indeed, we found high concordance between plasticity scores derived from each trial and the true scores, with average Pearson correlation coefficient of r = 0.895 (minimum r = 0.703) between subsampled and true scores for each epigenomic metacell. Subsampling transcriptomic metacells to 50% in this procedure maintained strong concordance (average r = 0.8565, minimum r = 0.6177).

##### Annotation of plasticity score with GSEA

Beyond identifying high-plasticity populations, our metric serves as a powerful quantitative tool to explore features of plastic cells. In particular, by ranking metacell profiles by plasticity score, we can correlate gene programs with epigenetic plasticity. To this end, we computed Spearman correlations between each gene’s accessibility score and the corresponding plasticity score per cell, and used these values to rank genes for Gene Set Enrichment Analysis (GSEA) (*57*). GSEA was performed with the gseapy python package using all genesets included in the “KEGG_2019_Mouse” library (*88, 89*), with an FDR of 0.1 (**Fig. 3G, S5E**).

#### Heterotypic cell-cell communication analysis with Calligraphy

Our results collectively suggest that inflammation is associated with plasticity in pre-neoplasia, prompting us to further explore potential interactions with the immune microenvironment in pre-malignancy. Many approaches exist to infer cell-cell interactions via expression of cognate receptor-ligand (R-L) pairs across cell types, defining significant interactions with respect to single pairs of R-L genes against a null model of R-L expression (*61, 62*). However, their sensitivity and specificity is impacted by the limited capture rate for individual genes (particularly stable cell-surface proteins), as well as weak support for each of many potential interaction pairs. Some methods address weak or noisy signal by incorporating gene expression downstream of signaling, but these approaches derive gene pathways from general databases without regard to cell-type-specific mechanisms (*62*). This has the disadvantage of ignoring context-specific, cell-intrinsic signaling, including the many pathways impacted by oncogene activation that are relevant to our setting.

In this study, both bulk and single-cell epigenetic data revealed considerable remodeling of chromatin structure proximal to communication genes (e.g. those encoding for inflammatory receptor and ligand proteins). Given this pervasive remodeling of a large number of communication genes and the modular nature of gene regulation, we hypothesize that the remodeling of communication genes is also organized into coregulated modules, which can be exploited to strengthen the detection of true signal in the data. Indeed, expression-based clustering of communication genes exhibits striking modularity (**Fig. 4A**). We therefore designed a new approach to crosstalk inference, Calligraphy, which is context-specific and uses gene coregulation to improve the sensitivity and robustness of inferred interactions.

Calligraphy is rooted in *modules* of inflammation-associated genes: each cell can receive signals based on its expressed receptors and send signals based on its expressed ligands. Specifically, modules are sets of communication genes that tend to be mutually expressed in the same populations, and hence summarize the possible incoming and outgoing communication for a particular state. To identify these patterns, Calligraphy builds a co-expression network of communication genes, from which robust inferences can be made across sub-populations representing coherent inflammatory programs. Prior knowledge of potential R-L relationships informs the communication potential across cell states based on their relative module usage, drawing from data on the numerous communication genes in each module to predict possible interactions. Below, we describe how we infer communication modules *de novo* from scRNA-seq data, then apply a module-based approach to map potential crosstalk in the pre-malignant pancreas.

##### Co-expressed communication module detection

We first mapped communication gene co-expression patterns across the pre-malignant landscape. To account for gene drop-out, we imputed gene expression using MAGIC (*90*), which has been shown to increase power to detect co-expression trends in single-cell data. Specifically, we applied MAGIC to the log-normalized count matrix of pre-malignant K1–K3 stages, which generates a cell-cell affinity graph using defaults k = 5 and t = 3. The t parameter was chosen to only produce modest smoothing along the manifold and avoid over-inflating the expression of true negative genes.

We visualized the MAGIC-imputed gene-gene correlation matrix across all annotated communication genes to expose potential modules (**Fig. 4A**). The matrix exhibits a striking degree of structure; blocks of mutually correlated communication genes representing inflammatory modules within the oncogenic epithelia are readily apparent. We also observed substantial variability in correlation magnitudes outside of coherent blocks, suggesting the need for a module detection approach that is robust to spurious correlations induced by imputation and noisy data. Furthermore, large blocks of off-diagonal correlation implied substantial sharing of specific communication genes across programs and/or cell-states. This finding highlights an important biological feature of signaling: many individual communication genes are expected to participate in the communication of multiple cell-states, including immune subsets and epithelial states. This motivated a second criterion for our module-based crosstalk approach: flexibility with respect to sharing of receptors or ligands across distinct communication modules.

Our module detection approach with Calligraphy thus begins with thresholding the gene-gene correlation matrix (Pearson’s correlation coefficient >= 0.4) to derive a graph, in which genes are nodes and correlation-weighted edges connect highly correlated genes. We stored and computed all graph information using the networkx python package for efficient network handling (*117*). To improve module coherence and remove spurious associations, we compute a Jaccard similarity metric accounting for the degree of neighbor-sharing between all pairs of genes, similar to (*101*), as follows:

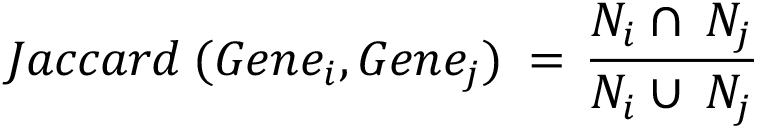

where *N_i_* is the set of neighboring genes for gene *i*. We define an upper and lower threshold for this Jaccard metric, retaining existing edges which meet the lower threshold and appending new edges which meet the upper threshold. This step ensures mutual neighbor sharing and thus removes edges which are likely due to spurious correlations.

To group genes according to this graph, typical community detection approaches may target high graph modularity, requiring communities to represent non-overlapping sets of nodes (*118, 119*) which is not ideal for our setting. To allow gene sharing between modules, we applied the Order Statistics Local Optimization Method (OSLOM) algorithm (*120*) for overlapping community detection on the Jaccard-neighbor graph. Briefly, the method clusters an undirected graph by optimizing a measure of cluster significance against a random null network lacking community structure. We applied the entire module-detection approach separately to the epithelial pre-malignant cells and matched samples from the immune data split into (1) T cell, NK cell and innate lymphoid cell (ILC), (2) myeloid cell, and (3) B cell subsets (**Table S13**) (**Fig. S8B,C**). These groupings allowed the discovery of more refined modules which correspond with co-expression patterns in rarer immune subsets, including Tregs and plasma cells. Hence, we obtained four sets of communication modules which we term epithelial, T/NK/ILC, myeloid, and B cell sets. Finally, we inspected each module to assess the biological plausibility based on knowledge of cell-state programs. As we occasionally observed nonsensical, lowly expressed genes in otherwise coherent modules (e.g. Natural Killer lectin -like receptors in epithelial modules), we added a filtering step to remove genes expressed in <10 cells in each subset. Performing this filtering post-module inference provides an interpretability benefit, as we were able to assess its impact on module plausibility guided by biological knowledge of gene groupings.

##### Association of modules with cell state

To utilize our communication modules for cell-cell signaling inference, we sought to associate cell-states with their expressed modules. As an initial step, we devised a strategy to quantify and visualize relative module expression per cell (**Fig. 4B**). We first defined a color-coding matrix of dimension *M x* 3, where *M* is the number of modules, and each row contains a user-defined, 3-dimensional RGB code that specifies a unique color for its corresponding module. We next assigned a single RGB color to each cell, based on its module usage, as a linear combination of module-specific colors, akin to “pseudo-coloring” used for microscopy images. Specifically, the matrix product of the color-coding matrix with an M *x* N matrix containing the average module expression in each cell (scaled between 0 and 1) gives a pseudo-coloring of each cell by its relative module expression. Cells with low expression for all modules will be close to black (R = 0, G = 0, B = 0); cells strongly skewed toward one module will assume a color value similar to that module’s code; and cells that highly express two or more modules will assume a hue that is intermediate between module colors. Hence, the predominantly monochrome, module-specific color distribution in the FDL visualization of pre-malignant cells (**Fig. 4B**) implies a strong degree of mutual exclusivity in communication module expression across the epithelia.

The partition of module activity across epithelia suggested that it would be reasonable to annotate each epithelial communication module based on its prevalence within a cell-state. We generated hard assignments for each cell based on the communication module with highest average expression (z-scored across cells for comparable scale), and confirmed that a visualization of the assignments mirrored the module pseudo-coloring (**Fig. S6B**). Checking the extent to which these assignments intersected cell-state definitions computed using Phenograph on the full dataset (see Progression Cohort coarse clustering in <u>Single-cell RNA-seq data processing</u>), we found that cells of sizable clusters (>200 cells) were largely assigned to a single module, with 62% of cells in each cluster, on average, assigned to the predominant communication module (**Fig. S6C**). Further, the Rand index between clustering and module assignment was relatively strong at 0.35. Given this degree of pairing between cell-states and communication modules, we were able to annotate each module based on the most closely associated cell-state (i.e., which cluster the cells associated with a module most frequently originate from; see **Fig. S8C** for immune module visualization). One epithelial communication module had low expression across all major populations, and was not annotated. We repeated this process of module annotation separately on the three major immune subsets (**Fig. S8C**).

##### Comparison of pre-malignant epithelial modules to normal pancreas and cancer

The correspondence between our modules and cell-states suggests that communication module expression can largely explain transcriptional heterogeneity in the pre-malignant epithelia. We asked whether this also holds in normal pancreas, normal regeneration and late-stage malignant disease. To this end, we computed average pre-malignant communication module expression in N1-N2 cells and K5-K6 cells, and visualized module distribution using the pseudo-coloring approach above (**Figs. 4C and 4D**). The majority of normal regeneration (N1, N2) cells exhibited low expression across all communication modules (dark pseudo-coloring), suggesting that pre-malignant inflammatory modules are largely inactive outside the oncogenic context. In contrast, malignant cells (K5, K6) expressed high levels of Progenitor, Gastric, and Bridge communication modules.

We additionally integrated our mouse-derived communication modules with publicly available human data (*32*) processed as described above. For each module defined in the pre-malignant pancreas, we computed the log of the average expression of each homologous gene across all human epithelial cells. **Fig. 4E** displays distributions of these scores across cells per module. This reveals that Progenitor, Gastric, and Bridge modules are up-regulated in human PDAC cells, consistent with the notion that these modules persist in advanced murine tumors.

##### Gene-centric module crosstalk algorithm

The communication modules define inflammatory programs that are differentially expressed across cell-states and are largely unique to the pre-malignant context. We next sought to identify module crosstalk that represents heterotypic communication driven by tissue inflammation in the context of oncogene activation. We first filtered the module graph described above, removing nodes which are not differential between mutant *Kras*-associated injury (K2) and regeneration (N2). Specifically, we identified genes upregulated in K2 compared to N2 from bulk RNA-seq data published in (*25*), retaining those with DESeq logFC > 2 and with adjusted P-value < 0.05. We also retained nodes which are cognate pairs of these dynamic genes, to capture potential cell-cell interactions and downstream effects of these differential programs. In total, 55 receptors and 46 ligands remained out of 340 initial candidates. For all crosstalk analysis, manually curated R-L cognate genes were extracted from (*80*) and completed with additional PDAC/immune-relevant R-L pairs from literature (**Table S16**).

Next, we modified the graph to include *directed* edges between ligands and their cognate receptors (rather than edges between co-expressed genes), representing potentially interacting molecules involved in cell-cell communication between modules. Similar to previous methods which identify crosstalk events between *cell-states*, in this *gene-centric* approach, we consider two modules to be potentially interacting if they share many cognate R-L pairs. As such, we enumerate the number of cognate interactions that occur between all genes in each pair of interacting modules. To identify only module pairs whose counts are higher than chance, we compute a random null distribution *R* on the pairwise interaction counts by shuffling the module labels for each gene and re-computing the counts for n = 5000 trials. We then compute empirical p-values for each interacting pair as 1 - p(*R* < observed) to identify significant interactions, akin to the procedure in CellPhoneDB (*61*) (**Fig. S9A**). Resulting networks are visualized in **Figs. S9B,C**, where each node represents one module and weighted edges represent statistically significant module-module interactions inferred by Calligraphy.

##### Per-module sender and receiver score

Our crosstalk inference approach provided a comprehensive map of candidate cell-cell interactions across communication modules and their associated cell-states. We visualized these interactions as a graph with modules as nodes and edge weights indicating strength of interaction (number of edges between modules) (**Fig. S9B,C**). We reasoned that modules with numerous interactions across epithelial and immune subtypes represent central (‘hub’) communicators of injury-driven neoplasia, and sought to quantify this notion. For each module, we summed counts of outgoing and incoming edges in the full gene-gene interaction network (filtered for statistically significant interactions) to quantify a “sensing” and “re-modeling” score, respectively. Visualization of these scores as bar charts along a heatmap of pairwise significant module-module interactions highlights modules with high sending and/or receiving propensity (**Fig. 4F**).

##### Analysis of sequential paths through crosstalk network

Sensing and re-modeling scores annotate important communication modules with substantial pairwise interactions between cell-states. However, many consequential intercellular events may involve stepwise interaction between more than two epithelial or immune populations. For instance, *feedback loops*, wherein a module signals back to itself through one or more intermediate populations, have been identified in late-stage cancer (*121*). To identify putative feedback loops in our Calligraphy network, we performed a search for cycles using networkx’s simple_cycles function. This identified a single loop involving Gastric module (E6) and Treg/ILC module (T8).

A major goal toward translating cell-cell communication networks to actionable targets is the identification of molecules whose expression has widespread impacts on downstream targets. To determine the relative impact of any one ligand within our feedback loop, we computed hierarchical paths beginning with a single “source” ligand, downstream to its immediate binding partners, and from each of these sequentially through all possible paths in the Calligraphy network. For computational efficiency, we began this search by identifying “sink” modules in the network which have no significant outgoing edges and hence represent dead-end absorbing states in a walk on the graph from any origin. For a particular source ligand contained in a particular module, we identified all its receiving modules containing its cognate binding partners as inferred by Calligraphy. We then rapidly built the downstream hierarchy of all possible paths emanating from these downstream modules by applying networkx’s all_simple_paths function to enumerate paths of all possible lengths to each sink. We visualized paths using plotly’s sunburst function, annotating the inner circle with source ligand and each outward layer as a possible step along the path hierarchy (**Fig. 5D**).

To then determine the impact of the source ligand on any given module, we annotated each downstream module by the level of its earliest appearance in the hierarchy. Assuming that communications occurring with fewer intermediates are most likely and/or most potent, these scores signify the putative impact of the source ligand (with lower scores suggesting stronger impact). To then associate these module impact scores to cell-states, we annotate each cell by its most highly expressed module as described above (see *Association of modules with cell state*). Transfer of module impact scores to their associated cells allowed us to assess the breadth of phenotypes which may be affected by expression of a given ligand (**Fig. 5E**).

#### Single-cell analysis of KC-shRNA cohorts

Analysis of ligand-specific paths in our crosstalk network revealed an IL-33-driven feedback loop involving pre-malignant populations and their microenvironment. To determine the impact of this crosstalk on epithelial cells, we modeled phenotypic shifts in each stage and tissue type of our Perturbation Cohort, comparing sh*Il33* to control samples with the Milo algorithm (*74*). Milo groups cells into partially overlapping local neighborhoods on a kNN graph, and then computes statistics for differential neighborhood abundance across conditions using a negative binomial generalized linear model (GLM). We applied Milo with k = 20 for neighborhood detection and took advantage of its GLM to further account for batch confounding by modeling the experimental mouse cohort as a covariate. We visualized the distribution of log fold changes in each analysis (K2 epithelial and immune, and K3 epithelial and immune) separately as a measure of degree of perturbation in each condition (**Fig. 6E**).

K3 epithelium displayed the strongest and most consistent phenotype for further exploration of cell-states impacted by perturbation. We were primarily interested in understanding whether our perturbation alters the natural progression of disease. To model an axis of progression in this time point, we applied pseudotime inference with Palantir (*75*) to neighborhoods derived from Milo. Palantir requires assignment of a starting cell in order to seed the direction of pseudotime. In our case, we chose to use *Nes*^+^ progenitor cells as the primary cell of origin, supported by CellRank’s identification of this state as an initiating population (**Fig. S3A**) as well as multiple previously described measures suggesting the potential of this population to give rise to downstream cell states (**Figs. S3E** and **3F**).

To identify a *Nes*^+^ progenitor neighborhood which could serve as a starting cell for pseudotime, we first annotated each neighborhood in Milo using the annotateNhoods function with cell-state annotations assigned to each original single cell as described above for the Progression Cohort (see <u>Single-cell RNA-seq data processing and basic analysis</u> for list of marker genes) (**Fig. S11C**). We then computed diffusion maps with Palantir (n_components=10) to identify major axes of variation through the neighborhoods. We selected a starting cell at the extreme of the second diffusion component, which tracked from a *Nes*^+^ progenitor to downstream cell types. Palantir was run with 500 waypoints to identify an axis of pseudotime from this starting position (**Fig. S11C**). In **Fig. 6G**, we visualize the Milo differential abundance with neighborhoods sorted along this axis, finding increased abundance in sh*Il33* for the latest subset of *Nes*^+^ progenitors.

We further find that this population maintains expression of genes highly correlated with the plasticity score across epigenomic metacells (Pearson correlation Bonferroni corrected p-value < .01). Finally, to interpret these results in light of our inferred module interaction networks, we evaluated the extent to which cell-states predicted to be downstream of IL-33-centric crosstalk pathways overlap with those impacted by the perturbation. To this end, we divided epithelial modules into ‘connected’ modules which are directly or indirectly downstream of IL-33 in the network and ‘unconnected’ modules, which are not. We reasoned that connected modules are more likely to be impacted by perturbation, provided that our module interaction network captures true interactions driven by this cytokine. To utilize neighborhood-specific scores from Milo as a measure of these module-level impacts, we first associated modules with neighborhoods, similar to our approach in <u>Heterotypic cell-cell communication</u>. Specifically, we computed hard module assignments for each neighborhood based on the module with the highest average z-scored expression (**Fig. S11D**). We then compared the average absolute value Milo log fold change in all neighborhoods assigned to IL-33-connected modules versus unconnected modules, finding significantly higher perturbation impact in connected modules (one-sided t-test; t = -4.6711, *P* = 2.0551 × 10^-6^), and thus experimentally validating connections in the inferred IL-33 crosstalk pathway (**Fig. 6H**).

---

**Figure S1.**
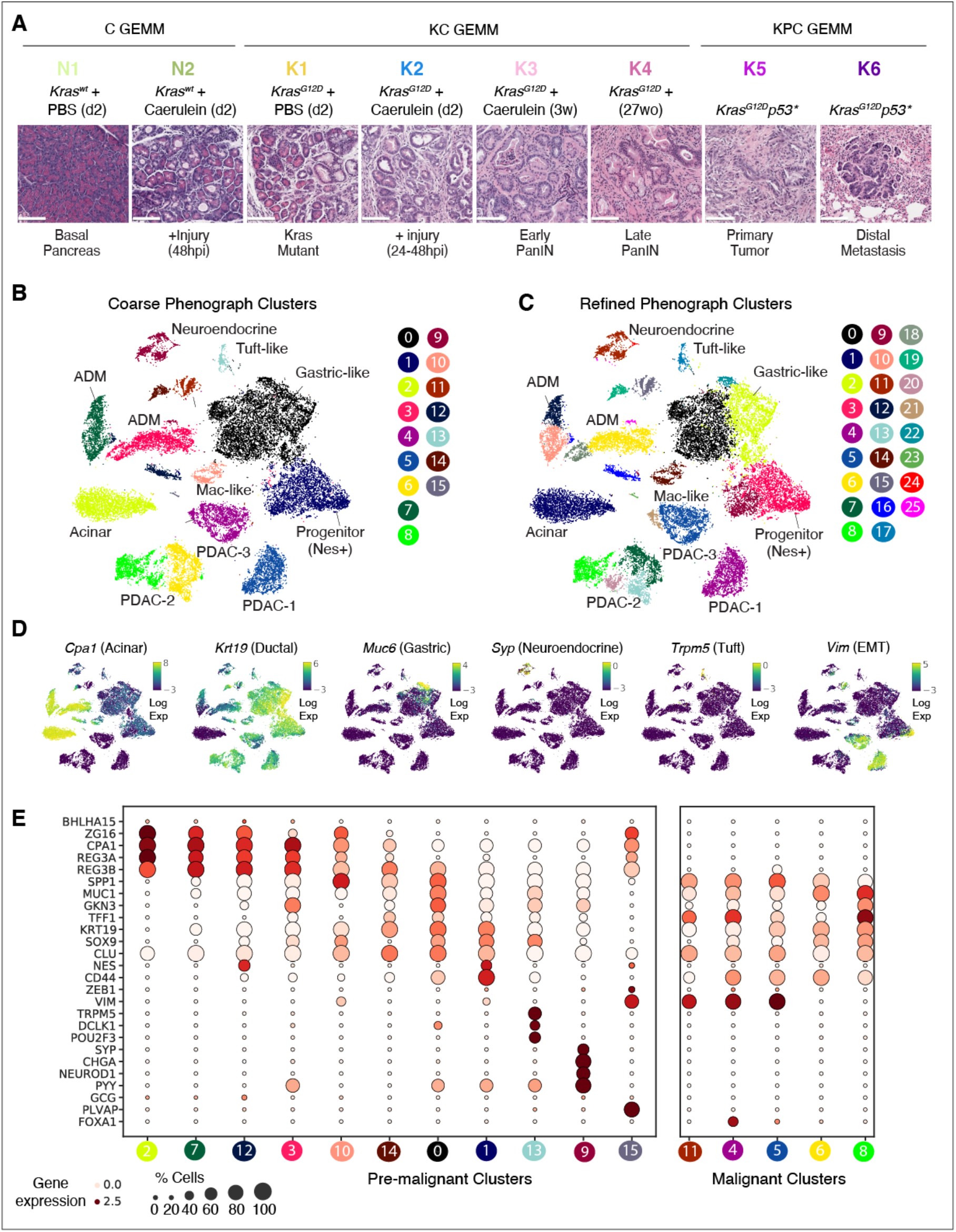
Intra- and inter-tissue epithelial heterogeneity and dynamics during cancer progression. (A) Experimental settings to interrogate pancreatic epithelial heterogeneity *in vivo*. Representative H&E of pancreata from the indicated mouse models and treatment conditions (as in Fig. 1A), for which mKate2+ pancreatic epithelial cells were isolated and subjected to transcriptional or chromatin accessibility profiling. (B) tSNE visualization of epithelial (mKate^+^) scRNA-seq profiles from all collected settings in (A), colored by coarse cluster membership computed with Phenograph (*101*); clusters are annotated manually (**Methods**). ADM denotes cells actively undergoing acinar-to-ductal metaplasia in *Kras*-wild-type or mutant tissue (*20*). (C) tSNE map as in (B), colored by fine cluster membership computed with Phenograph. (D) tSNE map as in (B), colored by log-normalized pancreatic epithelial cell-type marker gene expression. Colors are scaled between the 1st and 99th percentile. (E) Expression of pancreatic epithelial cell-state markers (rows) across coarse Phenograph clusters (columns). Dot size scales with the proportion of cells in a given cluster that express each gene; color indicates average z-scored, log-normalized expression. Cells in pre-malignant conditions gradually lose expression of acinar-associated markers (*Bhlha15, Zg16, Cpa1*) and gain expression of metaplasia-associated markers (*Krt19, Sox9*). Other populations express distinct cell-state markers including *Syp*, *Neurod1*, and *Pyy* in neuroendocrine-like cells (cluster 9) and *Trpm5*, *Dclk1*, and *Pou2f3* in tuft cells (cluster 13).

**Figure S2.**
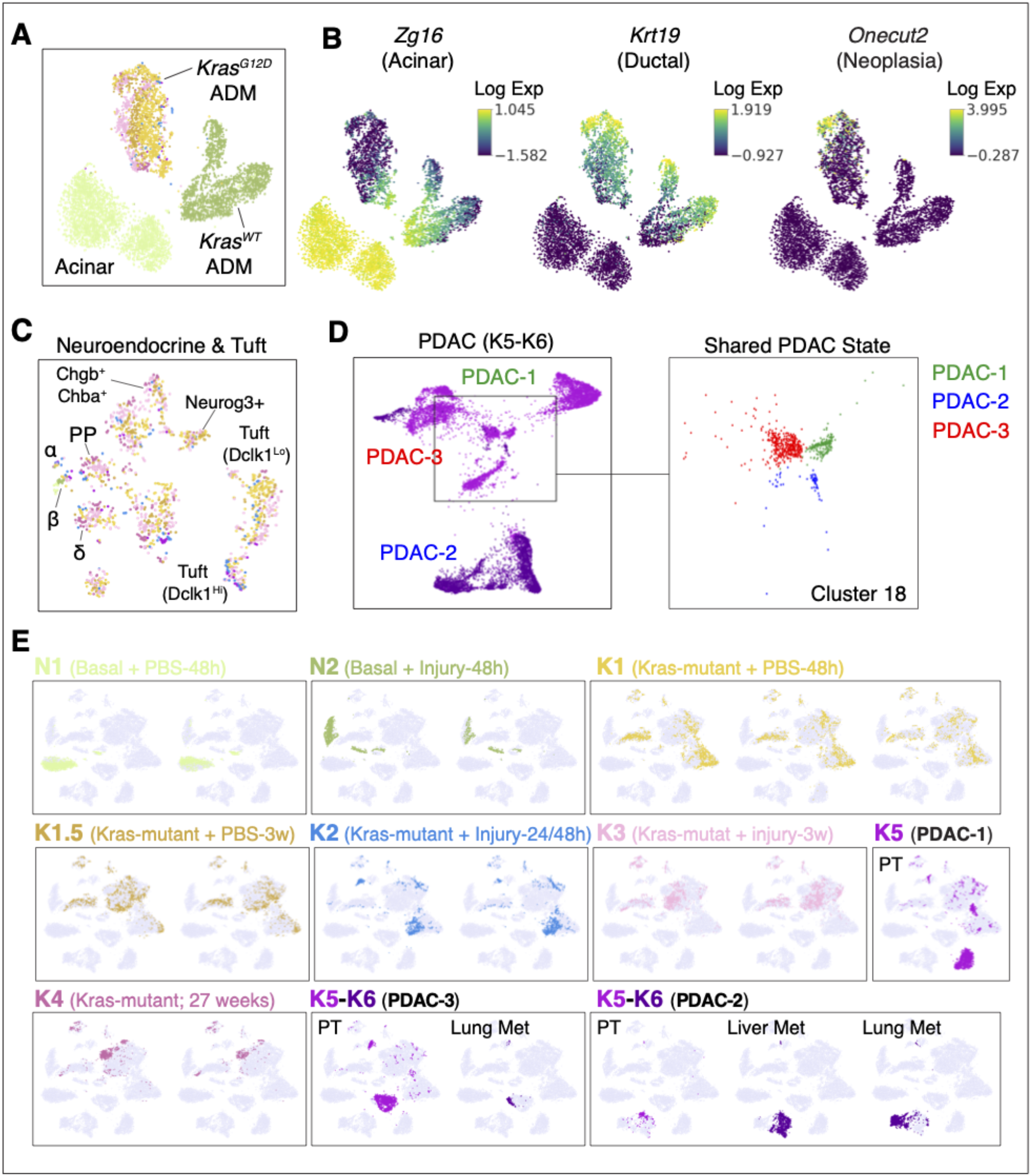
Resolution and reproducibility of pre-malignant epithelial cell states. (A) tSNE map of cells derived from coarse Phenograph clusters 2, 3, 7 and 12 (see **Fig. S1B**) undergoing ADM in regeneration (N1, N2) and tumor progression (K1–K6). Normal and oncogenic *Kras* conditions exhibit little overlap. (B) tSNE map as in (A), colored by log-normalized expression of acinar gene *Zg16*, which decreases across both *Kras-*wild-type and *Kras*-mutant ADM; ductal gene *Krt19*, which increases along both; and neoplasia-associated gene *Onecut2* (*77, 122*), which is uniquely activated along ADM in the *Kras* mutant. (C) tSNE map of neuroendocrine and tuft cells derived from coarse clusters 9 and 13, showing significant heterogeneity in *Kras*-mutant cells (largely absent from *Kras*-wild-type epithelia, which only contain an extremely rare population of beta -like cells) and rare cell populations captured through epithelial cell enrichment. Substantial mixing of cells across conditions further underscores reproducibility of transcriptomic profiles. (D) FDL of malignant tumor cells from full-blown PDAC samples and isogenic distal metastases (K5, K6) highlights phenotypic divergence between individual mice (PDAC-1 to -3), with the exception of a single shared state (cluster 18) from all 3 malignancies (inset). (E) tSNE map as in Fig. 1B, highlighting biological replicates (independent mice) for each condition (**Table S1)**. Each tSNE represents an individual mouse, colored to indicate constituent cells. Normal (N1), regenerating (N2) and pre-malignant (K1-K4) pancreatic epithelial cells are highly reproducible across replicates, whereas malignant replicates (K5, K6) diverge. Primary tumors and metastases derived from a single mouse are grouped together in one box, highlighting the phenotypic overlap in cells derived from the same primary tumor.

**Figure S3.**
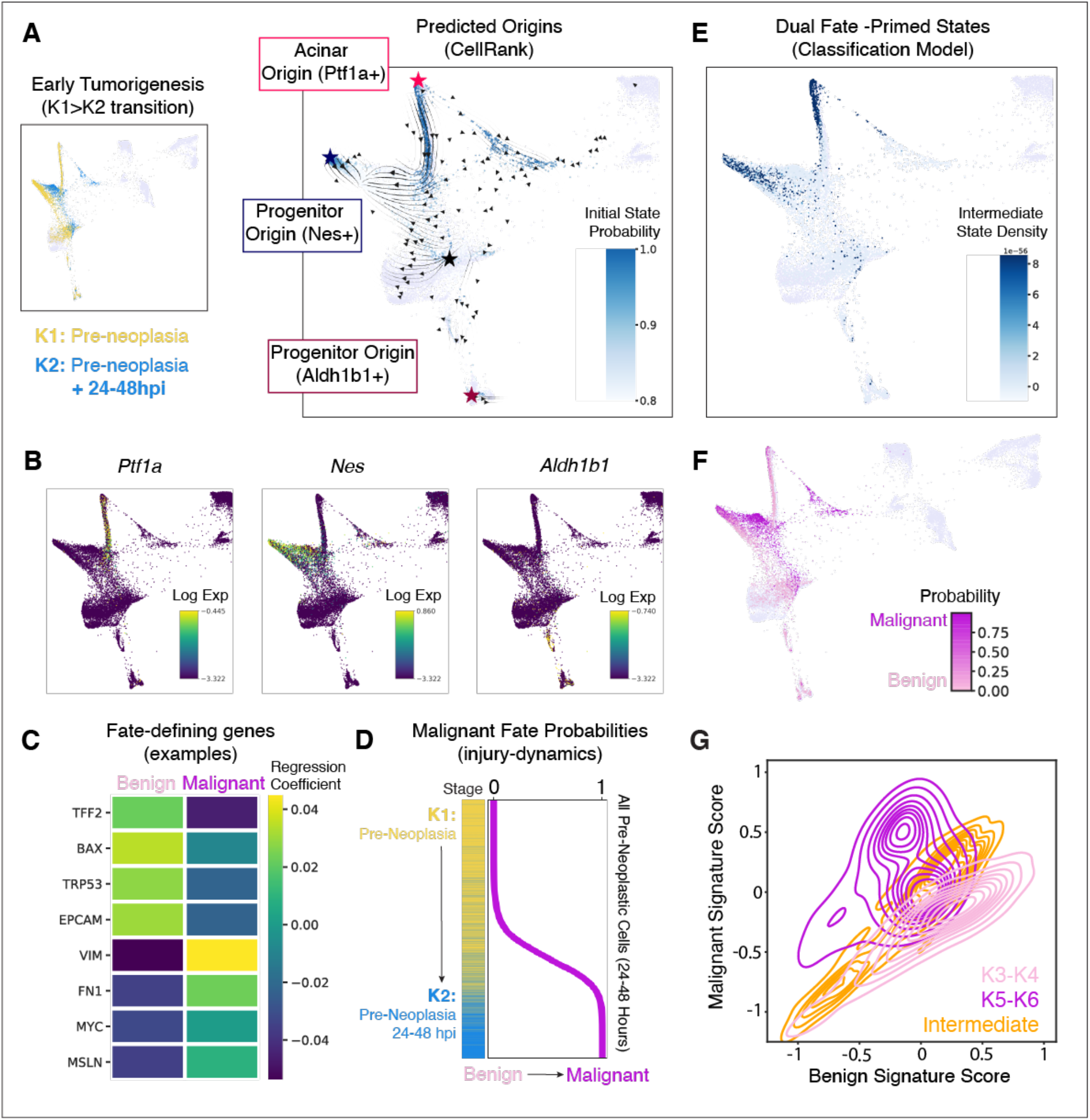
Identification of specific, injury-sensitive *Kras*-mutant subpopulations with multi-lineage potential. (A) CellRank (*47*) predicts multiple initiating populations of pancreatic tumorigenesis. Left, FDL of all *Kras*-mutant cells (K1–K6) as in Fig. 2A, colored to reveal pre-neoplastic cells (K1, K2). Right, region of the same FDL encompassing all K1 and K2 cells, overlaid with a streamplot of scVelo-inferred RNA velocity vectors (*48*); cells are colored by CellRank-computed probability of being an initiating state. High-probability initiating states are indicated with stars and correspond (by marker gene expression in (B)) to initiating populations identified by lineage tracing studies (*27, 50–53*). (B) Region of FDL in (A) colored by expression of known cell-of-origin marker genes (*27, 50– 53*). Populations expressing each gene roughly correspond to distinct initiating states inferred in an unbiased manner with CellRank. (C) Logistic regression model coefficients for Benign (left) or Malignant (right) fates for select genes. (D) Classification of all pre-neoplastic (K1, K2) cells to late-stage fates. Each row represents one cell from K1 (yellow) or K2 (blue) conditions, sorted from highest Benign probability (top) to highest Malignant probability (bottom). Corresponding Malignant fate probability is plotted at right. (E) FDL region as in (A), highlighting cells with an intermediate probability (0.4 < p < 0.6) of classifying to a Benign or Malignant phenotype. Cells are colored by the density of intermediate cells in the phenotypic space (darker corresponds to more intermediate cells in the local region). Density was derived using a Gaussian kernel density estimate of intermediate cells in PC space. (F) FDL as in (A), colored by probability of classification to Benign (pink) or Malignant (purple) fates. Shift to states with high Malignant probabilities mirrors the shifts induced with injury from K1 to K2 conditions as in (A). (G) Intermediate cells exhibit evidence of dual priming for divergent fates. Contour plots display cell density based on expression of Benign and Malignant gene signatures for benign neoplasia (K3, K4) cells (pink), malignant (K5, K6) cells (purple), and intermediate (composite state) cells from K1 and K2 identified by the classification model (orange). Gene signatures are derived by intersecting genes with the top 200 largest coefficients in the classification models for K3 and K5 classes with genes uniquely associated with bulk ATAC-seq benign and adenocarcinoma modules (see Fig. 2C and **Table S2**), respectively. A per-cell signature score was computed as the average z-scored expression of each gene in the signature. We observe that individual intermediate cells from K1 and K2 co-express benign and malignant programs.

**Figure S4.**
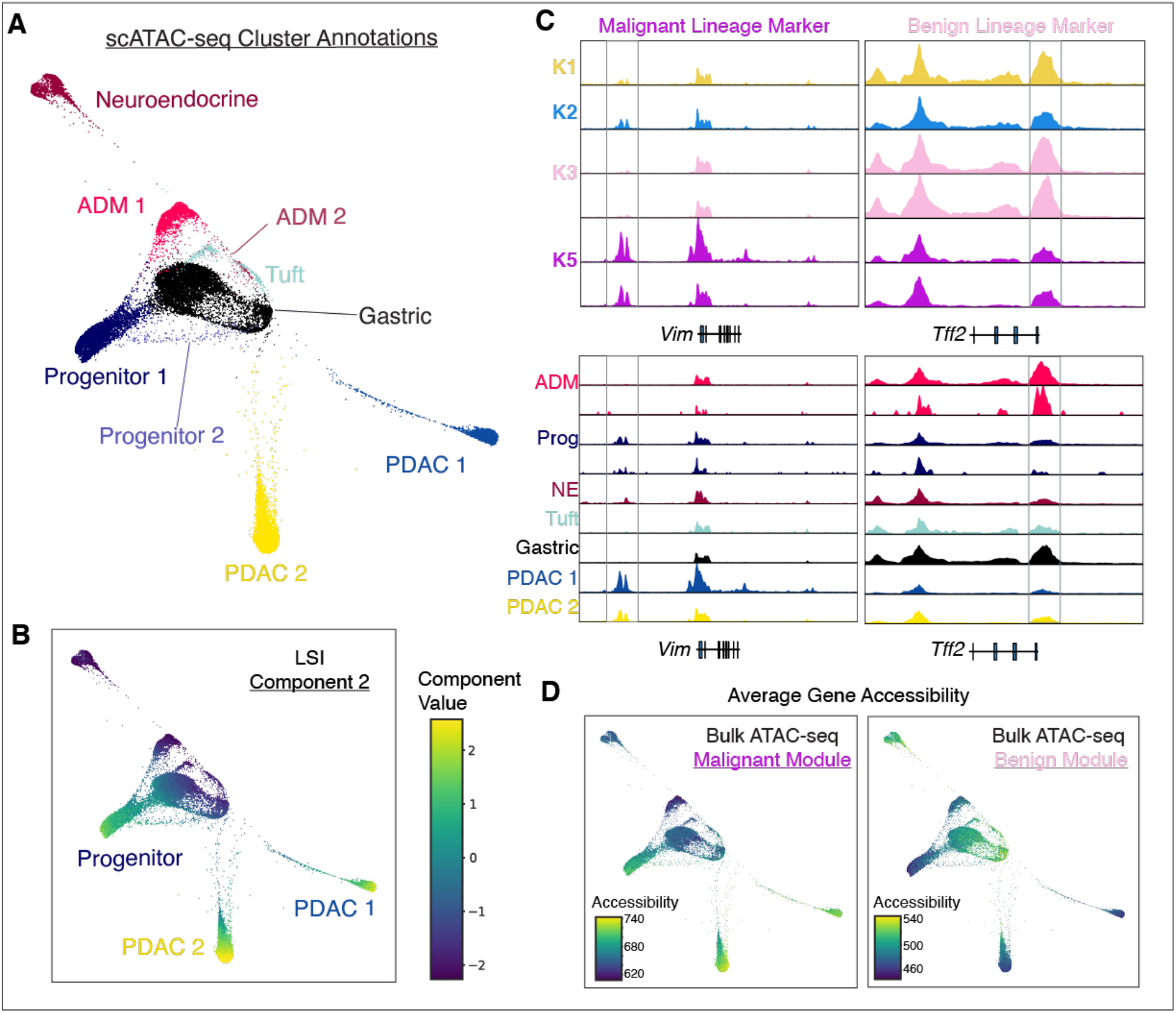
Epigenomic characterization of early and late *Kras*-mutant subpopulations. (A) FDL of scATAC-seq profiles from pre-malignant (K1-K3) and malignant (K5) stages, colored by merged Phenograph cluster and annotated manually (**Methods**). (B) FDL as in (A), colored by second component of Latent Semantic Indexing (LSI), revealing similarity between tumor and *Nes*^+^ progenitor states at the chromatin level. (C) Chromatin accessibility tracks for representative benign and adenocarcinoma-primed genes, aggregated across cells belonging to a given condition (top) or merged Phenograph cluster (bottom). x-axis, genomic coordinates around the indicated gene; y-axis, smoothed, depth-normalized counts of scATAC-seq fragments. *Tff2* is selected from the benign neoplasia classification model high-coefficient features (see **Fig. S2C**) and appears primed toward benign neoplasia in early tumorigenesis, whereas *Vim* is selected from the adenocarcinoma model and shows priming toward PDAC. (D) FDL of scATAC-seq profiles from pre-malignant and malignant stages, colored by sum of accessibility near genes (ArchR gene score) associated with Benign and Malignant chromatin modules (**Table S2**). Cells with high accessibility for these fate-associated genes fall within distinct regions of the map. Similar to patterns observed in Fig. 2C, Malignant programs are activated in *Nes*^+^ progenitor-like cells, where benign programs are enriched in gastric-like cells.

**Figure S5.**
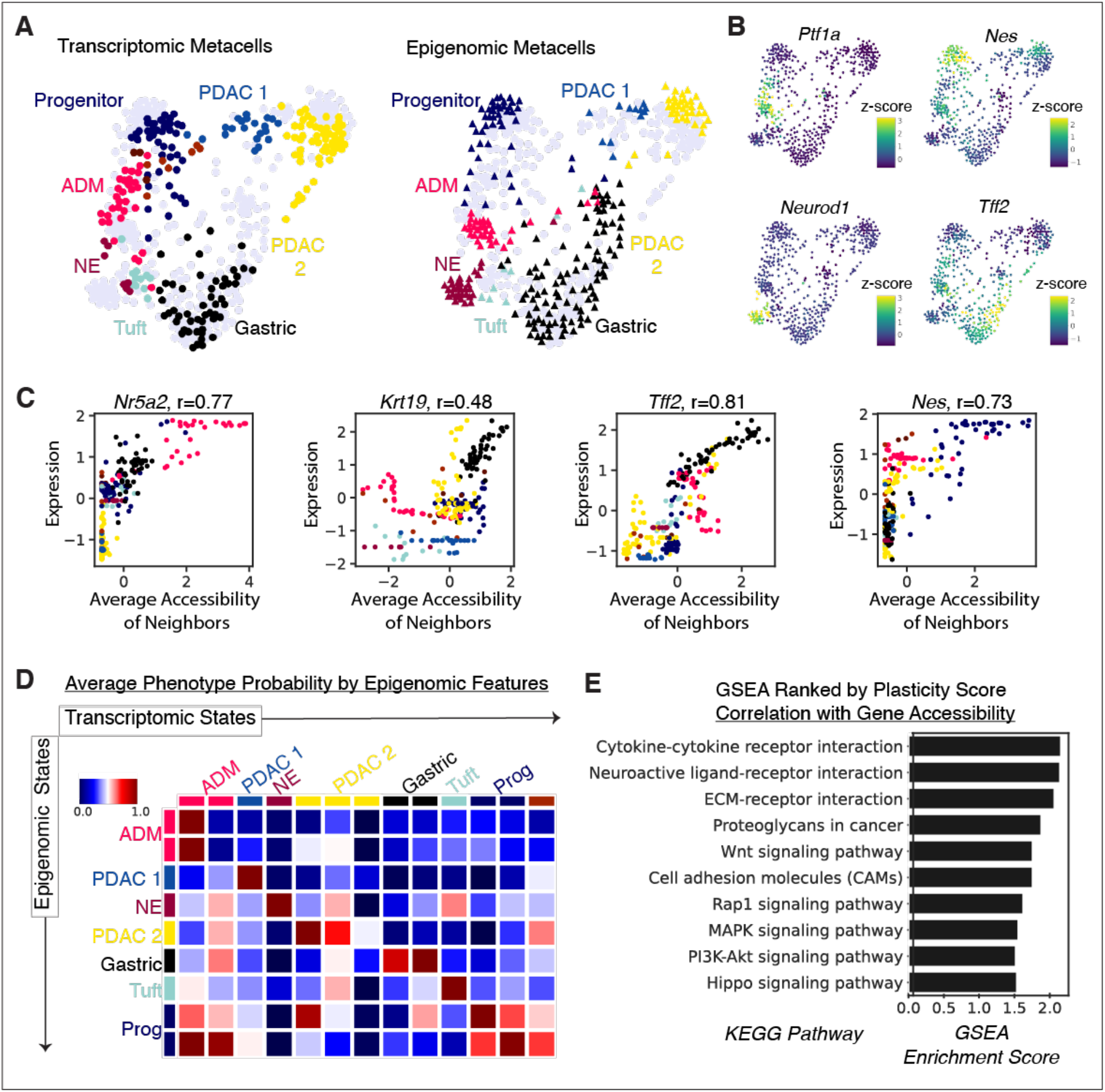
Integration of tumorigenesis-associated epigenomic and transcriptional cell states. (A) Integrated FDL visualization of metacells derived from scATAC-seq (triangles, right) and scRNA-seq (circles, left), with nodes colored by Phenograph cluster from each respective technology (**Figs. S1B** and **S4A**). The FDL is built on a composite graph which combines within-modality nearest neighbors with cross-modality mutually nearest neighbors (MNN) to emphasize similarities between cells both within and across modalities. (B) FDL as in (A), colored by expression score (the average of log-normalized counts from all cells in a metacell) or accessibility score (average of log-normalized ArchR gene accessibility scores) for each metacell, for a selection of known cell-state markers. Values are z-scored within each modality to obtain comparable scales. Acinar (*Ptf1a*), progenitor-like (*Nes*), neuroendocrine-like (*Neurod1*), and gastric-like (*Tff2*) markers show high concordance between modalities on the visualization. (C) Per-gene correlations of accessibility and expression across paired metacells. Each point corresponds to one transcriptomic metacell, colored by Phenograph cluster. Y-axis displays z-scored expression of the indicated gene in each metacell; x-axis displays the average z-scored accessibility score across the epigenomic metacell MNNs of that transcriptomic metacell. High correlation values indicate strong concordance between expression of a gene and its accessibility for matched epigenomic profiles. (D) Average phenotype probability by epigenomic features. Heatmap displays average log probability of classification to each transcriptomic cluster (columns) from metacells of each epigenomic cluster (rows). Row and column order correspond to that in the confusion matrix in Fig. 3E. Color values indicate the full log probability distribution (as opposed to the count of discrete predictions for cells of each epigenetic cluster depicted in Fig. 3E). (E) GSEA enrichment scores for select significantly (FDR < 0.1) enriched gene sets in genes ranked by Spearman correlation to inferred plasticity. High enrichment indicates a significant positive association of that program at the chromatin level with cell-states maintaining high plasticity as defined by the method in Fig. 3D.

**Figure S6.**
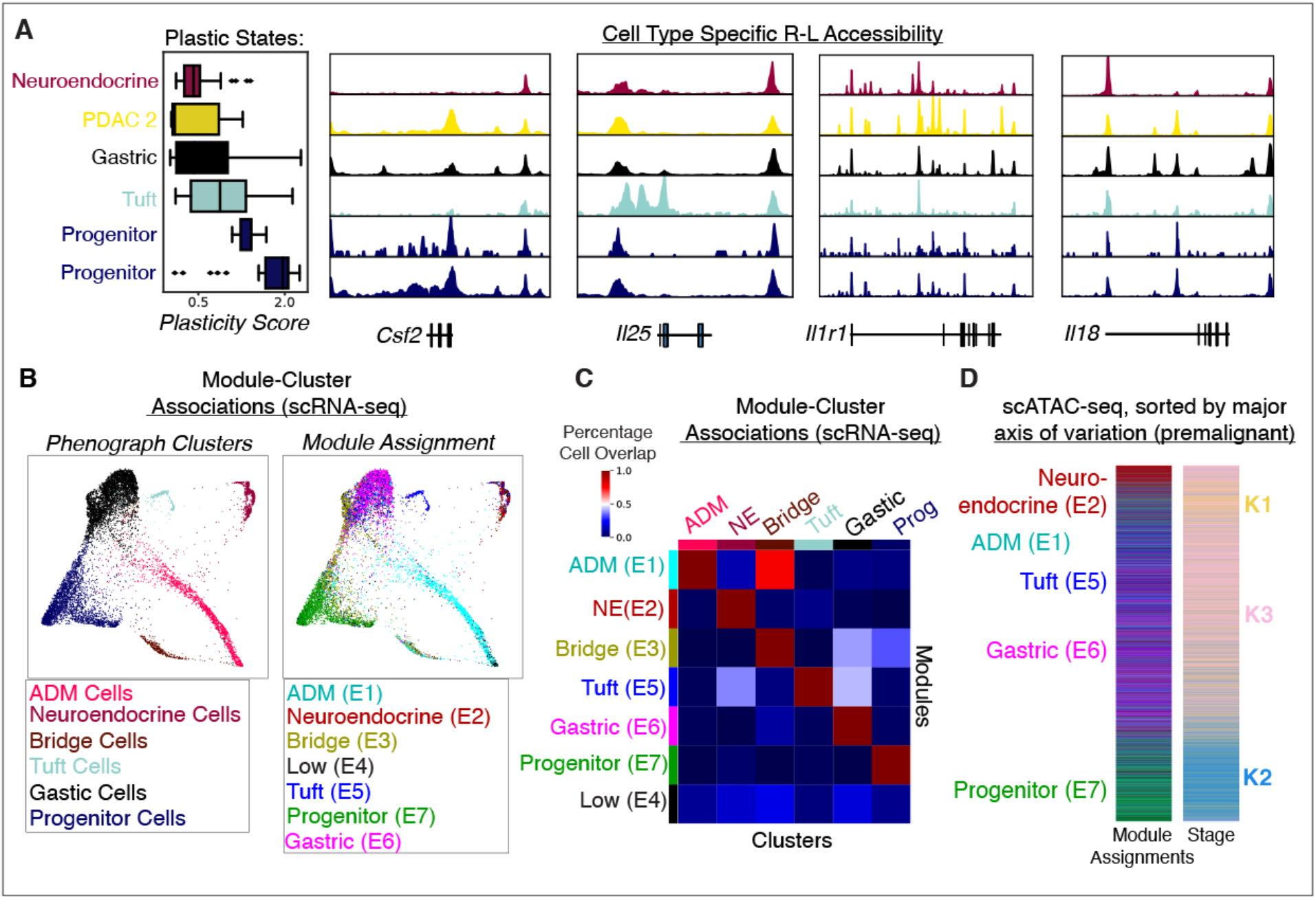
Distinct cell-cell communication modules are associated with defined neoplastic lineages. (A) Chromatin accessibility signal tracks from scATAC-seq data for select communication genes aggregated across cells from each Phenograph cluster of plastic cell-states. Clusters are ordered by increasing plasticity (reproduced at left here from **Fig. 3F**), highlighting greater accessibility near these genes in high-plasticity populations. (B) FDL of pre-malignant epithelial cells (K1–K3, see **Fig. 4B**), colored by coarse Phenograph cluster (left) or communication module assignment (right). Module expression is computed as the log of average normalized expression of each gene in that module; cells are assigned to the highest-expressed module. (C) Correspondence between cell membership in Phenograph clusters computed on all genes (columns) and communication modules computed only on communication genes (rows). Color values represent the proportion of cells in each cluster which map to each module, revealing tight concordance for all modules except Low (E4). (D) Communication module assignments of cells from chromatin accessibility data, based on average normalized accessibility (ArchR gene score) of communication genes within each module (**Methods**). Cells are ordered along the second component from LSI computed on scATAC-seq data.

**Figure S7.**
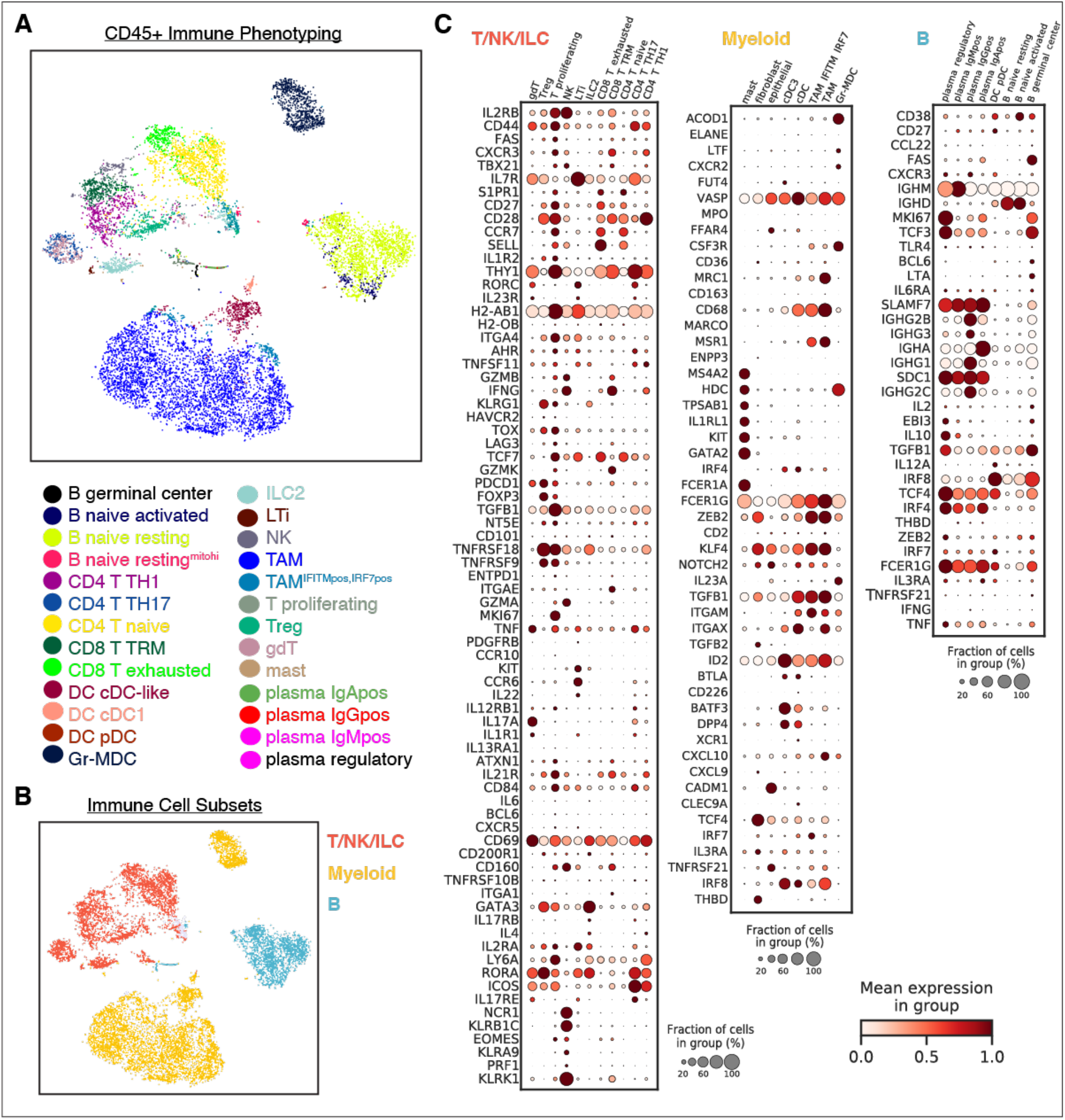
Characterization of immune heterogeneity in pancreatic tissue. (A) tSNE visualization (scRNA-seq) of CD45^+^-sorted immune populations from pre-malignant pancreata (K1–K3) colored by cell type annotation. (B) tSNE as in (A) displaying coarse immune cell subsets combining all T, NK, and ILC - derived cells, myeloid -derived cells, or B cell -derived cells into a group for downstream co-expression analysis and module determination (see **Figs. S8B,C**). (C) Immune marker expression (rows) across Phenograph clusters from CD45^+^-sorted immune cells in scRNA-seq (columns). Dot plots represent clusters derived from T/NK/ILC, Myeloid and B cell subsets from left to right.

**Figure S8.**
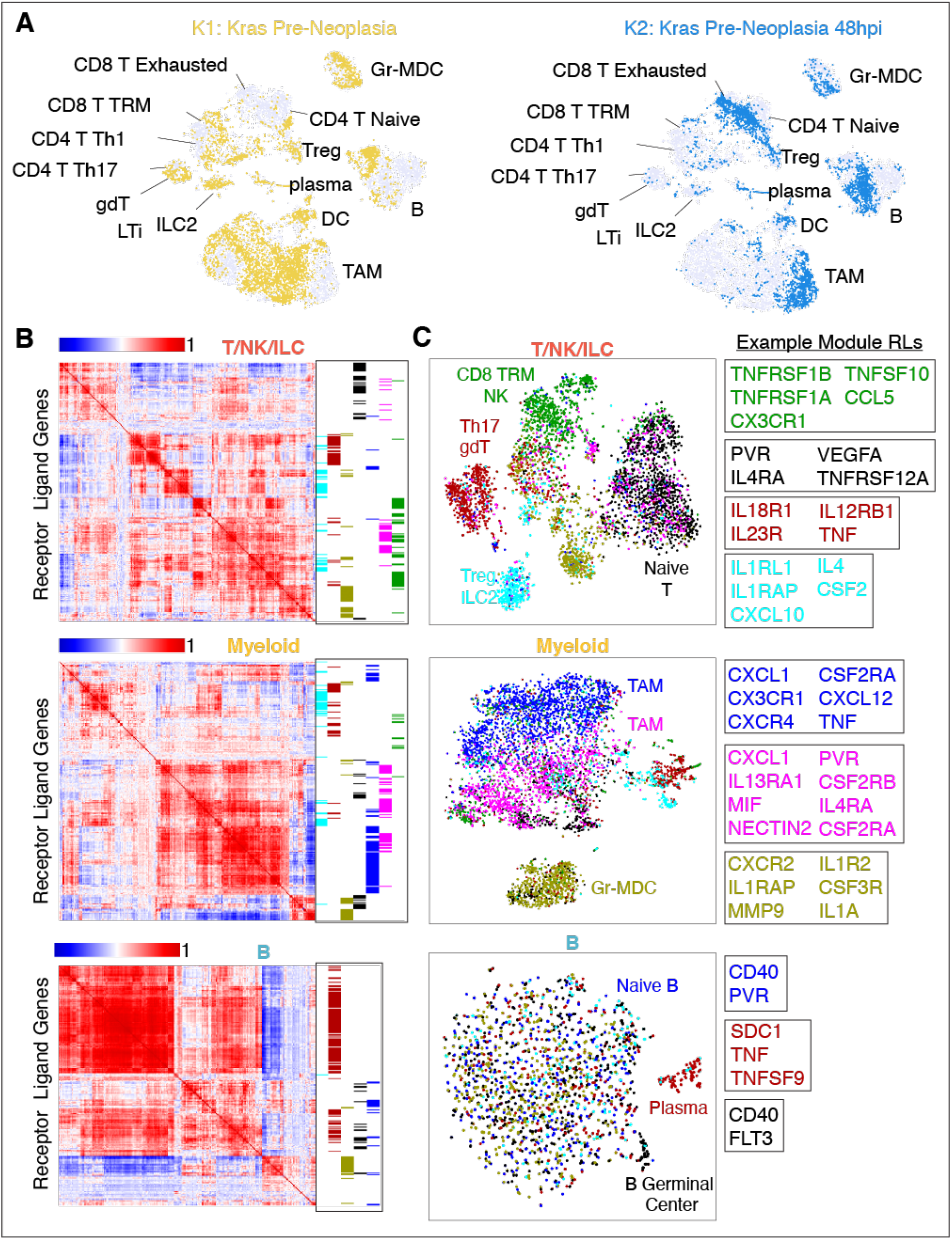
Communication module prediction in infiltrating immune cells. (A) tSNE visualization of CD45^+^-sorted immune populations from pre-malignant samples (K1– K3, see **Figs. S7A,B**). Coloring K1 (yellow) and K2 (blue) cells separately reveals the dramatic shifts in immune phenotypes that occur in *Kras*-mutant pancreata upon injury. (B) Communication gene co-expression modules in CD45^+^ immune cells derived from premalignant tissues separated into three major immune subsets as in **Fig. S7B**. Each row or column corresponds to one receptor or ligand, and color values represent the Pearson correlation coefficient between the expression of a pair of genes across cells of that subset. Blocks of highly correlated communication genes along the diagonal correspond to partially overlapping modules of genes that tend to be expressed in the same cell populations. Each column of inferred communication gene co-expression modules from the OSLOM community detection algorithm (right) depicts genes belonging to a single module. (C) tSNE visualization of major immune subsets in **Fig. S7B** with cells colored by module assignment from **Fig. S8B**. For each subset, modules are computed using OSLOM community detection (**Methods**), cells are assigned to the module with highest average z-scored, log-normalized expression. Right, representative communication genes from select modules. Individual genes can be shared between modules (e.g. CD40 in B cells) when genes are expressed in more than one cell type.

**Figure S9.**
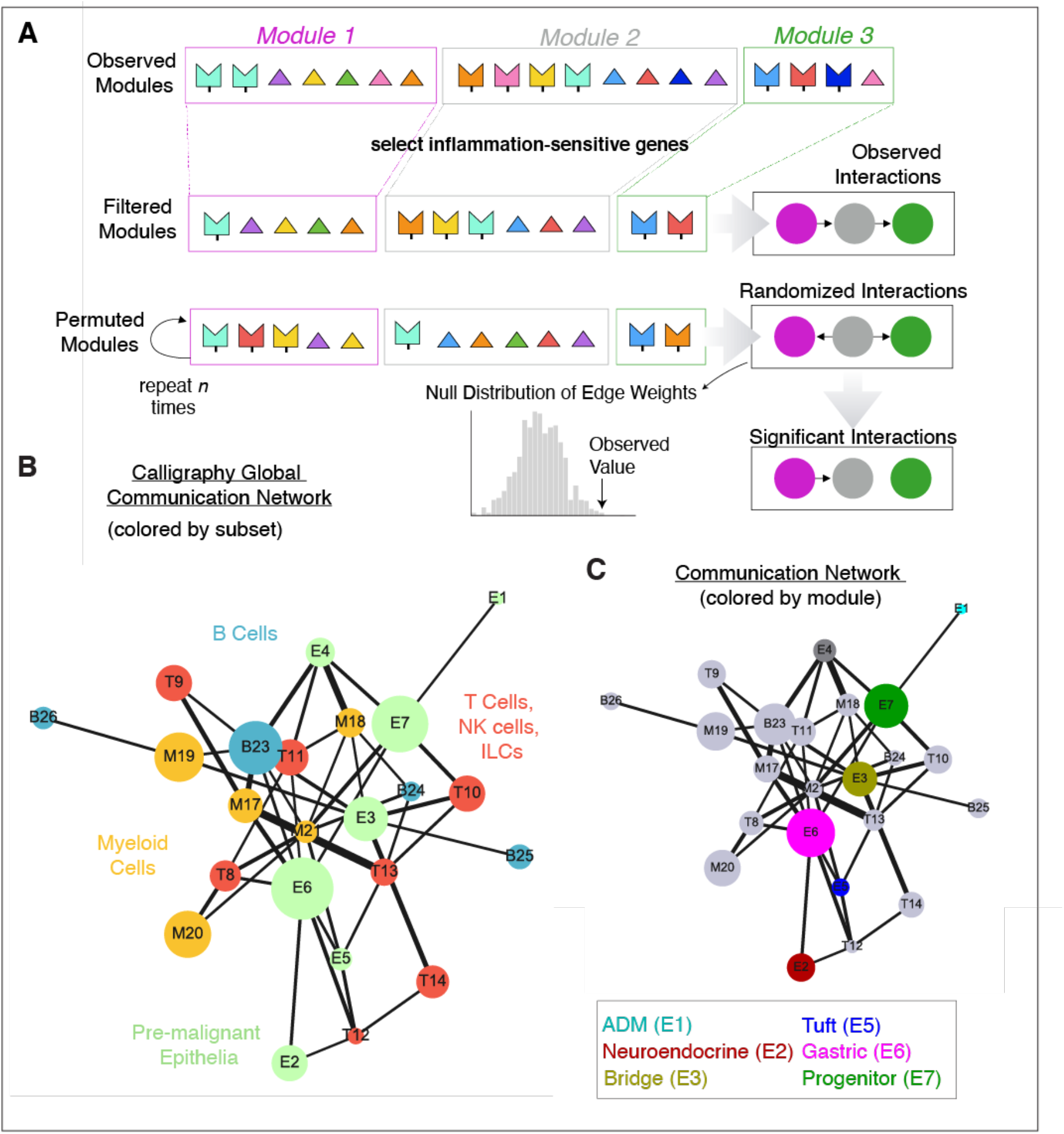
Calligraphy predicts tissue crosstalk networks between *Kras*-mutant cells and infiltrating immune cells recruited to their tissue environment. (A) Schematic of the module-based crosstalk algorithm, Calligraphy. Observed modules are first identified from a gene-gene co-expression graph of cell subpopulations using the OSLOM community-based detection approach. Each module contains a set of receptors (boxes) and ligands (triangles) which are co-expressed across subpopulations of scRNA-seq data, colored by cognate pairs known to physically interact. Inferred gene modules are then filtered for communication genes that are significantly upregulated in early tumorigenesis (K2) compared to normal regeneration (N2). Cognate R-L pairs spanning filtered modules are enumerated to suggest potential cross-module interactions. Then, modules are randomly permuted *n* times and cross-module interactions are recounted in each trial to derive a null distribution for each pairwise module-module interaction. A *P*-value is obtained for each pair of modules using this null distribution. (B) Global crosstalk network inferred by Calligraphy, colored by cell subset. Each node represents one module, with size proportional to the number of communication genes in that module. Edges connect significant module-module interactions, with widths proportional to the significance of interaction (–10 log(*P*-value + pseudocount)). (C) Global crosstalk network inferred by Calligraphy, colored by epithelial module as in Fig. 4A, or gray for immune modules.

**Figure S10.**
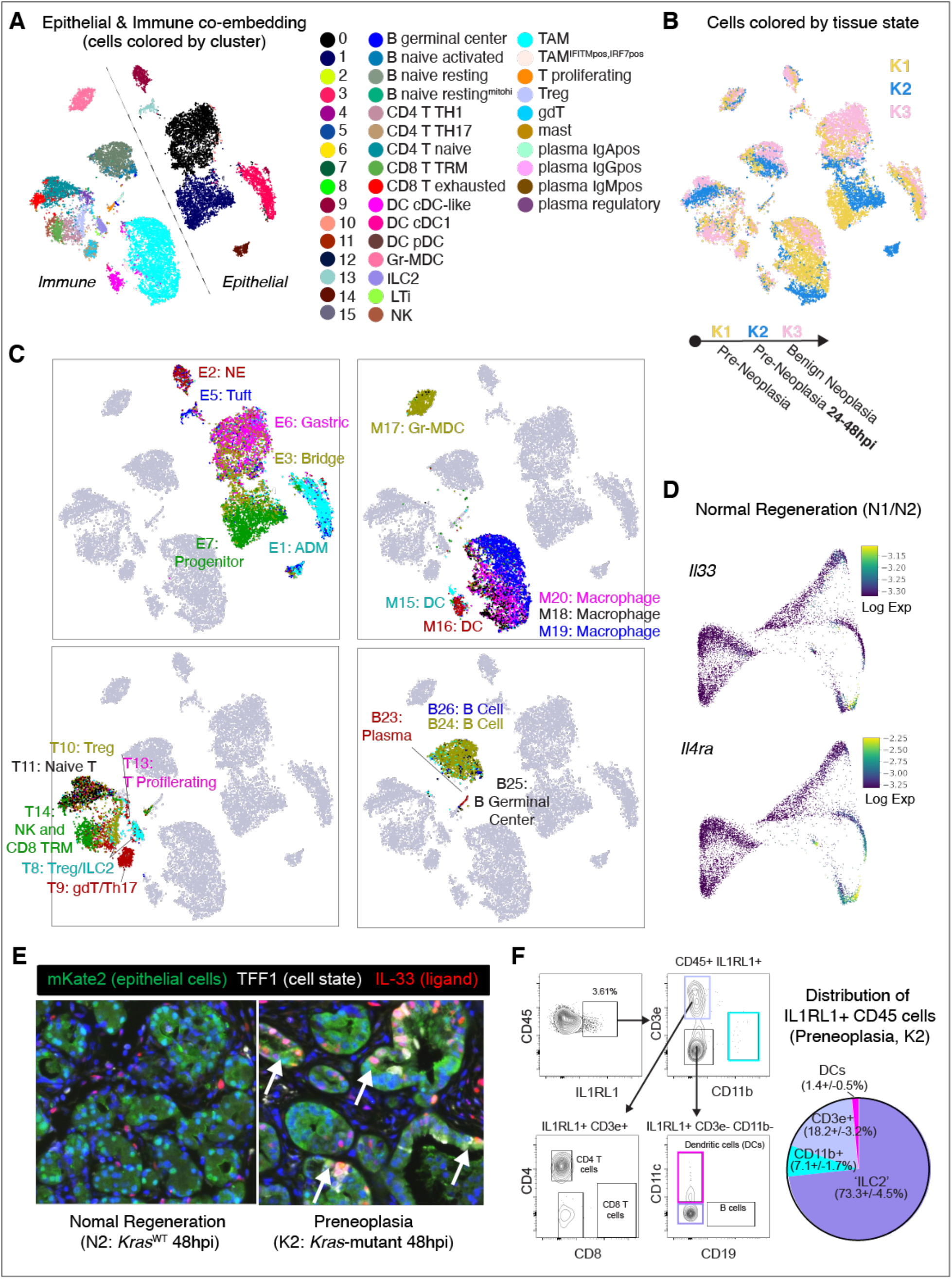
Identification of *Kras*-mutant specific epithelial-immune crosstalk networks. (A) tSNE of integrated immune and epithelial scRNA-seq data from pre-malignant stages (K1-K3, see Fig. 5B) colored by Phenograph cluster. (B) tSNE as in (A), colored by progression stage. (C) tSNE as in (A), separated by epithelial (top left), myeloid (top right), T/NK/ILC (bottom left), and B cell (bottom right) compartments. Cells of each compartment are colored in each plot by their module assignments. Module expression is computed as the log of average normalized expression of each gene in that module; cells are assigned to the highest-expressed module of the corresponding subset. (D) FDL of normal regeneration (N1, N2, see Fig. 4C) colored by MAGIC imputed log-normalized expression of *Il33* (middle) or *Il4ra* (bottom). Colors are scaled between the 1^st^ and 99^th^ percentile. (E) Representative immunofluorescence of IL-33 (red), TFF1 (white) or mKate2 (epithelial cells, green), visualizing the selective injury-driven induction of IL-33 in a subset of *Kras*-mutant pancreatic epithelial cells expressing the gastric-cluster marker TFF1 (right) but not in *Kras*-wild-type injured counterpart (left). (F) Representative FACS plots indicating gating strategy to characterize IL1RL1 positivity in myeloid and lymphoid compartments of Kras-mutant pancreata at 48hpi (left). The fraction of lymphoid and myeloid cells among IL1RL1^+^ immune cells in *Kras*-mutant pancreata at 48 hpi are quantified in the pie chart, consistent with the expected positivity in CD4^+^ T-reg cells (CD3^+^, CD11b^−^), DCs (CD11c^+^CD3^−^, CD11b^−^), Non DC-myeloid (CD11b^+^CD3^-^CD11c^−^), and ILC2 (CD3^−^, CD11b^−^, CD11c^−^). Pooled data are presented as mean ± s.e.m (n = 6 independent mice).

**Figure S11.**
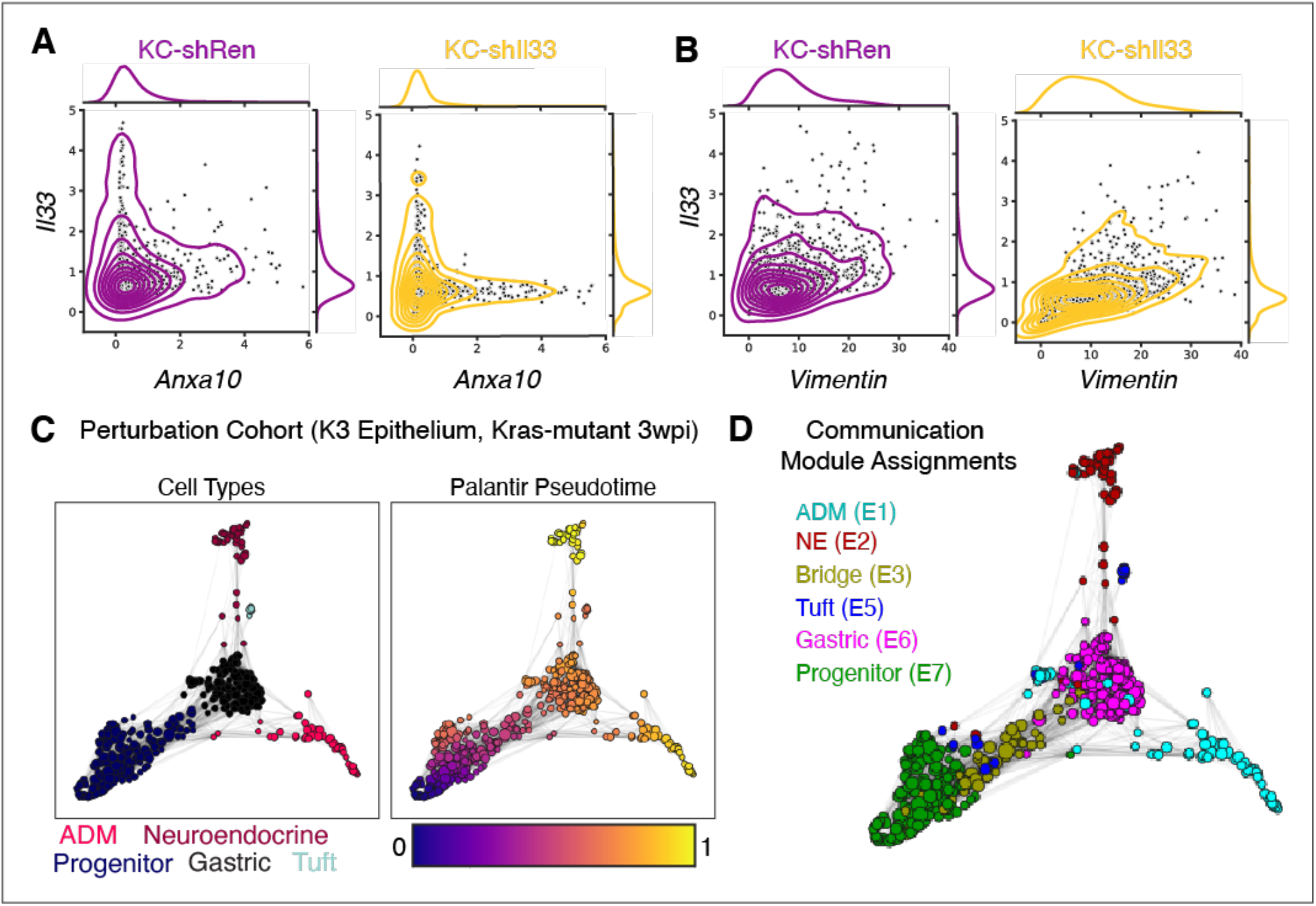
Impacts of *Il33* perturbation assessed with IMC and scRNA-seq. (A) Scatter plots with contours showing co-expression of ANXA10 (x-axis) and IL-33 (y-axis) in 30 pixel square patches (roughly representing single lesions) in IMC data. A control K2 image (left) is compared to that of an sh*Il33* K2 image (right). sh*Il33* represses the spatial co-expression relationship between ANXA10 and IL-33. (B) Scatter plots with contours showing co-expression of VIM (x-axis) and IL-33 (y-axis) in 30 pixel square patches in IMC data. A control K2 image (left) is compared to that of a sh*Il33* K2 image (right). sh*Il33* does not substantially impact the spatial co-expression relationship between VIM (marking primarily stromal elements) and IL-33. (C) Milo neighborhoods overlaid on FDL built on scRNA-seq of K3 control and sh*Il33* cells. Each point represents one neighborhood scaled to size (number of cells) and is colored by either cell-state annotation (left) or Palantir pseudotime (right). (D) Milo neighborhoods overlaid on FDL in (C) colored by communication module assignment. Module expression per neighborhood is computed as the log of average normalized expression of each gene in that module; neighborhoods are assigned to the highest-expressed module.

